# Epithelial Ca^2+^ waves triggered by enteric neurons heal the gut

**DOI:** 10.1101/2023.08.14.553227

**Authors:** Afroditi Petsakou, Yifang Liu, Ying Liu, Aram Comjean, Yanhui Hu, Norbert Perrimon

**Affiliations:** Department of Genetics, Harvard Medical School, Boston, USA; Howard Hughes Medical Institute, Boston, USA

**Keywords:** *Drosophila*, gut, nerve-dependent regeneration, bioelectric signaling, enteric neuro-epithelial communication, cholinergic enteric neurons, nicotinic Acetylcholine Receptors, Acetylcholinesterase, gap junctions, innexins, Hippo signaling

## Abstract

A fundamental and unresolved question in regenerative biology is how tissues return to homeostasis after injury. Answering this question is essential for understanding the etiology of chronic disorders such as inflammatory bowel diseases and cancer. We used the *Drosophila* midgut to investigate this question and discovered that during regeneration a subpopulation of cholinergic enteric neurons triggers Ca^2+^ currents among enterocytes to promote return of the epithelium to homeostasis. Specifically, we found that down-regulation of the cholinergic enzyme Acetylcholinesterase in the epithelium enables acetylcholine from defined enteric neurons, referred as ARCENs, to activate nicotinic receptors in enterocytes found near ARCEN- innervations. This activation triggers high Ca^2+^ influx that spreads in the epithelium through Inx2/Inx7 gap junctions promoting enterocyte maturation followed by reduction of proliferation and inflammation. Disrupting this process causes chronic injury consisting of ion imbalance, Yki activation and increase of inflammatory cytokines together with hyperplasia, reminiscent of inflammatory bowel diseases. Altogether, we found that during gut regeneration the conserved cholinergic pathway facilitates epithelial Ca^2+^ waves that heal the intestinal epithelium. Our findings demonstrate nerve- and bioelectric-dependent intestinal regeneration which advance the current understanding of how a tissue returns to its homeostatic state after injury and could ultimately help existing therapeutics.

## Introduction

How a tissue returns to homeostasis after injury without undergoing inadequate or excessive repair is a fundamental question in regenerative biology. As long as this question remains unresolved the etiology and even cure of various disorders remains elusive. Several diverse and complex mechanisms have been proposed to influence repair (inflammation and regeneration), ranging from neuroimmune responses and activity of conserved signaling pathways to endogenous ion currents (bioelectric signaling) ^1–7^. However, it is unclear how these mechanisms return a previously injured tissue to its homeostatic state. Especially, for an organ like the intestine that regularly needs repair due to ingested irritants answering this question is pressing. Excessive repair in the intestine results in chronic intestinal pathologies such as inflammatory bowel diseases (IBDs, such as colitis) which are rising in the population remain without a cure and can lead to cancer ^3,8–13^.

The cholinergic pathway is an ancient conserved pathway^14^ used extensively by peripheral neurons to communicate with internal organs^15^. The two types of cholinergic receptors, nicotinic and muscarinic, and enzymes that modulate acetylcholine (ACh) metabolism, e.g. Acetylcholinesterase (AChE/Ace) are widely expressed in non-neuronal tissues^15^. Cholinergic receptors regulate ion transport in the intestinal epithelium and this is vital for water and nutrient absorption^16^. Recently, attention has been given to the anti-inflammatory properties of the cholinergic pathway, with reduced ACh responsiveness associated with intestinal diseases^5,16–18^. Furthermore, the nicotinic receptor (nAChR) agonist nicotine promotes growth and differentiation of mammalian intestinal organoids^19^. However, the regenerative properties of the cholinergic pathway in the gut are not well understood^20^.

The *Drosophila* midgut is equivalent to the mammalian small intestine and has been used extensively to identify the conserved molecular pathways that trigger inflammation and regeneration in the injured epithelium^21–23^. The midgut epithelium is single-layered and comprised of enterocytes (ECs), large polyploid epithelial cells specialized in absorption, secretory enteroendocrine cells (EEs) and progenitor cells (PCs)^21,24,25^. The visceral muscle and trachea surround the midgut epithelium whereas anterior and posterior midgut regions are innervated by enteric neurons^21^. When the epithelium is injured intestinal stem cells (ISCs) divide rapidly, giving rise to daughter cells (EBs) which differentiate into ECs and EEs^21–23^. Depending on the type of injury or infection, a multifaceted interplay of conserved inflammatory and regenerative pathways (e.g., EGFR, JAK-STAT, Wnt, BMP, Yki/Yap) activate ISC proliferation so that a sufficient PC (ISC/EB) pool is generated to replenish the damaged epithelium^23,26–38^. Despite the in-depth understanding of how repair is triggered in the midgut, it is unclear how the activities of these interlinked pathways are later dampened once the epithelium is ready to transition to homeostasis. Following specific damage such as infection by *Ecc15* or treatment with Bleomycin, the BMP signaling pathway has been reported to have dual roles first promoting proliferation and later promoting ISC-quiescence^4,7^. However, increase of BMP signals is injury-dependent^39^, suggesting that other mechanisms are also involved in ISC-quiescence.

Here, we provide evidence for the fundamental role of an epithelial bioelectric mechanism under the control of cholinergic enteric neurons that occurs as the midgut transitions from colitis-like injury to homeostasis, we refer to this transition as recovery. We show that ECs downregulate *Ace* and upregulate *nAChR subunit β3* (*nAChRβ3*) during recovery. Together, this makes ECs more sensitive and receptive to ACh that they receive from a specific population of cholinergic enteric neurons, which we named ARCENs for <u>A</u>nti-inflammatory <u>R</u>ecovery- regulating <u>C</u>holinergic <u>E</u>nteric <u>N</u>eurons. During recovery cholinergic signaling from ARCEN- innervations triggers elevated nAChRβ3-mediated Ca^2+^ influx in nearby ECs, that is then propagated across the epithelium via Innexin2/Innexin7 gap junctions. This advances EC maturation, reduces ISC proliferation and returns the gut to homeostasis. Inhibition of nAChRβ3-mediated Ca^2+^ currents during recovery cause ion imbalance and EC deficiency followed by increase of Egr/TNF and Yki/Yap, leading to excessive inflammatory and proliferative responses, including hyperplasia, that prevent return to homeostasis.

Altogether, we demonstrate that transition to homeostasis after injury relies on the healing functions of nAChR-mediated Ca^2+^ currents in the intestinal epithelium which are initiated locally by specific cholinergic enteric neurons and spread by gap junctions. Our study broadens the current understanding of how regeneration ends by showing *in vivo* how cooperation between peripheral neurons and epithelial bioelectric signaling directs a tissue towards homeostasis.

## Results

### Sensitivity of ECs to ACh is required for recovery

To study the *Drosophila* intestinal epithelium while it transitions to homeostasis after injury, we damaged the gut with dextran sodium sulfate (DSS). This induces colitis in mammals^40–42^ and has been used previously in *Drosophila* to identify conserved proliferative pathways^22,32^. We fed flies DSS for 4 days (injury period) followed by two or four days of standard food (recovery period, Fig. 1A). Damaging the gut with DSS elevated expression of the effector *Drosophila* caspase 1 (Dcp1), indicative of cell death (Fig. 1B), and of conserved inflammatory cytokines such as the IL6-like *Unpaired 3* (*Upd3*)^28,43^ and the TNF homolog *eiger (egr)*^44^ (Ext. Fig. 1A), resembling DSS-induced colitis in mammals^40–42^. Once flies were transferred to standard food, the epithelium required 4 days to return to homeostasis and be indistinguishable from an unchallenged gut as determined by the levels of i) inflammatory cytokines (Ext. Fig. 1A), ii) cell death (Fig. 1B), iii) proliferation (ISC mitotic divisions using the mitotic marker anti-pH3, Fig. 1C), and iv) the expression levels of PC marker *escargot* (*esg*; Ext. Fig. 1B) and of two markers of mature epithelial cells, *pdm1* a marker of ECs, and *prospero* (*pros*), a marker of EEs (Ext. Fig. 1B).

**Fig. 1.**
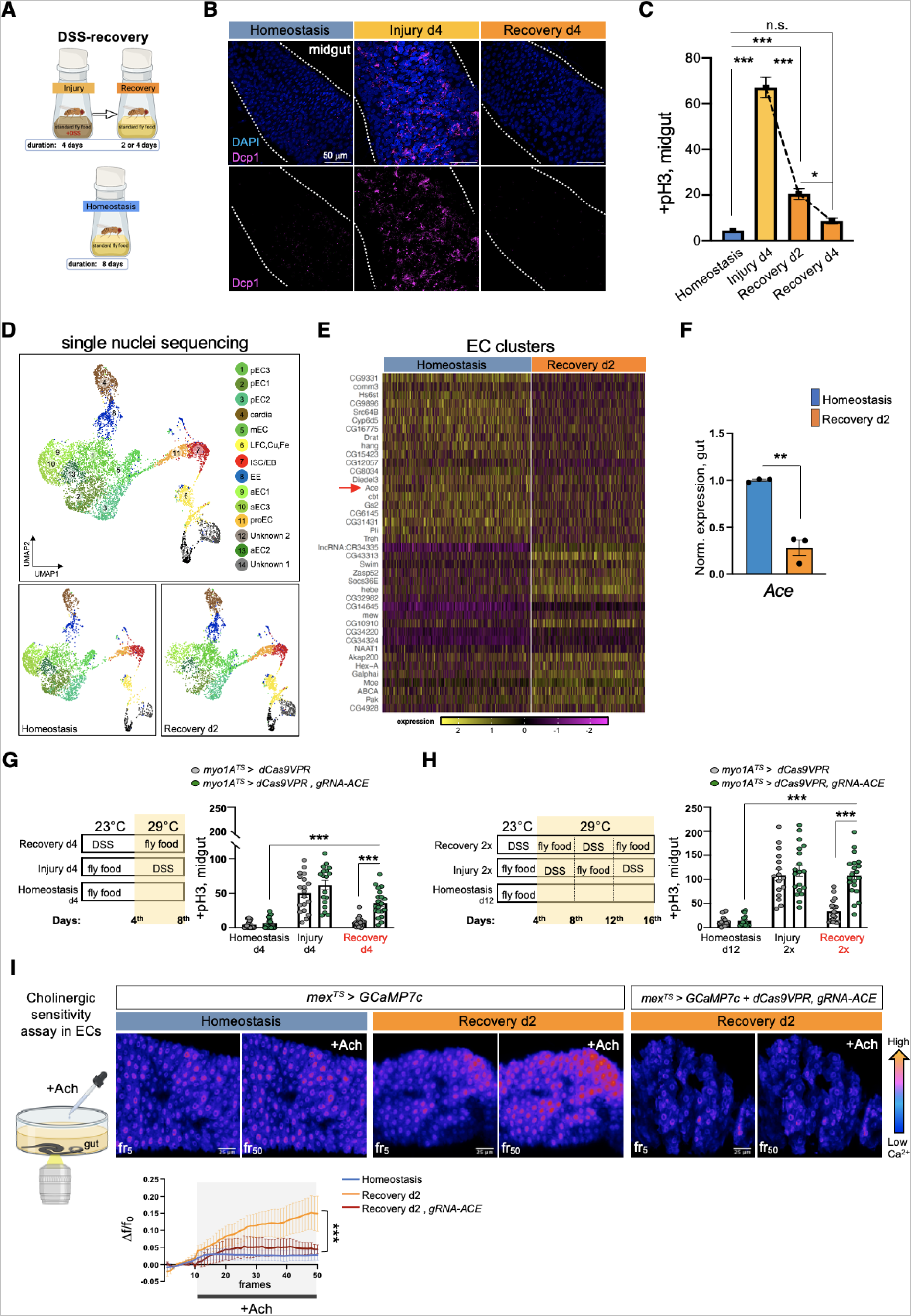
Sensitivity of ECs to ACh is required for recovery. **(A)** Experimental design for epithelial damaged guts in flies, consisting of Injury (4 days of DSS-feeding), Recovery (2 or 4 days after DSS feeding) and Homeostasis (unchallenged flies). We used DSS to damage the epithelium because it stimulates regeneration slower than bacterial infections (DSS: few days, infection: few hours)^32,128^, thus giving more time to experiment on pathways activated during and after injury. **(B)** Midgut confocal images stained with anti-Dcp1 (*Drosophila* caspase 1, magenta) and DAPI (nuclei, blue) during Homeostasis, Injury d4 and Recovery d4. Each condition was tested at 29°C and conducted as shown in 1a. **(C)** Mitotic division counts based on anti- PhoshoHistone-3 (pH3) staining of proliferating ISCs from midgut of control flies during Homeostasis (blue), Injury d4 (light orange), Recovery d2 and Recovery d4 (orange). Conditions are as described in 1B. N=29-32 guts per conditions. **(D)** Annotated gut cell type clusters visualized with UMAP plot of *Ore R* flies during Homeostasis and Recovery (d2) after DSS- injury. Two clusters that resembled previous unidentified clusters^45^ were assigned as unknown. Of the midgut clusters, seven clusters belonged to ECs, one to EEs and two clusters to PCs. The seven EC clusters are the anterior (aEC 1-3), middle (mEC) and posterior ECs (pEC 1-3) all of which highly express EC markers like *pdm1*, *myo1A* and *mex* (Ext. Fig. 1D) based on previous single-cell profiling^45^. The two PC clusters (ISC/EBs, proECs) are enriched for mesenchymal markers like *esg* and *Notch* (Ext. Fig. 1D). The ISC/EB cluster contains ISCs and EBs as determined by markers from previous single-cell profiling^45^ and the proECs cluster (progenitors of ECs) consist of EBs dedicated to EC differentiation based on markers like the elevated *Sox21a* expression^129^ (Ext. Fig. 1D). The EE cluster expresses EE markers *pros*, *piezo*, *AstA* (Ext. Fig. 1D). **(E)** Heatmap illustrating the most significantly upregulated and downregulated genes between Homeostasis and Recovery d2 in EC clusters (S. Table 1-2). *Acetylcholinesterase* (*Ace*, encodes the enzyme that breaks down ACh, red arrow) is the 14^th^ most significantly downregulated gene in ECs during Recovery. (**F)** Expression levels of *Ace* in *Ore R* guts during Homeostasis (blue) and Recovery d2 (orange), normalized to Homeostasis. N=3 biological samples per condition. **(G-H)** Experimental illustrations and accompanying graphs showing mitotic division counts (based on counts with anti-pH3 mitotic marker) from midgut of control flies (grey, *myo1A^TS^ > dCas9VPR*) and flies with *Ace* conditionally overexpressed in ECs using the EC driver *myo1A*-Gal4 (green, *myo1A^TS^ > dCas9VPR, gRNA- Ace*) in different conditions. All conditional perturbations (unless otherwise stated), refer to perturbations with the ubiquitous temperature-sensitive Gal4 inhibitor, *Tubulin-Gal80^TS^* (^TS^), that allows Gal4 expression only at warm temperature (>27°C). N=14-20 guts per genotype and condition. **(I)** Illustration of cholinergic sensitivity assay and color-coded sequential frames of midgut before (fr_5_) and after (fr_50_) ACh administration during Homeostasis and Recovery d2 of flies conditionally expressing the Ca^2+^ reporter GCAMP7c with the EC-driver *mex*-Gal4 (*mex^TS^ ^>^ GCaMP7c*) and flies overexpressing *Ace* (*mex^TS^ ^>^ GCaMP7c + dCas9VPR, gRNA-Ace*) in ECs (Movies S1-S3). For Recovery d2, flies were transferred for 2 days to standard food at 29°C after 4 days of DSS-feeding at 23°C. For Homeostasis, the experimental regime is similar to Recovery without DSS-feeding. The accompanying graph demonstrates the relative fluorescence intensity (ΔF/F_0_) per frame and per genotype (Homeostasis: blue line, Recovery d2: orange line, Recovery d2+*gRNA-Ace*: burgundy line). N=4-7 guts were used for each genotype per condition. All p-values were obtained with one-way Anova or two-way Anova or t-test; *: 0.05>p>0.01, ***:p<0.001, ns: non-significant. Mean ± s.e.m.

To search for recovery-specific differentially expressed genes, we performed snRNAseq on the 2^nd^ day of recovery (Fig. 1D). We identified 14 distinct clusters from the 8073 nuclei recovered (Fig. 1D), which we assigned to different epithelial and progenitor cell populations, as well as cells residing in the cardia and in the LFC/Cu/Fe gut region (Fig. 1D, Ext. Fig. 1C, Ext. Fig 1D), as determined by the expression of marker genes from a previous single-cell profiling study^45^. We next analyzed differential gene expression between homeostasis and recovery among epithelial and progenitor clusters. We observed that *Ace*, which encodes the cholinergic esterase Ace/AChE, is highly enriched in EC clusters and was significantly downregulated during recovery (Fig. 1E, Ext. Fig. 1E-1F, S. Table 1-2). We found ∼75% reduction of *Ace* during recovery in the gut (Fig. 1F). Supporting our results, previous intestinal RNA-seq profiling (Flygut-EPFL data)^46^ after different bacterial infections (*Ecc15* and *Pe*) detected *Ace* downregulation, indicating that this change in *Ace* levels occurs for different types of epithelial damage. Ace hydrolyzes ACh to choline and acetate, and thus defines a cell’s sensitivity to ACh. Notably, Ace is highly conserved^47^ and Ace/AChE inhibitors are tested as therapeutics for human intestinal pathologies^18,48–51^.

To test the role of *Ace* during regeneration, we used CRISPR/Cas9 transcriptional activation^52^ to conditionally overexpress *Ace* (Ext. Fig. 1G). Strikingly, 4 days of *Ace* overactivation in ECs (using the Gal4^53^ *myo1A-*driver together with the repressor *Tubulin*-Gal80^TS^ ^54^, referred as *myo1A^TS^*) led to excessive ISC proliferation during recovery (Fig. 1G), while the same conditional activation during homeostasis or injury did not affect ISC proliferation (Fig. 1G). Similarly, consecutive DSS challenges, while *Ace* is conditionally overexpressed in ECs using the *myo1A^TS^*or *mex^TS^* driver (another EC driver), led to recovery specific ISC over-proliferation (Fig. 1H, Ext. Fig. 1H). We next tested if *Ace* perturbations in visceral muscle or immune cells could regulate ISC proliferation. Conditionally overexpressing *Ace* in visceral muscle and immune cells did not alter ISC proliferation when compared to control guts during homeostasis, injury, and recovery (Ext. Fig. 1I). Altogether, these findings reveal that overexpressing *Ace* in ECs during recovery prevents ISCs from becoming quiescent, causing an excessive regenerative response.

We also tested if this effect is related to the type of epithelial damage. To do this, we used *Ecc15* oral infection, which has been reported to trigger different pathways from DSS^39^. Conditionally overexpressing *Ace* in ECs for 18hrs after *Ecc15* infection also led to significant over-proliferation (Ext. Fig. 1J).

ACh has been proposed to modulate ion transport in the intestinal epithelium in a Ca^2+^- dependent manner^16,55^. Thus, to test for epithelial changes in ACh sensitivity during homeostasis and recovery, we visualized Ca^2+^ by conditionally expressing the Ca^2+^ indicator GCAMP*7c* in ECs. Strikingly, using *ex vivo* live gut imaging, we find that Ca^2+^ levels in ECs are significantly higher during recovery following ACh administration than during homeostasis, and that this increase in ACh sensitivity is attenuated upon overexpression of *Ace* in ECs (Fig. 1I and Movies S1-3). These results indicate that during recovery ECs become more sensitive to ACh by decreasing *Ace* and that this change in ACh sensitivity is required for the intestinal epithelium to transition to homeostasis after injury.

### nAChR***β***3 is required in ECs for recovery

Cholinergic receptors are highly conserved and are either G-protein-coupled muscarinic receptors (mAChR) or the ligand-gated ion channel nicotinic receptors (nAChR) made of five homomeric or heteromeric subunits (α1, α2, α3, α4, α5, α6, α7, β1, β2 or β3). To identify which cholinergic receptor in ECs becomes activated once sensitivity to ACh is increased during recovery, we screened all nAChR subunits and different mAChR subtypes by knocking down their expression in ECs. Interestingly, conditional RNAi expression for 4 days targeting *nAChRβ3* in ECs caused over-proliferation specifically during recovery (Fig. 2A, Fig. 2B, Ext. Fig. 2A, Ext. Fig. 2B) and knockdown combined with repeated DSS injury (Recovery 2x) led to hyperplasia (Fig. 2B, Fig. 2C). Conditional knockdown of *nAChRβ3* in ECs also significantly reduced the Ca^2+^ response after ACh administration during recovery (Ext. Fig. 2C). Taken together, these data suggest that reducing *nAChRβ3* in ECs leads to similar phenotypes as observed following *Ace* upregulation. Importantly, this effect is specific to ECs, as conditionally knocking down *nAChRβ3* in PCs, EBs alone, EEs, visceral muscle and hemocytes had no effect on proliferation (Ext. Fig. 2D).

**Fig. 2.**
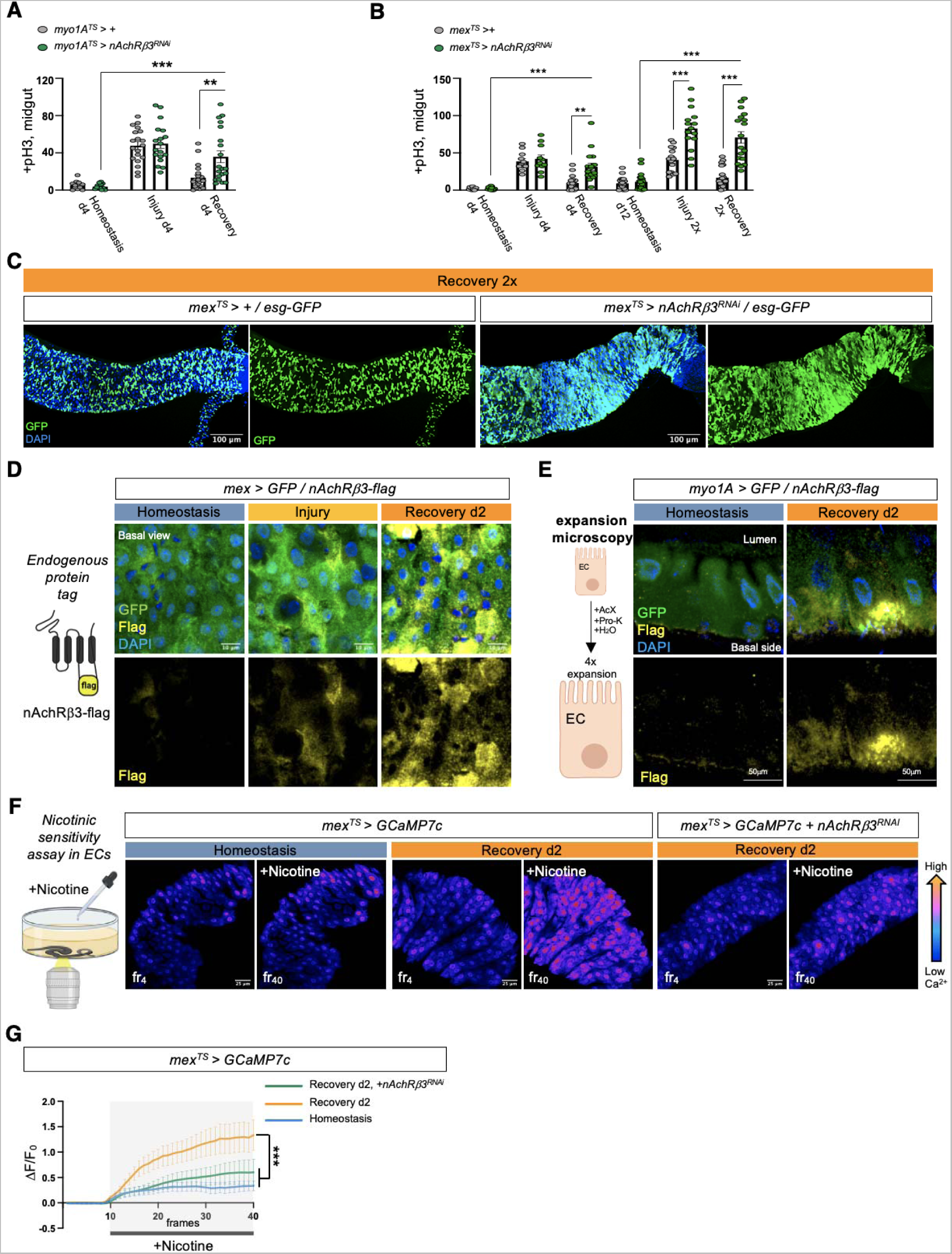
nAChRβ3 is required in ECs for recovery. **(A)** Mitotic division counts from midgut of control flies (*myo1A^TS^ > +,* grey) and flies with *nAChR subtype β3* conditionally reduced in ECs (*myo1A^TS^ > nAChRβ3^RNAi^*, green) during Homeostasis d4, Injury d4 and Recovery d4 (like Fig. 1G). N=12-20 guts per condition and genotype. **(B)** Mitotic division counts from midgut of control flies (*mex^TS^ > +,* grey) and flies with *nAChR subtype β3* conditionally reduced in ECs (*mex^TS^ > nAChRβ3^RNAi^*, green) during Homeostasis d4, Injury d4, Recovery d4, Homeostasis d16, Injury 2x, Recovery 2x (like Fig. 1G-H). N=11-20 guts per condition and genotype. **(C)** Confocal gut images of *mex^TS^> +* and *mex^TS^> nAChRβ3^RNAi^*flies co-expressing the PC marker esg-GFP (green, anti-GFP) during Recovery 2x (like Fig 2B). **(D)** Illustration of the endogenous tag for nAChRβ3 (nAChRβ3-flag) that was generated for this study and confocal images of midgut from flies expressing nAChRβ3-flag and GFP-expressing ECs (*mex >GFP*) during Homeostasis, Injury and Recovery d2. nAChRβ3-flag was visualized with anti-Flag (yellow) staining and ECs with anti-GFP staining (green). **(E)** Expansion microscopy schematic followed by confocal images of 4x expanded midguts from flies expressing nAChRβ3-flag (yellow) and GFP-expressing ECs (*myo1A >GFP,* green) during Homeostasis and Recovery d2. **(F-G)** Illustration of nicotinic sensitivity assay and color-coded sequential frames of midgut before (fr_4_) and after (fr_40_) nicotine administration during Homeostasis and Recovery d2 of *mex^TS^ > GCaMP7c* flies and *mex^TS^ ^>^ GCaMP7c+nAChRβ3^RNAi^* flies (Movies S4-S6). Bottom graph shows the relative fluorescence intensity (ΔF/F_0_) per frame per condition and genotype (Homeostasis: blue line, Recovery d2: orange line, Recovery d2+*nAChRβ3^RNAi^*: green line). Experimental conditions were like Fig. 1I. N=8-13 guts per condition and genotype. DAPI: nuclei. All p-values were obtained with one-way Anova or two-way Anova or t-test; *: 0.05>p>0.01, **: 0.01<p<0.001, ***: p<0.001. Mean ± s.e.m.

The profiling depth of our single nuclei sequencing was not sufficient to conclude if *nAChRβ3* expression is altered between homeostasis and recovery, despite being solely found in ECs (Ext. Fig. 2E). To visualize the expression of nAChRβ3, we used scarless CRISPR gene editing (https://flycrispr.org/scarless-gene-editing/)56 to insert a Flag tag into the intracellular loop region between the M3 and M4 transmembrane domains of nAChRβ3^57^ (nAChRβ3-flag) (Fig. 2D, Ext. Fig. 2F). Strikingly, we found that endogenous nAChRβ3 was significantly enriched in ECs by day 2 of recovery (Fig. 2D), whereas a decrease in *nAChRβ3* levels coincided with the return to homeostasis (Ext. Fig. 2G). Moreover, we found that nAChRβ3 was clearly localized to the basal side of ECs using expansion microscopy^58^ to uniformly enlarge the epithelium and increase imaging resolution (Fig. 2E). Interestingly, some ECs had more nAChRβ3 clustered on the basal side than others (Fig. 2D, Fig. 2E).

To further verify that nAChRs are enriched in ECs during recovery, we modified the cholinergic sensitivity assay by adding the cholinergic agonist nicotine, which activates only nAChRs and cannot be hydrolyzed by Ace (Fig. 2F). Nicotine administration significantly increased Ca^2+^ in ECs during recovery compared to homeostasis, reminiscent of ACh- sensitivity, and this increase was diminished when *nAChRβ3* was knocked down (Fig. 2F, Fig. 2G and Movies S4-S6). Overall, we conclude that *nAChRβ3* in ECs is essential for gut recovery and recovery-specific enrichment of nAChRβ3 provides an additional level of regulation that likely ensures that ECs are more responsive to ACh while the epithelium transitions to homeostasis.

### nAChR**β**3-mediated Ca^2+^ increase in ECs promotes recovery

ISC proliferation is triggered by the release of different cytokines which vary depending on the stimulus^22^. To identify the signaling pathways responsible for unrestrained proliferation during recovery after *nAChRβ3* knockdown in ECs, we tested the expression of known cytokines (Fig. 3A). The IL6-like *unpaired 2* (*upd2*) and *upd3* JAK-STAT ligands, together with the EGF-like ligand *vein* (*vn*) and TNF-homolog *egr*, were significantly upregulated in ECs with *nAChRβ3* knockdown during recovery (Fig. 3A). Upd2, Upd3 and Vn are released on activation of the Hippo pathway effector Yorkie (Yki/YAP) in damaged ECs^30,32^, whereas Egr is associated with cell death^59^. Supporting this, we found that knocking down *nAChRβ3* in ECs during recovery significantly increased cell death as well as the expression of the Yki target *Diap1* (Ext. Fig. 3A, Ext. Fig. 3B). Importantly, knockdown of *nAChRβ3* significantly reduced the transcript and protein levels of *pdm1* marker for mature ECs specifically during recovery (Fig. 3B, Ext. Fig. 3C), while the EE marker *pros* remained unchanged (Ext. Fig. 3D). Together, these data suggest that disruption of nAChRs in ECs during recovery impairs ECs causing Yki-signaling activation, cell death and the subsequent release of inflammatory signals that induce unwarranted ISC proliferation.

**Fig. 3.**
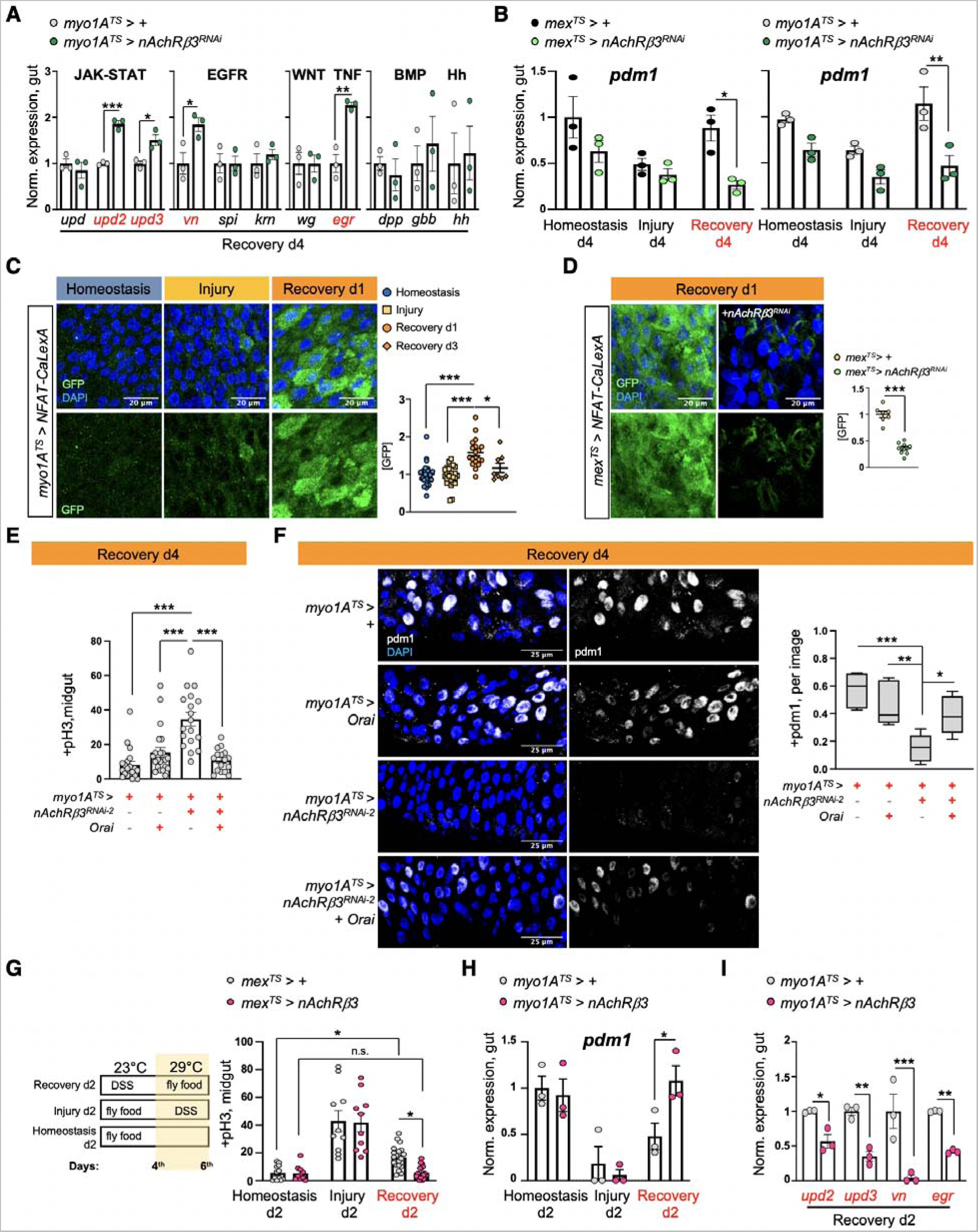
nAChRβ3-mediated Ca^2+^ increase in ECs promotes recovery. **(A)** Expression levels of proliferating cytokines during recovery in control guts (*myo1A^TS^ > +,* grey) and guts with *nAChRβ3* knocked down in ECs (*myo1A^TS^ > nAChRβ3^RNAi^*, green). Conditions are like in Fig. 1G. JAK-STAT ligands: *Upd* (*Unpaired*), *Upd2 (Unpaired 2*), *Upd3*. EGFR ligands: *vn* (*vein*), *spi* (*spitz*), *krn* (*keren*). Wnt ligand: *wg* (*wingless*). TNF ligand: *egr*. BMP ligands: *dpp* (*decapentaplegic*), *gbb* (*glass bottom boat*). Hh ligand: *hh* (*hedgehog*). N=3 biological samples per genotype. **(B)** Expression levels of the EC marker *pdm1* in control flies (*mex^TS^>+*, black and *myo1A^TS^>+*, grey) and flies with *nAChRβ3* knock down in ECs (*mex^TS^> nAChRβ3^RNAi^*, light green and *myo1A^TS^> nAChRβ3^RNAi^* dark green). Conditions are like in Fig. 1G. N=3 biological samples per genotype and per condition. **(C)** Posterior midgut images with the Ca^2+^ activity transcriptional reporter NFAT-CaLexA conditionally expressed in ECs (*myo1A^TS^ >NFAT- CaLexA*) during Homeostasis, Injury and Recovery d1. The reporter was expressed for 2 days at 29°C in each condition. ECs with elevated endogenous Ca^2+^ (high GFP, bright green) were visualized with anti-GFP. The accompanying graph shows the fold change of GFP fluorescence per image and condition. Homeostasis: blue circle, Injury: light orange square, Recovery d1: orange circle, Recovery d3: diamond. N=25-8 guts per condition. **(D)** Posterior midgut images with the NFAT-CaLexA reporter in control guts during Recovery d1 (*mex^TS^ > NFAT-CaLexA)* and when *nAChRβ3* is conditionally reduced in ECs (*mex^TS^ >NFAT-CaLexA+nAChRβ3^RNAi^*). Flies were fed DSS-food for 4 days at 23°C and then transferred to standard food at 29°C for 24hrs. N=7-9 guts per genotype. **(E)** Mitotic division counts of midguts from control flies (*myo1A^TS^>+*), flies conditionally overexpressing the Ca^2+^ channel Orai in ECs (*myo1A^TS^>Orai*), flies conditionally reducing *nAChRβ3* in ECs (*myo1A^TS^>nAChRβ3^RNAi^*) and flies conditionally knocking down *nAChRβ3* while overexpressing *Orai* in ECs (*myo1A^TS^>nAChRβ3^RNAi^ + Orai*). Conditions are like in Fig 1G. N=16-22 guts per genotype. **(F)** Confocal images of posterior midgut from similar flies and condition as in Fig. 3E stained with the EC marker anti-pdm1 (grey). The following boxplot indicates the ratio of pdm1+ nuclei versus all nuclei (DAPI+) per image and genotype. N=7 guts per genotype. **(G)** Experimental conditions schematic and graph with mitotic division counts of midguts from control flies (*mex^TS^>+,* silver) and flies with *nAChRβ3* overexpressed in ECs (*mex^TS^> nAChRβ3,* pink) during Homeostasis d2, Injury d2 and Recovery d2. N=10-21 guts per genotype and condition. **(H-I).** Expression levels of the EC marker *pdm1* and of inflammatory cytokines *upd2, upd3, vn, egr* from control flies (*myo1A^TS^>+,* silver) and flies with *nAChRβ3* overexpressed in ECs (*myo1A^TS^> nAChRβ3,* pink). Conditions like Fig.3G. N=3 biological samples per genotype. DAPI: nuclei. All p-values were obtained with one-way Anova or two-way Anova or t-test; *: 0.05>p>0.01, **: 0.01<p<0.001, ***: p<0.001. n.s.: non-significant. Mean + s.e.m.

Cholinergic receptors regulate ion transport in the mammalian epithelium^16^. We asked whether nAChRs have similar functions in ECs using dyes that detect Na^+^ (SodiumGreen) or Cl^-^ (MQAE), as well as the Ca^2+^ transcriptional reporter NFAT-CalexA^60^. We found that reduction of *nAChRβ3* in ECs during recovery caused significant ion imbalance in the epithelium, with reduced Cl^-^ and Na^+^ levels (Ext. Fig. 3E). Strikingly, we observed that Ca^2+^ is significantly upregulated for a short period of time early in recovery before returning to levels resembling homeostasis (Fig. 3C). This endogenous Ca^2+^ increase disappears when *nAChRβ3* expression is knocked down in ECs (Fig. 3D). In addition, Ca^2+^ increase occurs only in ECs during recovery, as ISCs that use Ca^2+^ for proliferation ^61,62^ show Ca^2+^ decline during recovery (Ext. Fig. 3F). To examine the importance of nAChR-mediated Ca^2+^ during recovery, we genetically compensated for Ca^2+^ in *nAChRβ3-*deficient ECs. We found that conditional overexpression of the Ca^2+^ channel Orai combined with knockdown of *nAChRβ3* in ECs was sufficient to restore i) ISC proliferation (Fig. 3E), ii) pdm1+ ECs (Fig. 3F) and iii) Cl^-^ (Ext. Fig. 3G) to levels identical to controls.

We also conditionally overexpressed *nAChRβ3* in ECs (Ext. Fig. 3H), which doubles the amount of Ca^2+^ influx in ECs after nicotine administration (Ext. Fig. 3I). We found that *nAChRβ3* overexpression in ECs significantly expedited recovery of the intestinal epithelium with ISC proliferation and *pdm1* expression reaching levels indistinguishable from unchallenged guts in 2 days, half the normal time (Fig. 3G, Fig. 3H). Overexpressing *nAChRβ3* in ECs also significantly reduced inflammatory cytokine levels during recovery (Fig. 3I). Notably, *nAChRβ3* overexpression in ECs during homeostasis and injury did not change overall proliferation levels (Fig. 3G). Taken together, our data show that nAChR-mediated Ca^2+^ influx in ECs controls intestinal epithelium recovery by promoting EC maturation and ion balance. Disruption of nAChR-mediated Ca^2+^ influx causes EC deficiency and ion imbalance which culminates in over-inflammation and over-proliferation reminiscent of lasting unresolved injury. In contrast, overexpression of *nAChRβ3* expedites the return of the epithelium to its homeostatic state.

### Neuro-EC interactions promote nAChR-mediated gut recovery

ACh is released by neuronal and non-neuronal cells that express *ChAT* (*Choline acetyltransferase*), the enzyme that catalyzes the synthesis of ACh^15^. To identify the source of endogenous ACh responsible for promoting nAChR-mediated gut recovery, we first tested midgut cells (PCs, ECs, EEs), the visceral muscle and immune cells (hemocytes). ISC proliferation during recovery remained unaffected when *ChAT* was conditionally knocked down in these cells (Ext. Fig. 4A, Ext. Fig. 4B). Similarly, EE-less guts^63^ do not over-proliferate during recovery (Ext. Fig. 4C). Altogether, these data point to a neuronal source of ACh. The importance of neurons during regeneration has been reported in various contexts from limb regeneration to heart injury^64–66^. For the gut, recent studies have highlighted the anti-inflammatory properties of mammalian enteric (gut-surrounding) neurons^17,67–69^ while limited associations have been made between neurons and ISC proliferation in *Drosophila*^70,71^.

Since ACh is a short-distance neurotransmitter/local neurohormone^14^, the most likely neuronal source of ACh during recovery is the enteric nervous system. However, an in-depth characterization of cholinergic enteric neurons in *Drosophila* has been lacking. *Drosophila* enteric neurons have the unique feature of innervating the gut even though their cell bodies reside in the brain, the hypocerebral ganglion (HCG), or the thoracicoabdominal ganglia (TAG or adult ventral nerve cord)^21,72^. Combined with the coiling structure of the gut, this feature makes descending enteric innervations vulnerable to damage during dissection and difficult to visualize, especially in the abdomen, in which most of the midgut resides^21,46^. To keep both cholinergic enteric innervations and the midgut intact we fixed whole flies and sectioned them with a vibratome (Ext. Fig. 4D). Cholinergic enteric innervations, which we marked with GFP using a *ChAT*-driver, are found in anterior (R1, R2) and posterior midgut regions (R4, R5) and originate from brain, TAG and HCG enteric neurons (Ext. Fig. 4D, Ext. Fig. 4E, Ext. Fig. 4F). Cholinergic enteric innervations extensively arborize, although the overall gut coverage is not vast (Ext. Fig. 4D, Ext. Fig. 4E), and remained in place after injury (Ext. Fig. 4E). We found that cholinergic innervations run between the muscle and the epithelium in both the anterior and posterior midgut regions (Ext. Fig. 4G, Ext. Fig. 4H) consistent with a previous report on enteric innervations^70^. Thus, cholinergic enteric innervations are adjacent not only to muscle cells but also to ECs (Ext. Fig. 4H).

Next, we used the Syt1 antibody to detect the synaptic vesicle membrane protein Synaptotagmin1, which is essential for neurotransmitter release^73^. Our data revealed the presence of several Syt1^+^ swellings along cholinergic enteric arborizations that innervate the muscle and the epithelium during recovery (Ext. Fig. 4I). This indicates that enteric innervations release ACh in the vicinity of both muscle and epithelium, likely interacting with both. We refer to these Syt1^+^ swellings as presynaptic boutons because they resemble *en passant* varicosities described in the autonomous nervous system, and are where a neurotransmitter diffuses to receptors located in the nearby tissue^74^. Interestingly, during homeostasis we observed that cholinergic enteric innervations carry less Syt1^+^ boutons (Ext. Fig 4J), potentially indicating less ACh release during homeostasis. Taken together, our data show that cholinergic enteric innervations are in the right position to release ACh to ECs during recovery.

The majority of cholinergic midgut innervation originate from the TAG region, as determined after co-expressing the TAG Gal80 inhibitor (Tsh-Gal80)^75^, which repressed GFP in almost all abdominal cholinergic innervations (Ext. Fig 4K). To find a driver specific to cholinergic enteric neurons, we screened through different neuronal drivers^76^ expressed strongly in TAG. We identified the *R49E06-Gal4* driver, which is strongly expressed in the abdominal ganglion of the TAG, has no expression in the gut and very limited expression in the brain (Ext. Fig. 4L&FlyLight). We found that in the abdominal ganglion about 20 of the ∼35 neurons are ChAT+ and are therefore cholinergic (Ext. Fig. 4L and Fig. 4A). Notably, descending projections from R49E06*-*neurons are cholinergic (ChAT+) (Fig. 4A) and innervate the midgut in R2, R4 and R5 (Fig. 4B, Fig. 4C and Ext. Fig. 5A), suggesting that this driver targets a subpopulation of cholinergic enteric neurons. To have more genetic flexibility to study how these neurons interact with ECs we used the LexA binary system^77, 77,78^ to generate an EC- LexA driver (*mex-LexA::GAD* together with *Tubulin*-Gal80^TS^, referred to as *mexLexA^TS^*) (Ext. Fig. 5B). We used this driver to test if R49E06-innervations are in close proximity to ECs. Specifically, we conditionally expressed GFP in ECs with the *mex-LexA^TS^* driver and used the Gal4/UAS^53^ system to express the synaptic-vesicle marker Syt1HA in R49E06-neurons. We observed that during recovery Syt1HA-innervations from R49E06-neurons are in close proximity to muscle cells and ECs (Fig. 4D, Ext. Fig. 5C, Ext. Fig. 5D and Movie S7), suggesting that R49E06-innervations could be the source of ACh during gut regeneration.

**Fig. 4.**
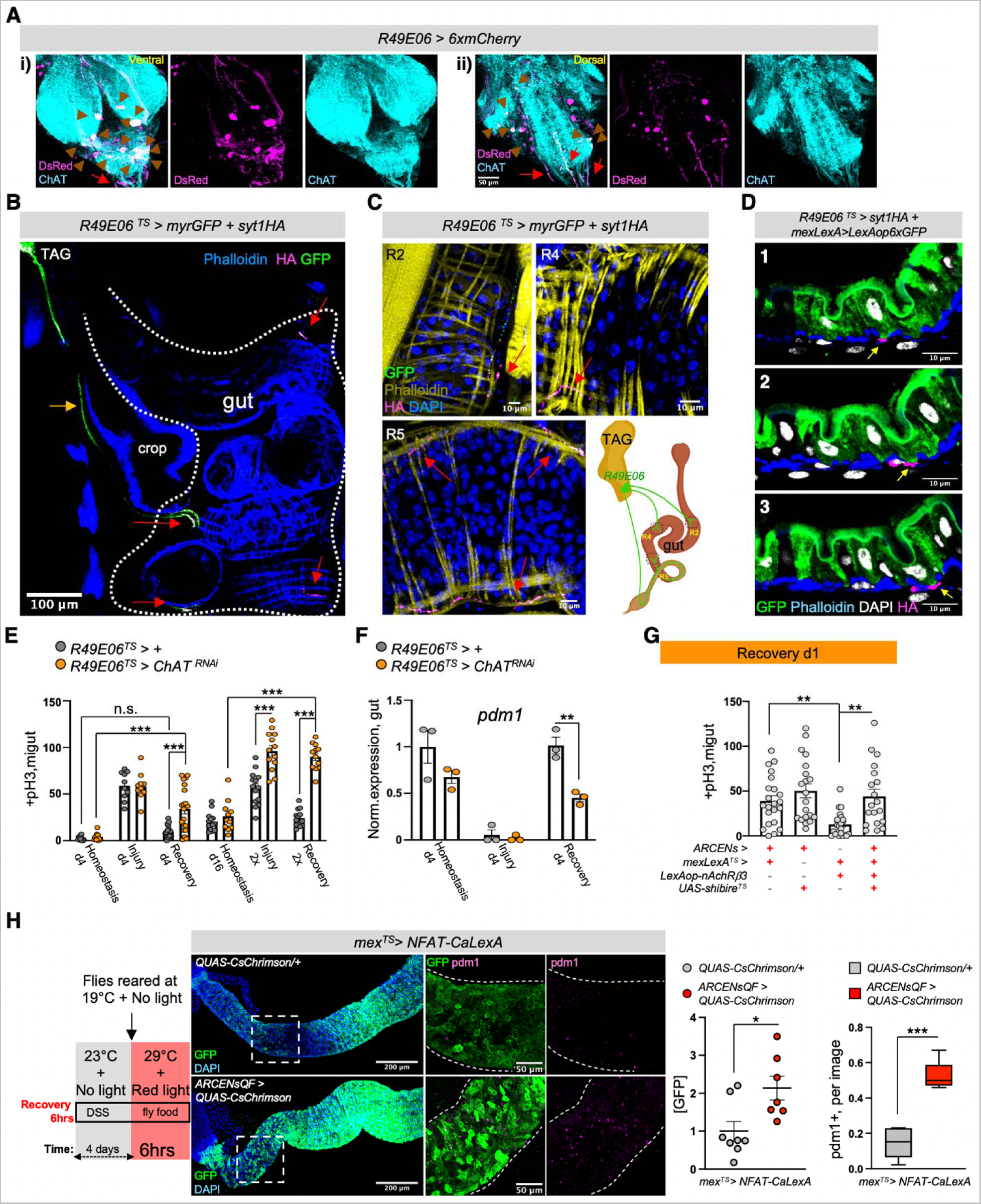
Neuro-EC interactions promote nAChR-mediated gut recovery. **(A)** Confocal image of the ventral (i) and dorsal (ii) side of the abdominal ganglion from *R49E06>6xmCherry* fly stained with anti-DsRed (magenta) and the cholinergic marker anti-ChAT (cyan). The confocal stack of one projection image is split in two (ventral, dorsal) for better visualization. Brown arrowheads: ChAT+ R49E06*-*neurons. Red arrows: ChAT+ R49E06*-*descending projections. **(B)** Confocal image after vibratome-sectioning (as in Ext. Fig. 4D) of the abdominal midgut (white dots) in *R49E06^TS^>myrGFP+syt1HA* flies. R49E06-enteric innervations were visualized with anti-GFP (green) and the presynaptic marker syt1HA was visualized with anti-HA (magenta). Orange arrow: descending projection. Red arrow: R49E06-innervations expressing syt1HA. Fly was sectioned during Recovery d2 (4 days DSS-food at 23°C and then transferred to standard food for ∼48hrs at 29°C). Phalloidin: muscle (blue). **(C)** Confocal images of R2, R4 and R5 innervated areas by R49E06-neurons after vibratome-sectioning from *R49E06^TS^> myrGFP+syt1HA* flies during Recovery d2 (like Fig. 4B). Red arrow indicates R49E06- innervations (anti-GFP, green) expressing syt1HA (anti-HA, magenta). Phalloidin: muscle (yellow). DAPI: nuclei (blue). Following illustration shows midgut regions innervated by R49E06 based on vibratome sections. **(D)** Sequential confocal images (1-3) from R49E06-innervated R4 region (Movie S7) after vibratome-sectioning of R49E06*^TS^>syt1HA+mexLexA>6xLexAopGFP* during Recovery d2 (like Fig 4B). Yellow arrows show R49E06-innervations with the synaptic- vesicle protein syt1HA (anti-HA, magenta) in close proximity to ECs (anti-GFP, green). Phalloidin: muscle (blue). DAPI: nuclei (white). **(E)** Mitotic counts from control flies (*R49E06^TS^>+,* grey) and flies with *ChAT* conditionally knocked down in R49E06*-*expressing neurons (*R49E06^TS^>ChAT^RNAi^,* orange). Conditions are like in Fig. 1G-H. N=10-20 guts per conditions per genotype. (**F)** Expression levels of the EC marker *pdm1* from guts of *R49E06^TS^>+* (grey) and *R49E06^TS^>ChAT^RNAi^* (orange) flies. Conditions are like in Fig. 1G. N=3 biological samples per genotype and per condition. Levels normalized to control guts during Homeostasis. From now on *R49E06-*expressing neurons will be referred as ARCENs: <u>A</u>nti- inflammatory <u>R</u>ecovery-regulating <u>C</u>holinergic <u>E</u>nteric <u>N</u>euron<u>s</u>. **(G)** Midgut mitotic division counts of flies with conditional overexpression of *nAcRβ3* in ECs (*mexLexA^TS^ > lexAop- nAcRβ3*), conditional thermo-silencing (>27°C) of ARCENs with *shibire^TS^* (*ARCENs>shibire^TS^*) and flies with both (*ARCENs^TS^>shibire^TS^+ mexLexA^TS^>lexAop-nAcRβ3*) during Recovery d1. Control: *ARCENs > + / mexLexA^TS^ > +*. Flies were fed for 4 days DSS-food at 23°C and then transferred to standard food for ∼24hrs at 29°C. N=19-24 guts per genotype. **(H)** Schematic of experimental conditions describing thermo- and opto-genetic inductions for 6hrs during recovery followed by images of posterior midgut from flies expressing the endogenous Ca^2+^ reporter in ECs (*mex^TS^>NFAT-CaLexA / QUAS-CsChrimson*) and flies that concurrently have ARCENs depolarized with the CsChrimson red light-gated cation channel (*mex^TS^> NFAT-CaLexA / R49E06QF > QUAS-CsChrimson*). ECs with elevated levels of endogenous Ca^2+^ are visualized with anti-GFP (green) and mature ECs are visualized with anti-pdm1 (magenta). DAPI: nuclei (blue). The following graphs show the GFP fold change and the ratio of pdm1+ nuclei. N= 7-8 guts. All p-values were obtained with one-way Anova or two-way Anova or t-test; *: 0.05>p>0.01, **: 0.01<p<0.001, ***: p<0.001. n.s.: non-significant. Mean + s.e.m.

To test whether enteric R49E06-neurons regulate gut regeneration, we conditionally reduced *ChAT* in these neurons in conditions of gut homeostasis, injury, and recovery. This caused: i) ISC over-proliferation after 4 days of recovery, which intensified after repetitive injury (Fig. 4E), ii) recovery-specific reduction of ECs (Fig. 4F), iii) no significant change in EEs (Ext. Fig. 5E) and iv) increase of gut inflammatory cytokines during recovery (Ext. Fig. 5F). Therefore, reduction of ACh synthesis from R49E06-neurons during recovery leads to lasting unresolved injury, resembling *nAChRβ3* and *Ace* deregulation in ECs. These data show that intestinal repair after injury is under neuronal control, so we named these neurons <u>A</u>nti-inflammatory <u>R</u>ecovery-regulating <u>C</u>holinergic <u>E</u>nteric <u>N</u>eurons (ARCENs).

To test whether ARCENs are required for nAChR-mediated advance in recovery, we blocked neurotransmitter release with the *UAS-shibire^TS^* transgene^79^ while simultaneously conditionally overexpressing *nAChRβ3* in ECs (Fig. 4G, Ext. Fig 5B). We found that 24hr expression of *shibire^TS^* in ARCENs was sufficient to prevent *nAChRβ3* overexpression in ECs from fast-reducing ISC proliferation (Fig. 4G). To verify that ARCENs release ACh to the intestinal epithelium, we overexpressed the ion channel *TrpA1*^80^ and thermo-genetically depolarized ARCENs for 6 hrs (Ext. Fig. 5G). TrpA1-induction of ARCENs during the first 6 hours of recovery significantly reduced ISC proliferation, whereas additionally expressing the cholinergic repressor *ChAT*-Gal80 restored proliferation to levels identical to controls (Ext. Fig. 5G). In addition, TrpA1-induction of ARCENs significantly reduced the expression of gut inflammatory cytokines during the first 6 hours of recovery (Ext. Fig. 5H). Moreover, we used the QF system^78^ to generate a R49E06-QF (ARCEN-QF) driver (Ext. Fig. 5B). We used this QF driver together with the light-gated cation channel CsChrimson^81^ to depolarize ARCENs while conditionally expressing in ECs the Ca^2+^ transcriptional reporter *NFAT-CaLexA*^60^ (Fig. 4H). Opto-genetic activation of ARCENs during the first 6 hours of recovery was sufficient to significantly increase both endogenous Ca^2+^ and the levels of pdm1+ ECs (Fig. 4H). Strikingly, ARCEN activation led to a broad Ca^2+^ spread across ECs (Fig. 4H) despite the limited innervation of ECs. Altogether, these data support that ARCENs provide ACh to ECs during gut regeneration.

Several studies have provided evidence for a protective role for peripheral neurons during inflammation^67–69,82,83^, with the cholinergic anti-inflammatory reflex proposed to sense inflammatory signals like TNF and reduce them^2,18,84^. *Drosophila* peripheral neurons have been reported to respond to TNF/Egr signaling through *wengen* (*wgn*)^85^, one of the two known TNF receptors (*wgn* and *Grindelwald*)^86,87^. To test whether the protective role of ARCENs during gut regeneration is linked to inflammatory signals like TNF/Egr, we conditionally knocked down *wgn* and *grnd* in ARCENs (Ext. Fig. 5I). Reduction of *wgn* and not *grnd* led to significant ISC over-proliferation during recovery (Ext. Fig. 5I). Importantly, knocking down *wgn* in ARCENs did not cause over-proliferation during homeostasis or injury (Ext. Fig. 5I), resembling the phenotypes following reduction of *ACh* in ARCENs (Fig. 4E) and reduction of *nAnchRβ3* in ECs (Fig. 2A-B) . These data support that TNF signaling via Wgn regulates the regenerative roles of ARCENs, reminiscent of the anti-inflammatory reflex response.

Taken together, our data support that ARCENs release ACh to ECs during recovery and that ARCEN-EC interactions are required to promote transition to homeostasis after injury by activating nAChR-mediated Ca^2+^ influx in ECs, increasing mature ECs, reducing proliferation, and decreasing inflammation in the intestinal epithelium. Our findings demonstrate nerve-dependent intestinal regeneration, placing the intestinal epithelium among the many tissues whose ability to regenerate depends on neurons^64^.

### nAChR-mediated Ca^2+^ propagates via inx2/inx7 gap junctions

Gap junctions propagate signals like ions and small metabolites between cells. We asked whether gap junctions are involved in the spreading of Ca^2+^ among ECs during recovery by targeting innexins which form gap junctions in invertebrates^88–90^. Single nuclei profiling indicated that *inx2* and *inx7* are similarly enriched in ECs whereas their expression in PC and EE clusters is lower (Ext. Fig. 6A, Ext. Fig 6B, Ext. Fig. 6C). We found that knocking down of either *inx2* or *inx7* in ECs for 24 hrs during recovery weakened Ca^2+^ and pdm1+ levels, resembling the effects of *AChRβ3* knockdown (Fig. 5A). Further, conditionally reducing *inx2* in ECs for 4 days caused recovery-specific over-proliferation, whereas no significant changes occurred during homeostasis or injury (Fig. 5B). Strikingly, *inx7* reduction in ECs disrupted Inx2 gap junction formation (Ext. Fig. 6D, Ext. Fig. 6E), suggesting that Inx2 and Inx7 form heteromeric gap junctions among ECs. Gap junctions are activated by membrane potential changes, including opening of gap junctions after increase in intracellular Ca^2+^ levels ^88,91,92^. Together, these data support the model that Inx2/Inx7 gap junctions between ECs are activated during recovery.

**Fig. 5.**
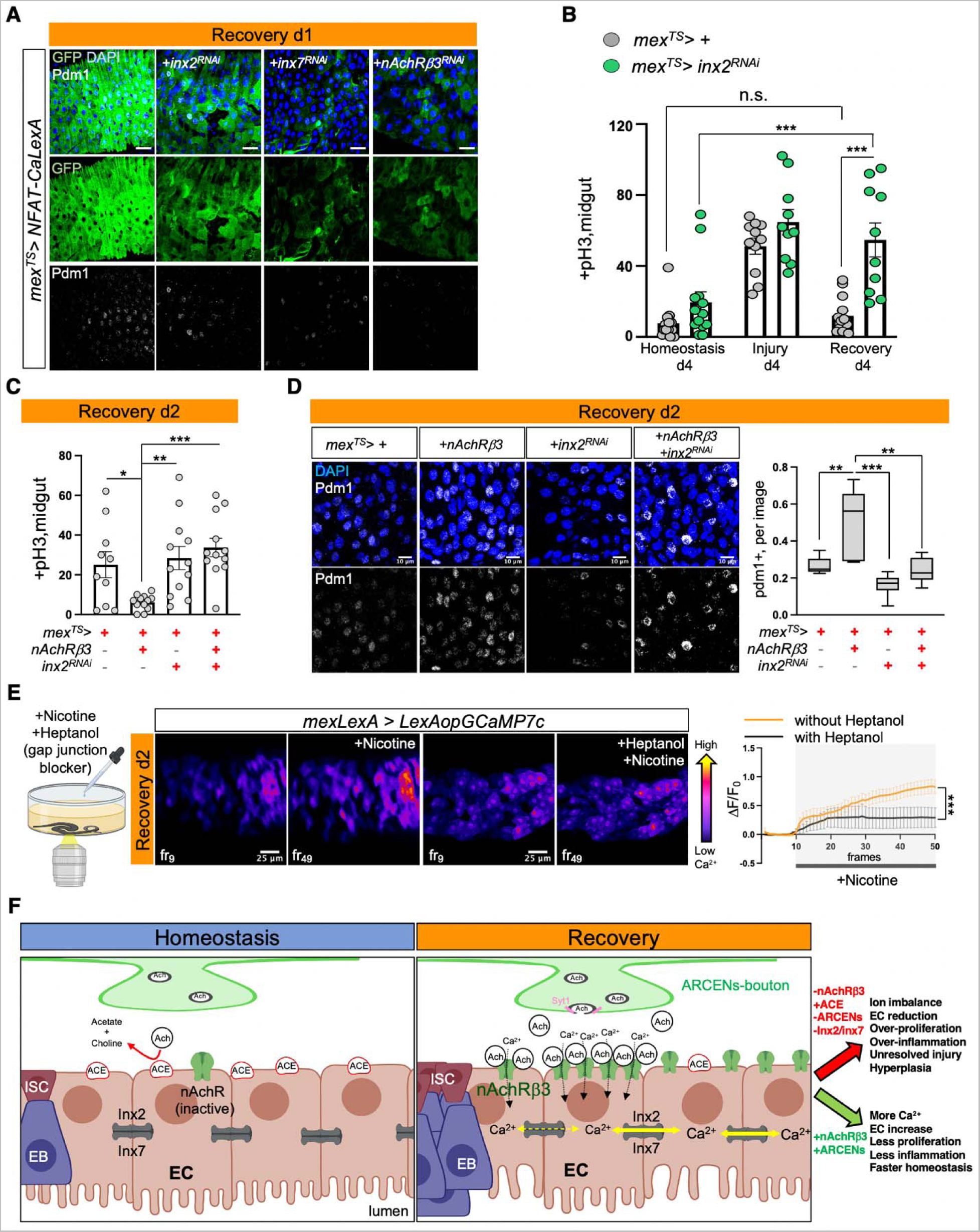
nAChR-mediated Ca^2+^ propagates via Inx2/Inx7 gap junctions. **(A)** Posterior midgut images with the NFAT-CaLexA Ca^2+^ reporter in control guts during Recovery d1 (*mex^TS^ > NFAT- CaLexA)* and when *inx2* or *inx7* or *nAChRβ3* is conditionally knock down in ECs (*mex^TS^ >NFAT- CaLexA+inx2^RNAi^, mex^TS^ >NFAT-CaLexA+inx7^RNAi^, mex^TS^ >NFAT-CaLexA+nAChRβ3^RNAi^*). Endogenous Ca^2+^ in ECs is visualized with anti-GFP (green) while mature ECs are marked with anti-pdm1 (grey). Scalebar 20μm. Flies were fed DSS-food for 4 days at 23°C and then transferred to standard food at 29°C for 24hrs. **(B)** Mitotic division counts from midguts of control (*mex^TS^ > +*, grey) flies and flies with conditional decrease of *inx2* in ECs (*mex^TS^> inx2^RNAi^*, green). Conditions are like in Fig. 1G. N=10-14 guts. **(C)** Midgut mitotic division counts of flies with conditional overexpression of *nAcRβ3* in ECs (*mex^TS^>nAcRβ3),* conditional knock down of *inx2* in ECs (*mex^TS^>inx2^RNAi^*) and flies with both (*mex^TS^>nAcRβ3+inx2^RNAi^*) during Recovery d2. Control: *mex^TS^ > +*. Flies were fed for 4 days DSS-food at 23°C and then transferred to standard food for ∼48hrs at 29°C. N=10-12 guts per genotype. **(D)** Posterior midgut images during Recovery d2 from control flies (*mex^TS^ > +*), *mex^TS^>nAcRβ3, mex^TS^>inx2^RNAi^* and *mex^TS^>nAcRβ3+inx2^RNAi^* flies. Conditions are like in Fig. 5C. Mature ECs were visualized with anti-pdm1(grey). Accompanying boxplot shows the ratio of pdm1+ nuclei. N=6-7 guts per genotype. DAPI: blue (nuclei). **(E)** Experimental illustration and color-coded sequential midgut frames from *mexLexA > LexAopGCaMP7c* flies during Recovery d2 before (fr_9_) and after (fr_49_) compound administration (nicotine alone or nicotine and heptanol) (Movies S8-S9). Following graph shows the relative fluorescence intensity (ΔF/F_0_) per frame per condition (+Nicotine -Heptanol: orange line, +Nicotine+Hepantol: black line). N=4 guts per condition. **(F)** Model: During recovery, ECs become sensitive (*Ace* reduction) and receptive (*nAChRβ3* increase) to cholinergic signaling. This allows them to receive ACh from ARCENs and to trigger nAChR-mediated Ca^2+^ influx that propagates via Inx2/Inx7 gap junctions and advances transition to homeostasis. Reduction of this pathway causes ion imbalance and keeps the epithelium in a state of unresolved injury with propensity to hyperplasia by promoting EC loss, over-proliferation and over-inflammation. Whereas short-term induction increases Ca^2+^ among ECs which expedites transition to homeostasis. All p-values were obtained with one-way Anova or two-way Anova. *: 0.05>p>0.01, **: 0.01<p<0.001, ***: p<0.001. Mean + s.e.m.

To test whether gap junction signaling is downstream of nAChRs, we conditionally knocked down *inx2* in ECs while overexpressing *nAChRβ3* for 2 days. We found that knocking down *inx2* was sufficient to attenuate the rapid decrease in ISC proliferation and block fast increase of pdm1+ ECs, thereby blocking the expedited recovery triggered by *nAChRβ3* overexpression (Fig. 5C, Fig. 5D). Finally, we tested *in vivo* whether gap junctions regulate Ca^2+^ propagation by adding the gap junction blocker heptanol together with nicotine and recording intracellular Ca^2+^ changes in ECs (Fig. 5E). During recovery, addition of heptanol and nicotine prompted an uneven Ca^2+^ response among ECs and dampened Ca^2+^increase (Fig. 5E and Movies S8-S9). Altogether, our data support that Inx2/Inx7 gap junctions are required for nAChR-mediated Ca^2+^ to spread among ECs during recovery and disruption of this bioelectric signaling prevents the transition to homeostasis.

## Discussion

In this work we address a fundamental question in regenerative biology, how does transition to homeostasis occur after injury? We used the *Drosophila* intestinal epithelium to answer this question as the gut is frequently subject to insults during ingestion. We discovered that following injury the conserved cholinergic pathway directs the gut to return to its homeostatic state by coordinating diverse processes like neuro-epithelial interactions and bioelectric signaling. In detail, our data support that ECs exhibit higher sensitivity and receptivity to ACh during recovery due to changes in *Ace* and *nAChRβ3* expression, respectively. This elevated cholinergic responsiveness of ECs enables them to receive ACh from ARCEN innervations. ACh then opens nicotinic receptors that carry the β3 subunit. Subsequently, Ca^2+^ is elevated in ECs that reside near the vicinity of innervations. This in turn activates Inx2/Inx7 gap junctions spreading Ca^2+^ that advances mature ECs across the epithelium (Fig. 5F). Disrupting this process causes unresolved chronic injury consisting of EC reduction, ion imbalance, unwarranted Yki activation and release of inflammatory cytokines Upd2/3, Vn and Eiger, reminiscent of IBDs ^42,93,94^. Conversely, short-term over-activation of this process expedites transition to homeostasis.

### ARCEN/nAChR-induced Ca^2+^ waves among ECs during regeneration

Our findings reveal that nAChR-mediated Ca^2+^ influx in ECs is an essential checkpoint for recovery. This is consistent with previous reports in the airway epithelium that nAChR-mediated Ca^2+^ protects from hyperplasia after injury and promotes wound healing^95,96^. We propose that nAChR-mediated Ca^2+^ during gut recovery triggers regulatory mechanisms essential to epithelial maturation and function. For example, high Ca^2+^ influx after opening of nAChRs alters the function of other ion channels such as Cl^-^ channels, which is evident by the reduced Cl^-^ levels in the midgut upon *nAChRβ3* reduction (Ext. Fig. 3E, Ext. Fig. 3G). Interestingly, cholinergic receptors are reported to regulate Ca^2+^ signaling and to stimulate ion transport in the small intestine potentially by influencing the activation of Cl^-^ channel CFTR (cystic fibrosis transmembrane conductance regulator) whose mutations cause the multiorgan disease cystic fibrosis^55,97^. Therefore, one of the roles of nAChR-mediated Ca^2+^ in ECs is likely to stimulate ion transport and advance physiological gut functions like absorption and secretion.

The use of endogenous ion currents that electrically couple multiple cells through gap junctions and allow them to behave as one unit is linked to normal growth and tissue-patterning during development and regeneration^1,6,98,99^. We found this to be conserved during midgut regeneration since reduction of Inx2/Inx7 gap junctions in ECs causes irregular Ca^2+^ patterns and attenuates nAChR-mediated transition to homeostasis (Fig. 5A-E). Moreover, we discovered that elevated Ca^2+^ influx in ECs is first triggered locally by ARCEN-EC interactions and then spread by gap junctions. This is supported by the observation that cholinergic signaling from ARCENs to ECs is sufficient to initiate wide Ca^2+^ increase and is required for nAChR-mediated return to homeostasis (Fig. 4E, 4F, 4G, 4H and Ext. Fig. 5F, Ext. Fig. 5G), despite the limited epithelial coverage of ARCEN innervations (Fig. 4B).

### Nerve-dependent gut regeneration

Despite the increasing knowledge of the protective anti-inflammatory roles of peripheral neurons^67–69,82,83^, many aspects remain unclear, especially relating to intestinal epithelial renewal^18^. We observed that ARCEN innervations carry presynaptic boutons near ECs suggesting that ACh could be released directly to ECs during recovery (Fig. 4D, Movie S7, Ext. Fig. 5C, Ext. Fig. 5D). In addition, we discovered that ACh release from ARCENs is required during gut regeneration to promote nAChR-mediated Ca^2+^ influx, increase mature ECs as well as to reduce proliferation and to decrease gut inflammatory signals (Fig 4A-H, Ext. Fig 5F-H). Thereby, our data expand the current knowledge by finding direct roles for cholinergic enteric neurons in healing the intestinal epithelium, which could help advance our understanding of the link between neurological disorders and intestinal pathologies^100^.

The cholinergic anti-inflammatory reflex has been proposed to be a conserved neuro-immune response in mammals and nematodes counteracting inflammatory signals by first sensing them and then releasing ACh to reduce them^2,5,18,101^. We propose that a similar mechanism is conserved in *Drosophila*, consisting of enteric cholinergic neurons that respond to inflammatory signals when the gut is injured. Supporting this, conditionally reducing ACh in ARCENs during recovery increases conserved inflammatory cytokines *TNF/Egr* and *IL-6- like/Upd3* in the gut (Ext. Fig. 5F) and short ARCEN activation decreases them (Ext. Fig 5H). Further, conditionally knocking down the TNF receptor *wgn* in ARCENs causes gut over-proliferation specifically during recovery (Ext. Fig. 5I), resembling the phenotypes followed by reduction in cholinergic signaling.

Interestingly, we found that EC responses to ACh are subject to change during regeneration. Specifically, we discovered that ECs are more sensitive and responsive to ACh during recovery due to *Ace* downregulation and *nAChRβ3* upregulation (Fig. 1E-I, Fig 2D-G, Ext. Fig 2C). These changes in ECs together with the TNF-linked roles of ARCENs, indicate that during regeneration neuro-epithelial cholinergic signaling is plastic. We propose that these different levels of cholinergic regulation in neurons and epithelium work in coordination to precisely control the strength, initiation, and duration of nAChR-mediated Ca^2+^ currents across the regenerative gut.

### Epithelial cholinergic sensitivity during regeneration

For the intestinal epithelium to return to its homeostatic state after injury, two parameters need to be met. First, ISCs must return to quiescence once epithelial cells are replenished. Second, essential functions such as absorption and secretion must be restored to prior-to-damage levels (i.e. homeostatic levels) across the entire tissue. Thus, ISC-quiescent pathways are part of a larger-scale coordinated response, which orchestrates transition to homeostasis independent of the type of injury.

So far, BMP signaling is the only reported ISC-quiescent pathway in *Drosophila*, although it does not always influence ISC proliferation. For example, BMP signals are not triggered during DSS injury^39^ but regulate ISC proliferation after *Ecc15* infection or Bleomycin injury ^4,7,39^. Moreover, *AWD* induction, which is required for the dual functions of BMPs^4^, is not upregulated during DSS-recovery (our data) but is upregulated after bacterial infections ^4,46^(Flygut-EPFL). Also, over-proliferation triggered by knocking down *nAChRβ3* in ECs did not change the expression of BMP ligands Dpp and Gbb during DSS-recovery (Fig. 3A). However, *Ace* is consistently downregulated during recovery after DSS treatment or after different bacterial infections (Fig. 1F, Flygut-EPFL)^46^. In addition, our data support that overexpression of *Ace* in ECs prevents return to homeostasis independent of the type of epithelial damage (Fig.\ 1G, Ext. Fig. 1J).

Therefore, we propose that increase in intestinal epithelial cholinergic sensitivity due to *Ace* downregulation is essential for healing the epithelium and transitioning to homeostasis independent of the type of damage. It would be of great interest to explore if *Ace* downregulation is a conserved intestinal epithelial response after injury, especially since ACE/AChE inhibitors are tested as therapeutics for intestinal pathologies like IBDs^18,48–51^.

### Neuro-induced epithelial bioelectric signaling during gut regeneration

Bioelectric signaling during epithelial healing and regeneration has gained a lot of attention due to the benefits of using exogenous electric fields to enhance recovery^6,99,102–106^. However, dissecting *in vivo* how endogenous bioelectric signaling is initiated during regeneration and which specific regulatory networks are downstream, has been challenging despite having significant therapeutic value. Here, we provide an example *in vivo* of an endogenous neuro-induced epithelial bioelectric mechanism during gut regeneration. We found that ARCEN/nAChR-controlled Ca^2+^ flows exert their regenerative properties in the intestinal epithelium by promoting ion balance (Ext. Fig. 3E, Ext. Fig. 3G) and by reducing inflammatory cytokines (Fig. 3A, Fig. 3I). In addition, short-term ARCEN activation, reminiscent of vagus-nerve stimulation^107^, is sufficient to spread Ca^2+^ among ECs and enhance epithelial healing (Fig. 4H, Ext. Fig. 5G-H). Our findings are also supported by a recent study that detected Ca^2+^ waves in R3 of the midgut^108^. Relevant to our findings cholinergic, nicotinic, Hippo, TNF pathways, as well as vagus-nerve stimulation, are being tested as therapeutics for intestinal pathologies and cancer^9,10,13,18,93,109–114^. Therefore, this study introduces a potential link between bioelectric interventions and canonical treatments, which could ultimately advance therapeutics for chronic disorders like IBDs and cancer.

Altogether, our findings demonstrate how the conserved cholinergic pathway facilitates the cooperation of diverse mechanisms such as neuro-epithelial communication and bioelectric signaling to initiate and expand with precision endogenous Ca^2+^ currents that heal the epithelium and direct it towards homeostasis. This work expands our current knowledge of regeneration and could help identify the etiology of various disorders as well as provide new leads to therapeutic treatments.

## Acknowledgments

We thank Stephanie Mohr and Justin Blau for critical comments on the manuscript. Confocal imaging was conducted at MicRoN Facility at Harvard Medical School, and we thank Paula Montero Llopis for advice on imaging. We thank the Cepko lab at Harvard Medical School for sharing their vibratome. We also thank Hugo Bellen , Mike Levin and Jayaraj Rajagopal for discussions, Guy Tanentzapf, Kate O’Connor-Giles, Xiaohang Yang, Chris Potter, Hugo Bellen, Todd R. Laverty, Gerry Rubin, Janelia FlyLight, DSHB, DRSC/TRiP, VDRC and the Bloomington Stock Center for fly lines, antibodies, and reagents. We thank Sudhir Gopal Tattikota for advice on single nuclei sequencing, Rich Binary, Alice Zheng, Haofan Li for help in this project as well as Pedro Saavedra, Liz Lane, Jacob Paiano, David Doupé, Justin Bosch, Ben Ewen-Campen, Lucy Liu, Charles Xu, Patrick Jouandin, Misty Rose Riddle, Tyler Huycke, Frederik Wirtz-Peitz and Chiao-Lin Chen for advice and reagents. We thank Christians Villalta and Bestgene for the fly injections, Hunter Elliot and Marcelo Cicconet at the Image and Data Analysis Facility (IDAC) at Harvard Medical School for advice on Imaris segmentation, Shahar Alon for advice on expansion microscopy, and the Biopolymer and Computing facilities as well as the PCMM Flow Cytometry Facility at Harvard Medical School. Illustrations were created with BioRender.com.

## Author contributions

Conceptualization: AP, NP; Methods, Data Acquisition, and Visualization: AP, Resources, AP, AC, YL, YH, YL ; Writing – original draft: AP, Writing – review & editing: AP, NP.

## Data and materials availability

Raw RNA-seq reads have been deposited in the NCBI Gene Expression Omnibus (GEO) database under accession code: (XXX, will provide soon). The rest of the data are available in the main text, supplementary materials, and movies.

## Materials and Methods

### Fly stocks

Flies were crossed and raised between 19-23°C in standard fly food. All adult flies were tested 3-5 days after hatching. All experiments were done in female flies, except for experiments in Ext.Fig. 4 where both female and male flies were used to characterize cholinergic enteric innervations.

The following stocks used in this study have been described previously. *<u>Drivers:</u> mex-Gal4*^115^, *mex-Gal4 Tubulin-Gal80^TS^* (*mex^TS^*), *mex-Gal4 UAS-2x-GFP (mex > GFP), esg-sfGFP* (*esg-GFP*, generated by David Doupé)^116^, *ChAT^MI04508^-Gal4* (*ChAT-Gal4*, BDSC:60317)^117^, *ChAT^MI04508^-Gal4; Tubulin-Gal80^TS^* (*ChAT^TS^*), *ChAT^MI04508^-QF* (*ChAT-QF*, BDSC:60320)^117^, *ChAT^MI04508^-Gal80* (*ChAT-Gal80*, BDSC:60321)^117^, *Tsh-Gal80* (gift from Todd R. Laverty), *R49E06-Gal4* (*ARCENs*, BDSC:38689), *Tubulin-Gal80^TS^*; *R49E06-Gal4* (*R49E06*^TS^ and *ARCENs^TS^*), *Tubulin-Gal80^TS^* (BDSC:7018,7019), *Diap1-LacZ* (BDSC:12093). Perrimon lab stocks: *Myo31DF*^NP0001^*-Gal4* (*myo1A-Gal4*), *myo1A-Gal4 UAS-GFP* (*myo1A > GFP*), *Myo1A-Gal4 Tubulin-Gal80^TS^*(*myo1A^TS^*), *esg-Gal4 Tubulin-Gal80^TS^ (esg^TS^*), *Tubulin-Gal80^TS^; esg-Gal4 (esg^TS^*)*, esg-Gal4*, *Su(H)GBE-Gal4 UAS-CD8-GFP; Tubulin-Gal80^TS^* (*Su(H)GBE^TS^*), *Tubulin-Gal80^TS^; pros-Gal4* (*pros^TS^*), *elav-Gal4; Tubulin-Gal80^TS^* (*elav^TS^*), *+; hml-Gal4Δ UAS-GFP; Tubulin-Gal80^TS^*(*hml^TS^*), *Tubulin-Gal80^TS^ ; how^24B^-Gal4* (*how^TS^*). *<u>Reporters:</u> 20XUAS-IVS-jGCaMP7c* (*UAS-GCAMP7c,* BDSC: 79030), TOE.GS01624 (*gRNA-ACE*, BDSC: 79471), *UAS-3XFLAG-dCas9-VPR(49)* (*UAS-dCas9VPR*, BDSC: 66562), *UAS-nAChRβ3^RNAi^*(JF01947, BDSC: 25927)^118^, *UAS-2x-GFP* ( *2xGFP*, BDSC: 6874), *LexAop-CD8-GFP-2A-CD8GFP;UAS-mLexA-VP16-NFAT LexAop-rCD2-GFP* (*NFAT-CaLexA*, BDSC:66542)^60^, *UAS-Orai*^119^,*UAS-ChAT^RNAi^* (JF01877, BDSC:25856)^118^, *UAS-sc^RNAi^* (JF02104, BDSC: 26206)(*59*), *20XUAS-6XGFP* (*6xGFP*, BDSC: 52262), *10XUAS-mCD8::GFP* (*mCD8:GFP*, BDSC:32184, 32186), *UAS-mCD8::GFP QUAS-mtdTomato-3xHA* (*UAS-GFP QUAS-Tomato-HA*, BDSC:30118), *QUAS-mtdTomato-3xHA* (*QUAS-Tomato-HA*, BDSC: 30005), 20xUAS-6xmCherry-HA (*6xmCherry*, BDSC: 52268), *5xUAS-IVS-Syt1::smGdP-HA* (*Syt1-HA*, BDSC:62142)^120^, *10xUAS-IVS-myr::GFP* (*myrGFP*, BDSC: 32198), *13xLexAop2-6xGFP* (*LexAop6xGFP*, BDSC: 52265), *UAS-shibire^TS^*(BDSC: 44222), *QUAS-CsChrimson* (gift from Chris Potter), *13xLexAop-sfGFP* (*LexAopGFP*, generated by Pedro Saavedra), *UAS-TrpA1*^121^, *UAS-wgn^RNAi^* (HMC03962, BDSC: 55275), *UAS-grnd^RNAi^* (GD12580/v43454), *UAS-Inx2^RNAi^* (JF02446, BDSC: 29306), *UAS-Inx7^RNAi^* (JF02066, BDSC:26297), *13xLexAop-IVS-jGCaMP7c* (*LexAopGCaMP7c*, BDSC:80916), *UAS-emptyVK37* (gift from Hugo Bellen), *UAS-Luciferase^RNAi^* (JF01355, BDSC: 31603), *UAS-Luciferase* (BDSC: 35789).

The following stocks were generated in this study. *<u>Reporters</u>*: 1) To generate the *UAS-nAChRβ3^RNAi-2^* line, we used the Valium20 vector and followed the protocol as described in Ni et al^122^. Guide strand: CGGCGAGAAGATCATGATCAA, Passenger strand: TTGATCATGATCTTCTCGCCG. 2) To generate the *UAS-nAChRβ3* and *LexAop-nAChRβ3* line, we recombined *nAChRβ3* cDNA from DmCD00481061 (from FlyBi Consortium) into pValium10-roe and LexAop-GW, respectively, using Gateway™ LR Clonase™ II Enzyme Mix (Invitrogen). *nAChRβ3^RNAi-2^, nAChRβ3* and *LexAop-nAChRβ3* was integrated into the attp40 site following injection as described in Ni et al^122^. *LexAop-nAChRβ3* was also integrated into the VK27 site by BestGene Inc. *<u>Drivers</u>*: 1) To generate *mex-LexA::GAD,* we linearized the pDPPattB-LHG^77^ vector with Acc65I and BssHII enzymes and cloned gBlock1 and gBlock2 (Sup. Txt, generated by IDT) in the vector using Gibson Assembly (NEB). *Mex-LexA::GAD* was integrated into attp40 and attp2 sites^122^ as well as VK27 site (BestGene). 2) To generate *R49E06-QF* (*ARCENs-QF*), we amplified the R49E06 promoter with primers R49E06QF-F and R49E06QF-R from *Ore R* gDNA. R49E06QF-primers were based on the primers used to generate *R49E06-Gal4*^76^. We next linearized pattB-DSCP-QF#7-hsp70 (Plasmid #46133)^123^ using BamHI and NsiI and with Gibson Assembly (NEB) cloned *R49E06* promoter in the plasmid.

R49E06QF-F: ACATCCAGTGTTTGTTCCTTGTGTAGACTGACATTCCGTTGCCAAGAAGCGC

R49E06QF-R: TCGATCCCCGGGCGAGCTCGGATCAGCGTGTCCTGTAGTACCAGCATAC

*R49E06-QF* was integrated into attp40 and attp2 sites^122^.

### nAChRβ3-flag

We generated the nAChRβ3-flag genomic construct using Scarless gene editing as described in *Gratz et al*^124^. In brief, to generate ∼ 1kb Left and Right homology arms that flank *nAChRβ3* cleavage site (2L: 546,835-546,836), we designed primers Left-F, Left-R, Right-F and Right-R (see below). Right-R primer included a silent mutation to disrupt the PAM sequence. We used gDNA from Cas9 flies to amplify the homology arms which we cloned to pScarlessHD-3xFLAG-DsRed plasmid (gift from Kate O’Connor-Giles, Addgene plasmid # 80820) with Gibson Assembly (NEB).

Left-F: AATTGTAATACGACTCACTATAGGGGCGTGTGGCAACCGT,

Left-R: ACCTCCAGATCCACCACCTCGGTGTGGTTGATGCCCA

Right-F: TTCTGGTGGTTCAGGAGGTTCCGCCGTGTGCCAAGG

Right-R: AATTAACCCTCACTAAAGGGATTCCACGGTCTGATGGCCG.

*nAChRβ3* gRNA guides were cloned into the pCFD4 vector (Addgene #49411)^125^ as described at http://www.crisprflydesign.org/wp-content/uploads/2014/06/Cloning-with-pCFD4.pdf

Guide strand: ATCAACCACACCGAGGTGCC, Passenger strand: GGCACCTCGGTGTGGTTGATC. After injection, once CRISPR alleles were identified with the DsRed marker, we used pBac transposase to generate scarless *nAChRβ3-flag* genomic flies as described at https://flycrispr.org/scarless-gene-editing/.

### Inducing injury and recovery

For DSS-injury, flies were fed daily with 2% DSS (Dextran Sulfate Sodium Salt; MP Biomedicals) mixed in standard fly food for 4 days at either 23°C or 29°C as specified. To induce recovery following DSS-injury, flies were transferred to standard fly food at 29°C as specified. For immunocytochemistry experiments, fly guts were cleared for 2hrs with 5% sucrose when testing for homeostasis and recovery, and with 5% sucrose together with 2% DSS when testing for injury. For bacterial infection flies were starved for 2hrs and then fed with *Ecc15* at OD100 in 5% sucrose. To test for homeostasis, flies were processed as the experimental conditions without DSS-feeding.

### Immunocytochemistry

Vibratome-sectioned flies: Whole flies were fixed in 4% PFA and 0.4mg/ml Pierce^TM^ DSP (Thermo Scientific) in PBS while rotating for 4-5 hrs at 4°C. Fixed flies were then placed in warm 4% agarose solution inside a 15mmx15mmx5 mold/adaptor (VWR) and left on ice for 30min. Agarose coated flies were sectioned in 150μm slices with a Leica VT1000 vibratome in cold PBS using MX35 Ultra microtome blades (Thermo Scientific). Sectioned flies were removed from agarose and fixed again for 15min at RT. Samples were washed with 0.25% Triton-X in PBS 3 times for 10min at RT and then transferred to Blocking solution (5% Normal Donkey Serum, 0.25% Triton X-100 in PBS) overnight at 4°C. Primary antibodies were added with blocking solution for 48hrs at 4°C. Sectioned flies were then washed with 0.25% Triton-X in PBS, 5 times for 30min at 4°C. Secondary antibodies, Phalloidin and DAPI were added in blocking solution at 4°C overnight in the dark. Samples were then washed with 0.25% Triton-X in PBS, 5 times for 30min at 4°C in the dark and mounted with Vectashield medium using bridge and covered with No.1 thickness cover glass (ThermoScientific).

Dissected guts: Guts were dissected in cold PBS, fixed in 4% PFA in PBS for 25min at RT, washed 3 times for 10min in PBS and incubated for 1hr in Blocking solution (5% Normal Donkey Serum, 0.1% Triton X-100 in PBS). They were then stained with primary antibodies in Blocking Solution overnight at 4°C. Afterwards, guts were washed 3 times for 10min in 0.1% Triton X-100 in PBS and stained with secondary antibodies and DAPI in Blocking Solution at 4°C overnight in the dark. Guts were washed with 0.1% Triton X-100 in PBS 3 times for 10min and mounted in Vectashield medium.

Dissected TAG and Brain: Thoracicoabdominal ganglia (TAG) were dissected in cold PBS and fixed in 4% PFA in PBS for 30min at RT. They were washed for 10min with 1% Triton X-100 in PBS then 2 times for 10 min in 0.5 % Triton X-100 in PBS, incubated for 1hr in Blocking solution (5% Normal Donkey Serum, 0.25% Triton X-100 in PBS) and then stained with primary antibodies in Blocking Solution overnight at 4°C. Afterwards they were washed 3 times for 10min in 0.25% Triton X-100 in PBS. Next, they were stained with secondary antibodies and DAPI in Blocking Solution at 4°C overnight in the dark and washed with 0.25% Triton X-100 in PBS 3 times for 10min before mounted in Vectashield medium.

The following antibodies were used: rabbit anti-Flag (Sigma, F7425; 1:100), mouse anti-β-galactosidase (Promega Z378A; 1:500), rabbit anti-pH3 (Millipore #06-570; 1:3000), mouse anti-GFP (Invitrogen A11120; 1:500), mouse anti-ChAT (DSHB; 1:50) rabbit anti-GFP (Invitrogen A6455; 1:3000), rabbit anti-DsRED (Clontech #632496; 1:200), chicken anti-GFP (AVES; 1:2000), rat anti-HA (Sigma 3F10; 1:500), mouse anti-Pros (DSHB MR1A; 1:50), rabbit anti-Syt1 (gift from Hugo Bellen; 1:1500), rabbit cleaved anti-Dcp1(Cell Signaling Asp216; 1:100), rabbit anti-pdm1 (gift from Xiaohang Yang; 1:500), mouse anti-pdm1 (DSHB 2D4; 1:20), guinea pig anti-inx2 (gift from Guy Tanentzapf; 1:1000), Phalloidin Alexa-647 (Invitrogen A2284; 1:50), Phalloidin Alexa-633 (Invitrogen A2287; 1:50), Phalloidin Alexa-405 (Invitrogen A30104; 1:50), DAPI (1:3000), Alexa Fluor-conjugated donkey-anti-mouse, donkey-anti-rabbit, goat-anti-chicken, goat-anti-guinea pig and donkey-anti-rat secondary antibody (ThermoFisher; 1:1000).

### Ion dye assays

MQAE and SodiumGreen assays were conducted as described in Kim et al^126^. Specifically, flies were fed 2μM Sodium Green Tetraacetate Indicator (Invitrogen, S6900) or 2.5mM MQAE ((N-(Ethoxycarbonylmethyl)-6-Methoxyquinolinium Bromide) (Thermo Fisher, E3101) diluted in 5% sucrose overnight. Next guts were dissected and fixed in 4% PFA in PBS for 30min in the dark. After one rinse and one wash (10min) with PBS, DAPI or PI (1:3000) was added with PBS for 10min. Guts were washed one last time with PBS, mounted in Vectashield and taken immediately for confocal imaging.

### Optogenetic activation

For CsChrimson experiments flies were raised in the dark at 19°C. To induce recovery flies were first fed with 2%DSS-food for 4 days at 23°C in the dark before transferring to standard food. During recovery flies were at 29°C without light, except for pulsed red light (∼630nm) using 1-meter RGB LED Strip (SMD5050, eTopxizu) and fed standard food for 4hrs and then in 5% sucrose for 2hrs.

### Fluorescent imaging and data analysis

Confocal imaging was conducted with Zeiss LSM710 and LSM780 confocal microscope using 25x, 40x and 63x oil objective lenses with identical acquisition conditions for all samples of a given experiment. For whole fly imaging, tile scans were stitched while imaging. Fiji (https://imageJ.net/Fiji) was used to assemble all images and measure mean fluorescent. Brightness was adjusted equally across comparable images and background signal was subtracted (Despeckle) for clarity. 3D segmentation of confocal images was performed using Imaris provided by the IDAC facility of Harvard Medical School. The number of pH3+ cells were counted with an epi-fluorescence microscope. Prism (https://www.graphpad.com/scientific-software/prism/) was used to create all graphs as well as to perform statistical analysis with either unpaired two-tailed t-test or multiple comparisons with one-way Anova or two-way Anova (Tukey’s or Sidak’s multiple comparison test). All illustrations were created with BioRender.com.

### GCaMP7c live imaging

For *in vivo* live Ca^2+^ imaging, guts were dissected in HL3 buffer (1.5mM Ca^2+^, 20mM MgCl_2_, 5mM KCl, 70mM NaCl,10mM NaHCO_3_, 5mM HEPES, 115mM Sucrose, 5mM Trehalose) and placed in eight-well clear bottom cell culture chamber slides with HL3. Guts were stabilized with a Nylon mesh (Warner instruments, 64-0198) and paper clips cut in small identical pieces. Imaging was done using LSM710 and LSM780 microscopes with 40x water objective lens. Each frame is the maximum projection of 5-6 z-stacks (2.96μm/z) and was acquired with 488nm excitation for GFP. 5mM Acetylcholine (Acetylcholine Chloride, Sigma A2661) or 1mM Nicotine (Sigma, N3876) and 0.6mg/ml 1-Heptanol (Sigma, H2805) were added at the 10^th^ frame. Identical acquisition conditions were used for all samples and genotypes per experiment. All images were taken from similar areas in the posterior midgut between R4-R5. Fiji was used for assembly and calculation of fluorescence per frame. ΔF/F_0_ = F_fr_-F_0_/ F_0_. F_fr_ is the fluorescence per frame and F_0_ (baseline fluorescence) is the average fluorescence intensity of the first 9 frames (fr_1_-fr_9_).

### Quantitative Real-Time PCR

RNA was extracted from 15 adult fly guts or heads per biological sample using the Ambion PureLink RNA Mini Kit, including PureLink DNase. 500ng of RNA was amplified and converted to cDNA using the iScript cDNA Synthesis kit (Bio-Rad). cDNAs were analyzed using the SYBR Green kit (Bio-Rad) and Bio-Rad CFX Manager software. *Rpl32* or *α-tubulin* were used as internal controls. Each RT-qPCR was performed with at least three biological replicates.

*ACE*: F- AGGTGCATGTCTACACGGG, R- ACGTTGGTGTTGGGGTTCC

*Upd3*: F- ATCCCCTGAAGCACCTACAGA, R- CAGTCCAGATGCGTACTGCTG

*egr*: F- AGCTGATCCCCCTGGTTTTG, R- GCCAGATCGTTAGTGCGAGA

*nAChRβ3*: F- ATGACGACGACTCCCAAGATA, R- AAGAAGCATCCCCATTAGCATTT

*Pdm1*: F- AGCTGTCCTAACGAGTTCCG, R- ACATCGCGCATATTTGTGTCAA

*Esg*: F- ATGAGCCGCAGGATTTGTG, R- CCTCCTCGATGTGTTCATCATCT

*Prospero*: F- CTGCCCCAGAGTTTGGACAA, R- CCTGATGCGAGTGACTGGA

*Upd*: F- CAGCGCACGTGAAATAGCAT, R- CGAGTCCTGAGGTAAGGGGA

*Upd2*: F- CGGAACATCACGATGAGCGAAT, R- TCGGCAGGAACTTGTACTCG

*Vn*: F- TCACACATTTAGTGGTGGAAG, R- TTGTGATGCTTGAATTGGTAA

*Spi*: F- CGCCCAAGAATGAAAGAGAG, R- AGGTATGCTGCTGGTGGAAC

*Krn*: F- CGTGTTTGGCAACAACAAGT, R- TGTGGCAATGCAGTTTAAGG

*Dpp*: F- GCCAACACAGTGCGAAGTT, R- ACCACCTGTTGACTGAGTGC

*Gbb*: F- CGCTGGAACTCTCGAAATAAA, R- CCACTTGCGATAGCTTCAGA

*Wg*: F- GATTATTCCGCAGTCTGGTC, R- CTATTATGCTTGCGTCCCTG

*Hh*: F- GGATTCGATTGGGTCTCCTAC, R- GGGAACTGATCGACGAATCT

*ChAT*: F- CTGACACTCTACCCAAGGTGC, R- AGTAATAGGCCCAGTTATCCTCC

*Inx7*: F-CCTACAGGCCGGGAAGTGA, R-CAAAGGGCACCCACTGGTA

*α-tubulin*: F- CAACCAGATGGTCAAGTGCG, R- ACGTCCTTGGGCACAACATC

*Rpl32*: F- GCTAAGCTGTCGCACAAATG, R- GTTCGATCCGTAACCGATGT

### Expansion Microscopy

Expansion was conducted as described in *Asano et al.*^58^ In brief, guts were fixed, stained with primary and secondary antibodies (as described above), and after the final wash were placed briefly in PBS. Then PBS was exchanged with 0.1mg/ml AcX in PBS and guts were left overnight without shaking. The next day, gelation chambers were made consisting of a stack of 2 #1.5 coverslips in each side of a slide. Stock X, 4HT, TEMED and APS (47:1:1:1) were mixed at 4°C to generate the gelling solution. Guts were transferred in the solution and incubated for 30min in the dark at 4°C. Within 5 min after incubation, guts were placed in the gelation chamber (2 guts per chamber) with 20μl of gelling solution and covered with a lid without causing air pockets. The gelation chambers were then placed at 37°C for 2hrs in the dark and without moving. Next, using a razor the lid was removed and excess gel was trimmed off. A wet brush with digestion buffer (with ProK 1:100) was used to remove the gels (with the guts) which were then placed in a 6 well Glass Bottom Plate with Lid (Cellvis, 1-2 guts per well) and immersed in ∼2-3ml of digestion buffer overnight, at RT and in the dark. The next day, the digestion buffer was exchanged with PBS and stored at 4°C until imaging. Prior to imaging, gels were trimmed to the smallest possible size, then washed with UltraPure^TM^ Distilled water (Invitrogen) 3 times for 20min. During imaging most of the water was removed without drying the guts.

### Single nuclei gut profiling

Sample preparation: For homeostasis 3-5 days old *Ore R* female flies were fed standard lab food at 29°C for 6 days. For recovery d2 3-5 days old *Ore R* female flies were fed 2%DSS for 4 days at 29°C and then transferred to standard food at 29°C for 2 days. Single-Nucleus suspension and FACS: Single-nucleus suspension was conducted as described by Li et al^127^. Briefly, ∼70 guts per condition were dissected in cold Schneider’s medium, flash-frozen and stored at -80°C. Prior to FACs sorting, samples were spined down and Schneider’s medium was exchanged with homogenization buffer [250mM Sucrose, 10mM Tris pH8, 25mM KCl, 5mM MgCl, 0.1% Triton-X, 0.5% RNasin Plus (Promega, N2615), 50x protease inhibitor (Promega G6521), 0.1mM DTT]. Using 1ml dounce (Wheaton 357538), nuclei were released by 20 loose pestle strokes and 40 tight pestle strokes while keeping samples on ice and avoiding foam. Next, nuclei were filtered through 5 ml cell strainer (40 μm), and using 40 μm Flowmi (BelArt, H13680-0040). Nuclei were centrifuged, resuspended in PBS/0.5%BSA with 0.5% RNase inhibitor, filtered again with 40 μm Flowmi and stained with DRAQ7™ Dye (Invitrogen, D15106). Single nuclei were sorted with Sony SH800Z Cell Sorter at PCMM Flow Cytometry Facility at Harvard Medical School and 100k nuclei per sample were collected in PBS/BSA buffer.

10x genomics and sequencing: Single nuclei RNA-seq libraries were prepared using the Chromium Next GEM Single Cell 3’ Library and Gel Bead Kit v3.1 according to the 10xGenomics protocol. Approximately 16,500 nuclei were loaded on Chip G with an initial concentration of 700 cells/µl based on the ‘Cell Suspension Volume Calculator Table’. Sequencing was conducted with Illumina NovaSeq 6000 at Harvard Medical School Biopolymers Facility.

10x data processing: We used cellranger count pipeline 6.1.1 to process Chromium single-cell data and generated the feature-barcode matrices. The reads were aligned using the include-introns option with *Drosophila melanogaster* BDGP6.32 reference. All the matrices in different batches were aggregated into a single feature-barcode matrix by cellranger aggr pipeline and normalized by equalizing the read depth among libraries. Graph-based clustering in cellranger was applied to identify cell clusters (Fig. 1D). Heatmap (Fig. 1E) and Dot plot (Ext. Fig. 1D) were generated using the Seurat DoHeatmap and DotPlot functions. Statistics of the number of nuclei and gene expression per cluster were extracted from the single cell gene expression matrix and visualized by box plot and bar plot.

---

**Ext. Fig. 1.**
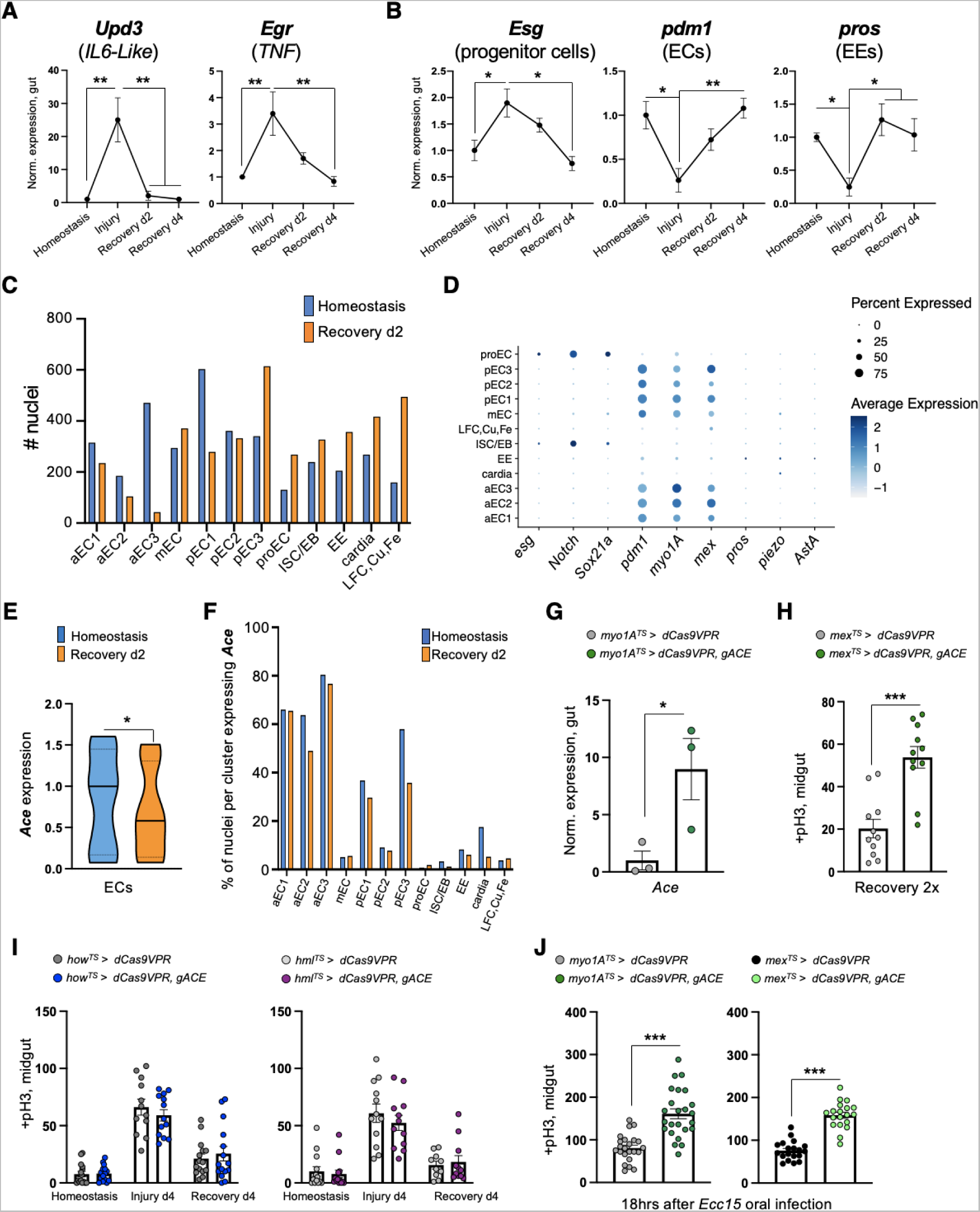
Sensitivity of ECs to ACh is required for recovery. **(A)** Expression levels of conserved inflammatory cytokines (*Unpaired-3*/*Upd-3* and *Eiger*/*egr*) in guts of *Ore R* flies undergoing DSS-induced repair. Expression levels are normalized to Homeostasis. N=3 biological samples per condition: homeostasis, injury, recovery d2, recovery d4. **(B)** Expression levels of markers for PCs (*escargot*, *esg*), ECs (*pdm1*), EEs (*prospero*, *pros*) in guts of *Ore R* flies undergoing DSS-induced repair. N=3 biological samples per condition. Expression levels are normalized to Homeostasis. **(C)** Number of nuclei recovered per cluster and per condition after gut single nuclei sequencing. **(D)** Dot plot illustrating the average expression (blue color range) and percent of expression (dot size) per cluster of marker genes for ECs (*pdm1*, *myo1A*, *mex*), EEs (*pros*, *piezo*, *AstA*) and for PCs (*esg*, *Notch*, *Sox21a*) [PCs: ISC/EB and proEC]. **(E)** Boxplot illustrating the mean expression level of *Ace* in EC clusters per condition. **(F)** Percentage of nuclei expressing *Ace* per cluster and condition. **(G)** Validation of *ACE* overexpression using CRISPR-OE. (**H)** Mitotic division counts from midgut of control (grey, *mex^TS^ > dCas9VPR*) flies and flies with conditional *Ace* overexpression (green, *mex^TS^ > dCas9VPR, gRNA-Ace*) in ECs during Recovery 2x (like Fig. 1H). N=11 guts per genotype. (**I)** Mitotic division counts from midgut of control (grey) flies and flies with conditional *Ace* overexpression in the visceral muscle (blue, *how^TS^ > dCas9VPR, gRNA-Ace*) and in hemocytes (immune cells, purple, *hml^TS^ > dCas9VPR, gRNA-Ace*). Conditions like Fig. 1G. N=11-15 guts per genotype per condition. **(J)** Mitotic division counts from midgut of control (grey, black) flies and flies with conditional *Ace* overexpression (green, *myo1A^TS^ > dCas9VPR, gRNA-Ace* and *mex^TS^ > dCas9VPR, gRNA-Ace* ) in ECs after infection (18hrs after *Ecc15* oral infection). Flies were transferred to 29°C upon *Ecc15* infection for ∼18hrs. N=19-24 guts per genotype. All p-values were obtained with one-way Anova or two-way Anova or t-test; *: 0.05>p>0.01, **: 0.01>p>0.001, ***: p<0.001, ns: non-significant. Mean ± s.e.m.

**Ext. Fig. 2.**
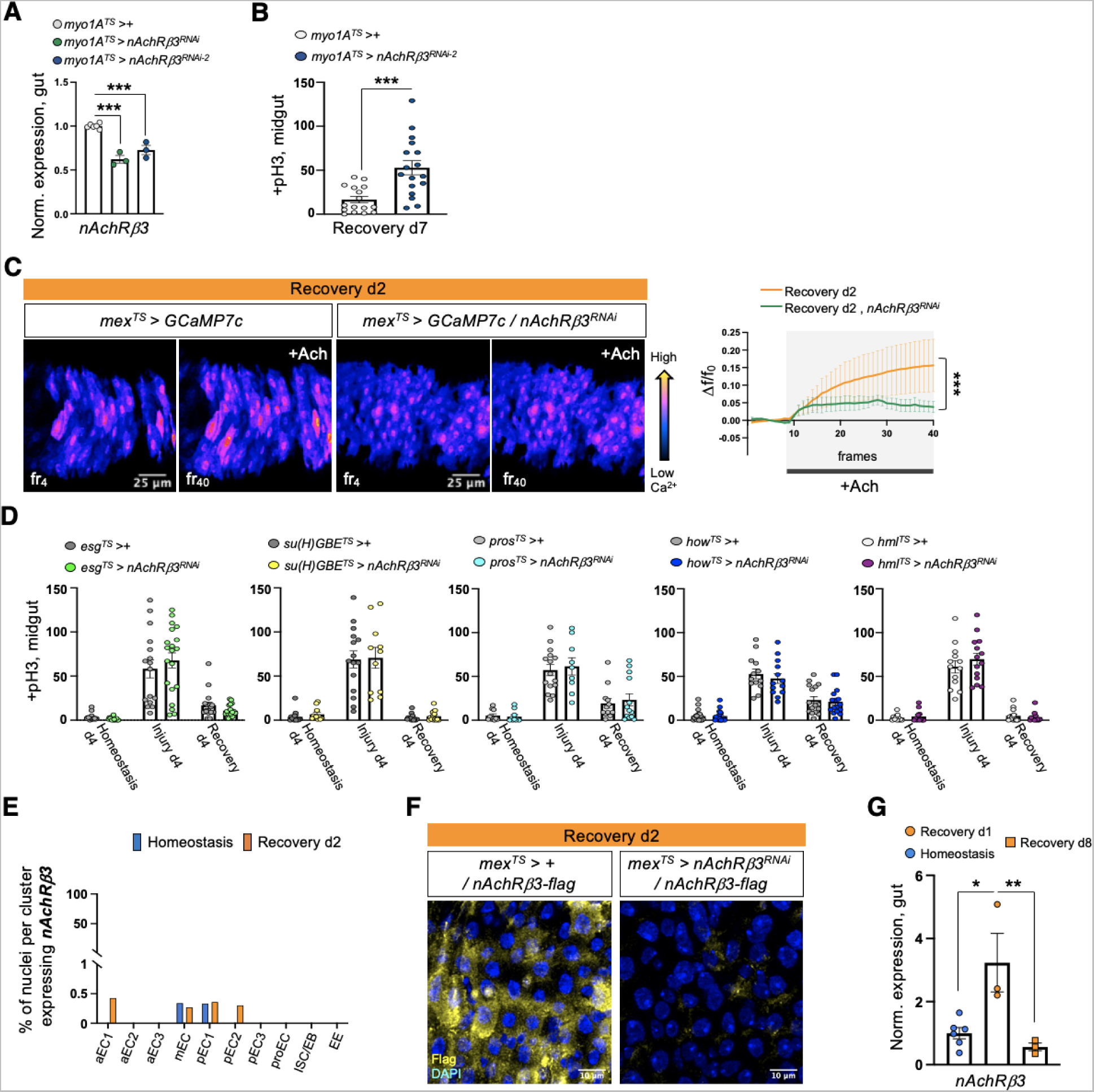
nAChRβ3 is required in ECs for recovery. **(A)** Validation of *nAcRβ3* RNAi knockdown with two different lines. **(B)** Mitotic division counts from midgut of control flies (*myo1A^TS^ > +,* white) and flies with *nAChRβ3* conditionally reduced in ECs (*myo1A^TS^ > nAChRβ3^RNAi-2^*, blue) 7 days after the end of DSS-feeding (Recovery d7). For Recovery d7, flies were transferred for 7 days to standard food at 29°C after 4 days of DSS-feeding at 23°C. **(C)** Color-coded sequential frames of midgut before (fr_4_) and after (fr_40_) ACh administration during Recovery d2 from *mex^TS^ > GCAM7c* flies and *mex^TS^ > GCAM7c +nAChRβ3^RNAi^* flies. Conditions are like in Fig. 1I. The accompanying graph demonstrates the relative fluorescence intensity (ΔF/F_0_) per frame and per genotype (Recovery d2: orange line, Recovery d2+*nAChRβ3^RNAi^*: green line). N=4 guts per genotype. **(D)** Mitotic division counts during Homeostasis d4, Injury d4 and Recovery d4 (like in Fig. 1G) from midgut of control flies (grey) and flies with conditional *nAcRβ3* reduction in PCs (*esg^TS^ > nAChRβ3^RNAi^*, green), EBs (*su(H)GBE^TS^ > nAChRβ3^RNAi^*, yellow), EEs (*pros^TS^> nAChRβ3^RNAi^,* cyan), visceral muscle (*how^TS^ > nAChRβ3^RNAi^,* blue) and hemocytes (*hml^TS^> nAChRβ3^RNAi^,* purple). N=10-20 guts per genotype per condition. **(E)** Percentage of nuclei expressing *nAChRβ3* per cluster in the gut during Homeostasis (blue) and Recovery d2 (orange). **(F)** Validation of nAChRβ3-flag construct. Confocal images of midgut during Recovery d2 from flies expressing nAChRβ3-flag (*mex^TS^ > +/nAChRβ3-flag*) and flies expressing nAChRβ3-flag while knocking down *nAChRβ3* in ECs (*mex^TS^ > nAChRβ3^RNAi^/nAChRβ3-flag)*. nAChRβ3-flag was visualized with anti-Flag (yellow) staining. DAPI: nuclei. **(G)** Expression levels of *nAChRβ3* in guts of *Ore R* flies during Homeostasis (blue circle), Recovery d1 (orange circle) Recovery d8 (homeostasis after injury, orange square). N=3-6 biological samples per condition. All p-values were obtained with one-way or two-way Anova or t-test; *: 0.05>p>0.01, **: 0.01<p<0.001, ***: p<0.001. Mean ± s.e.m.

**Ext. Fig. 3.**
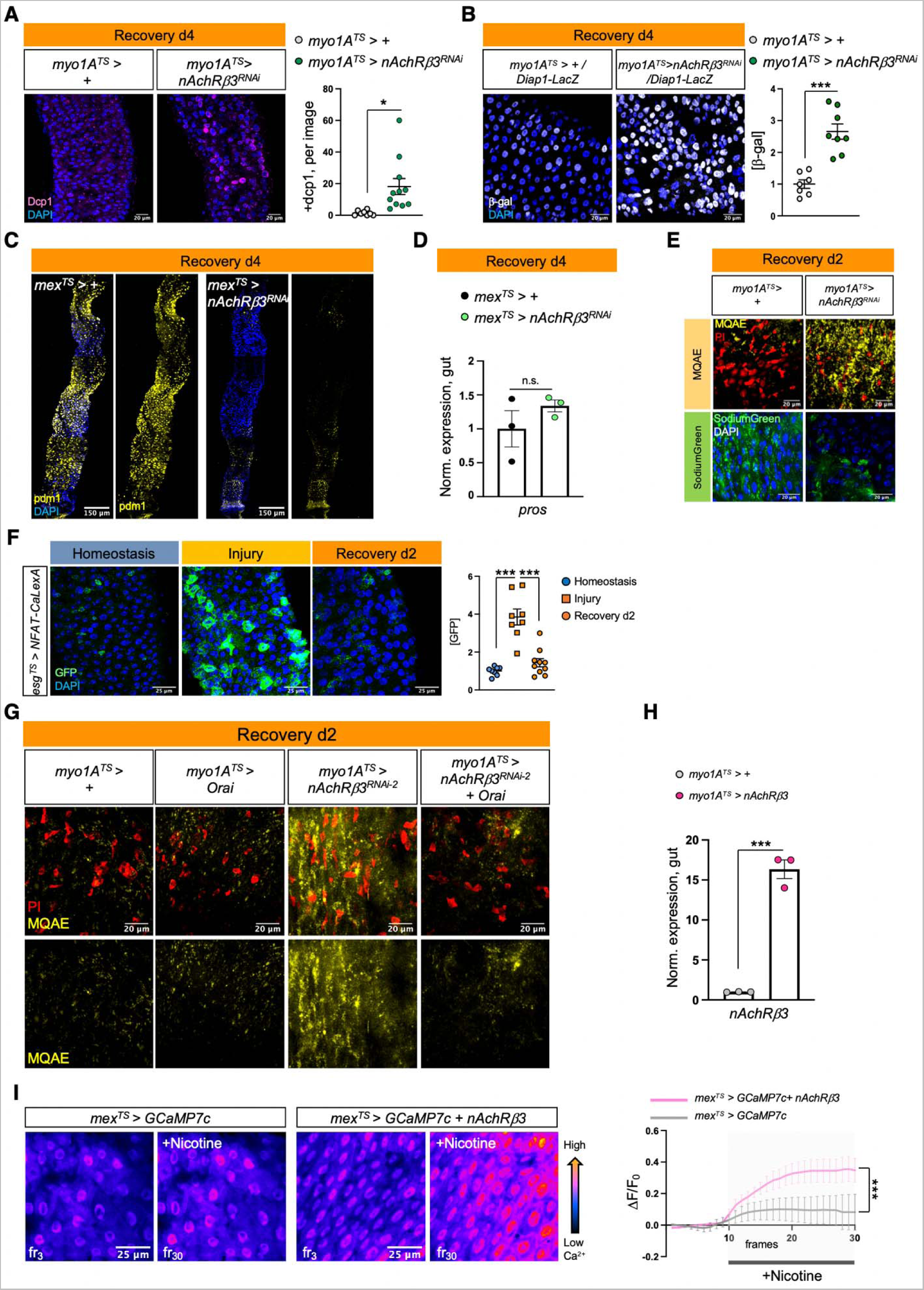
nAChRβ3-mediated Ca^2+^ increase in ECs promotes recovery. **(A)** Posterior midgut confocal images of *myo1A^TS^>+* and *myo1A^TS^> nAChRβ3^RNAi^*flies during Recovery d4 (like Fig.1G) stained for the cell death marker *Drosophila* caspase-1 (anti-Dcp1, pink). The following graph shows the Dcp1+ cells per image and genotype. N=8-11 guts per genotype. **(B)** Confocal images of posterior midguts during Recovery d4 (like Fig.1G) in control flies (*myo1A^TS^>+/Diap1-LacZ*) and flies with *nAChRβ3* conditionally knocked down in ECs (*myo1A^TS^> nAChRβ3^RNAi^ /Diap1-LacZ*) while expressing the LacZ reporter for the Yki target gene *Diap1*. LacZ was visualized with anti-β-gal staining (grey). The accompanying graph indicates the levels of β-gal per image and genotype. N= 7-8 guts per genotype. **(C)** Confocal images of posterior midgut during Recovery d4 (like Fig.1G) of *mex^TS^>+* and *mex^TS^> nAChRβ3^RNAi^* flies stained with the EC marker anti-pdm1 (yellow). **(D)** Expression levels of the EE marker *prospero* (*pros*) during Recovery d4 from guts of *mex^TS^>+* and *mex^TS^> nAChRβ3^RNAi^* flies. N=3 biological samples per genotype. Conditions are like in Fig.1G. **(E)** Confocal images of posterior midgut assayed with MQAE dye (detects intracellular Cl^-^ via diffusion-limited collisional quenching, yellow) and SodiumGreen dye (fluorescent indicator of intracellular Na^+^, green) from control flies (*myo1A^TS^>+*) and flies conditionally reducing *nAChRβ3* in ECs (*myo1A^TS^>nAChRβ3^RNAi^*). PI: nuclei (Propidium Iodide, red). Flies were fed DSS-food for 4 days at 23°C and then transferred at 29°C to standard food for 1 day and the 2^nd^ day to 5% sucrose and SodiumGreen or MQAE dye. **(F)** Posterior midgut images with PCs conditionally expressing the Ca^2+^ reporter *NFAT-CaLexA* (*esg^TS^ > NFAT-CaLexA*). PCs with high endogenous Ca^2+^ were visualized with anti-GFP (green). The reporter was expressed for 2 days (29°C) per condition. The accompanying graph shows the fold change of fluorescence per image and condition. N=8-10 guts per condition. **(G)** Confocal images of posterior midgut assayed with MQAE (like in Ext. Fig. 3E) from control flies (*myo1A^TS^>+*), flies conditionally overexpressing *Orai* in ECs (*myo1A^TS^>Orai*), flies conditionally reducing *nAChRβ3* in ECs (*myo1A^TS^>nAChRβ3^RNAi^*) and flies conditionally knocking down *nAChRβ3* while overexpressing *Orai* in ECs (*myo1A^TS^>nAChRβ3^RNAi^ + Orai*). **(H)** Validation of *nAcRβ3* transgene. (**I)** Color-coded sequential frames of midgut live imaging from *mex^TS^ > GCAM7c* and *mex^TS^ > GCAM7c* + *nAChRβ3* flies before (fr_3_) and after (fr_30_) administering nicotine. Following graph shows the relative fluorescence intensity (ΔF/F_0_) per frame and genotype (*mex^TS^ > GCAM7c:* grey line, *mex^TS^ > GCAM7c + nAChRβ3*: pink line). Flies were kept in standard food and transferred at 29°C for 2 days. N=7 guts per genotype. DAPI: nuclei (blue). PI: nuclei (Propidium iodide, red). All p-values were obtained with one-way Anova or t-test; *: 0.05>p>0.01, ***: p<0.001, n.s.: non-significant. Mean ± s.e.m.

**Ext. Fig. 4.**
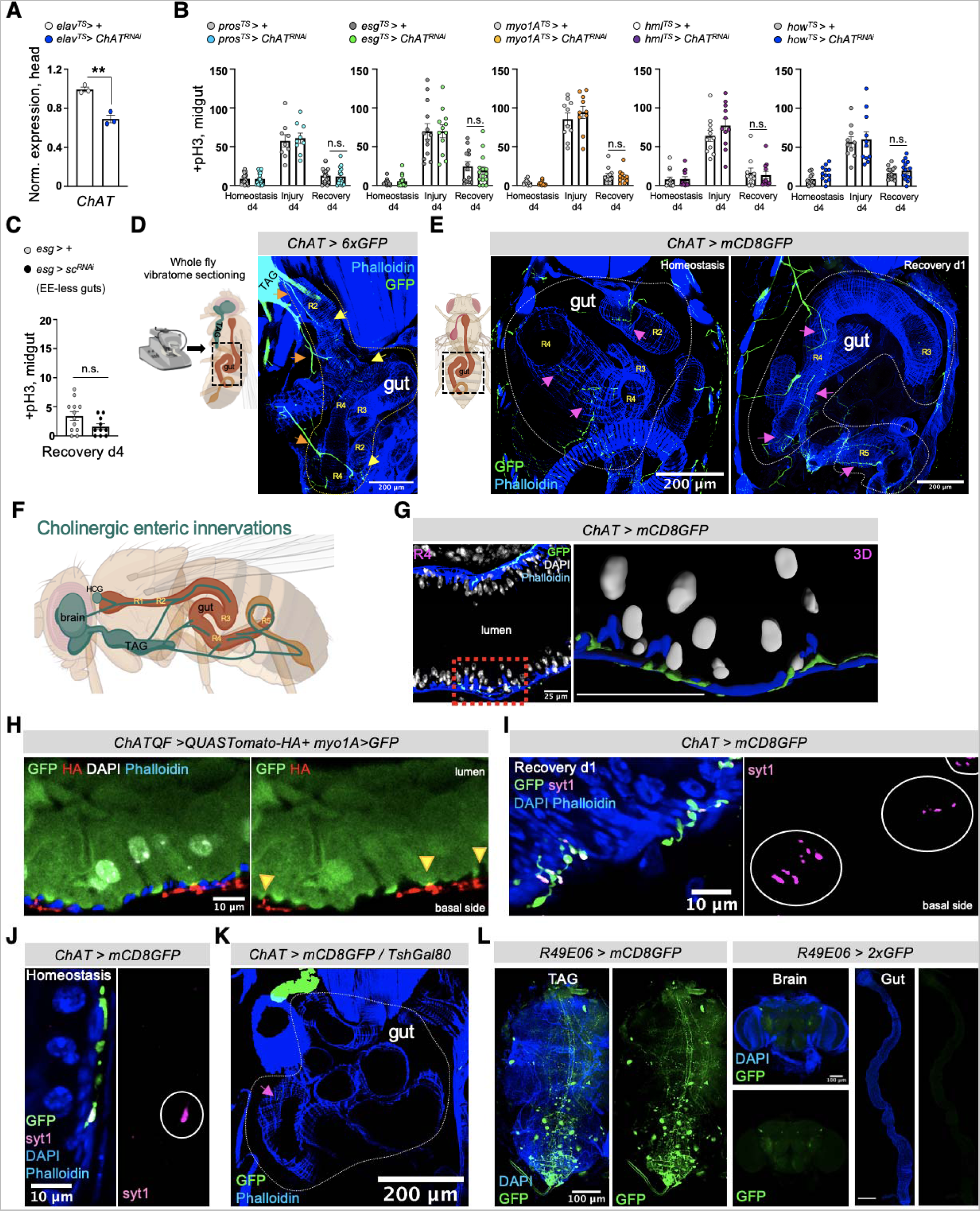
Neuro-EC interactions promote nAChR-mediated gut recovery. **(A)** Validation of *ChAT* RNAi knockdown with the neuronal driver *elav-Gal4*. N=3 biological samples per genotype. **(B)** Mitotic division counts from midgut of control flies (grey and white) and flies with conditional reduction of *ChAT* (*Choline Acetyltransferase,* enzyme responsible for ACh synthesis) using drivers for gut, visceral muscle, and immune cells: EEs (*pros^TS^>,* cyan), PCs (*esg^TS^*>, green), ECs (*myo1A^TS^>*, orange), hemocytes (*hml^TS^*>, purple), visceral muscle (*how^TS^>*, blue). Conditions are like in Fig.1G. N=10-15 guts per genotype. **(C)** Midgut mitotic division counts from control flies (*esg>+*) and flies without EEs in the midgut (*esg>sc^RNAi^*) during Recovery d4. N=10-12 guts per condition. **(D)** Schematic for imaging intact enteric innervations in abdominal midgut regions (R2, R3, R4, R5) and confocal image from a sectioned fly with the cholinergic driver *ChAT-Gal4* expressing GFP (*ChAT > 6xGFP*). Cholinergic neurons and innervations are stained with anti-GFP (green). Black dotted rectangular: region of the fly shown in confocal image. Orange arrows: arborizing of cholinergic enteric innervations, yellow arrows: cholinergic innervations in abdominal midgut region R2 and R4. **(E)** Confocal images from the same abdominal region (black dotted rectangular) after vibratome-sectioning of *ChAT>mCD8GFP* (membrane-targeted GFP) flies during Homeostasis and Recovery d1. pink arrows: cholinergic innervation in abdominal midgut region R2, R4 and R5. **(F)** Illustration of cholinergic enteric innervations after imaging vibratome-sectioned flies. TAG: Thoracicoabdominal ganglia. HCG: hypocerebral ganglia. **(G)** Confocal image of R4 midgut region innervated by cholinergic enteric neurons (*ChAT>mCD8GFP*) during Recovery d1 and stained with anti-GFP (green). Accompanying image is the 3D segmentation (using Imaris) of the innervated region indicated by the red dotted square (scale bar 25μm). DAPI: white (nuclei). Phalloidin: Blue (muscle). **(H)** Confocal image of innervated R2 in the abdominal midgut from vibratome-sectioned *ChAT-QF > QUAS-UAS-mtdTomato-HA/ myo1A > GFP (ChATQF>QUAS- HA + myo1A>GFP)* fly. Cholinergic enteric innervations (red) are marked with the cholinergic driver *ChAT-QF* expressing *mtdTomato-HA* and visualized with anti-HA staining. ECs are marked with GFP (*myo1A> GFP*) and stained with anti-GFP (green). yellow arrows: cholinergic innervations adjacent to ECs. DAPI: white (nuclei). **(I-J)** Confocal images of innervated R4 midgut regions from vibratome-sectioned *ChAT > mCD8GFP* flies during Recovery d1 (I) and Homeostasis (J). Cholinergic innervations are visualized with anti-GFP (green) and co-stained with the synaptic-vesicle marker Synaptotagmin1 (anti-Syt1, magenta). White circles: Syt1^+^ cholinergic *en passant* boutons (pre-synaptic swellings along the cholinergic enteric innervation). DAPI: blue (nuclei). Phalloidin: blue (muscle). **(K)** Abdominal confocal image (sectioned as in Ext. Fig. 4E) from a fly that is expressing GFP (green) in all cholinergic neurons except the ones residing in TAG (*ChAT>mCD8GFP /TshGal80*). Pink arrow: cholinergic innervations in the midgut. Phalloidin: muscle (blue). **(L)** Confocal images of TAG, brain, and gut (scale bar 200μm) from flies expressing the neuronal driver *R49E06-Gal4* (*R49E06>mCD8GFP* and *R49E06>2xEGFP*). anti-GFP: green. DAPI: blue. All p-values were obtained with one-way Anova or two-way Anova or t-test; **: 0.01<p<0.001, n.s.: non-significant. Mean ± s.e.m.

**Ext. Fig. 5.**
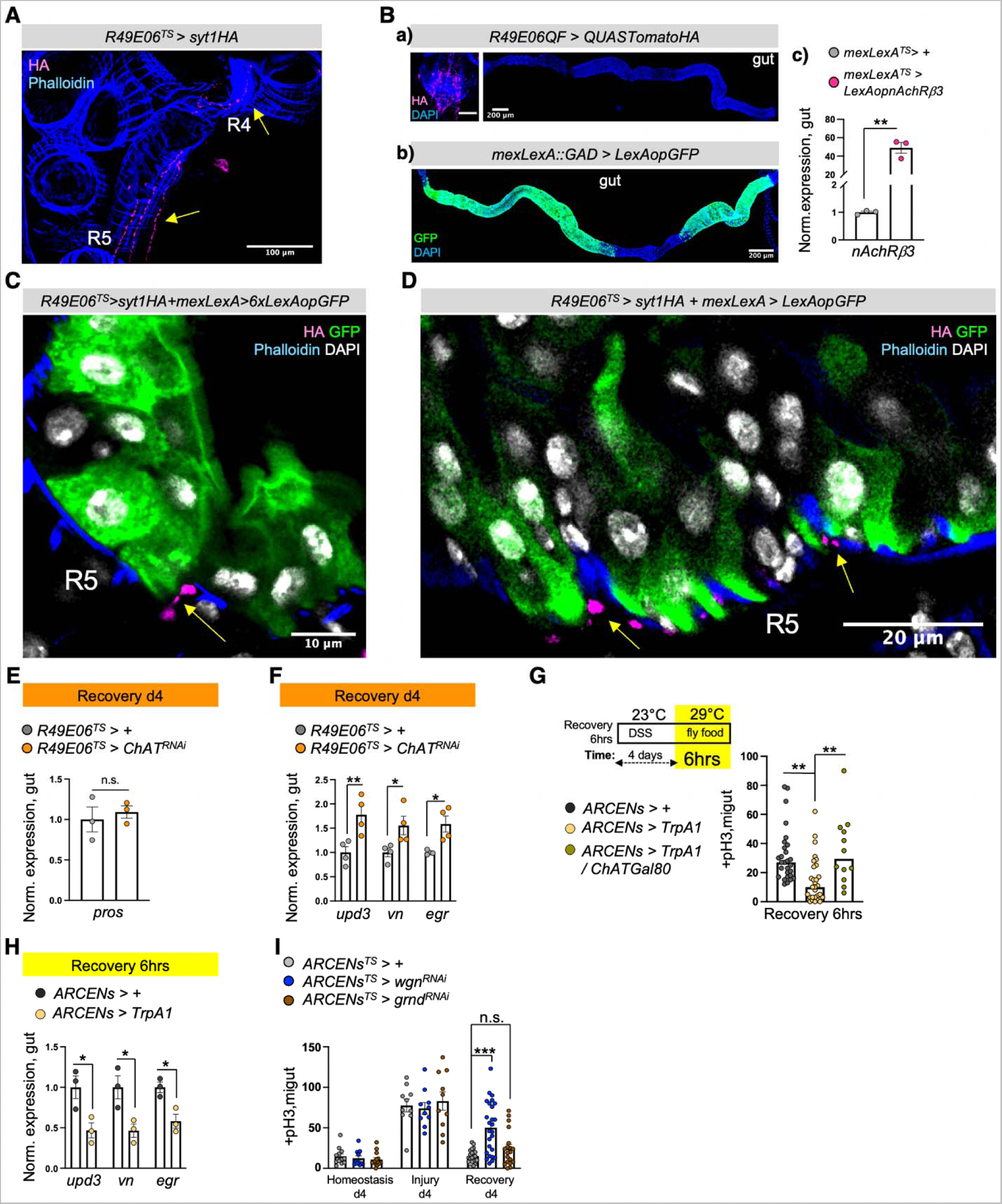
ARCEN-EC interactions promote nAChR-mediated gut recovery. **(A)** Confocal image from the abdomen of *R49E06^TS^>syt1HA* fly during Recovery d2 (sectioned like Ext. Fig. 4E). Yellow arrows point to R5 and R4 innervated by R49E06-innervations expressing Syt1HA (anti-HA, magenta). Phalloidin: muscle (blue). **(B)** Validation of a) *R49E06QF*: confocal images of posterior TAG (scale bar 100μm) and gut, b) *mexLexA* (*mexLexA::GAD)*: gut confocal image, c) *LexAop-nAChRβ3* transgene. **(C-D)** Confocal images showing R5 from *R49E06^TS^> syt1HA+ mexLexA> 6xLexAopGFP* and *R49E06^TS^>syt1HA+mexLexA> LexAopGFP* flies during Recovery d2 (Conditions like Fig. 4D). Yellow arrows: R49E06-innervations with the synaptic-vesicle protein syt1HA (anti-HA, magenta) in close proximity to ECs (anti-GFP, green). Phalloidin: muscle (blue). DAPI: white (nuclei). **(E)** Expression levels of the EE marker *pros* during Recovery d4 (like Fig. 1G) from guts of *R49E06^TS^>+* (grey) and *R49E06^TS^>ChAT^RNAi^*(orange) flies. N=3 biological samples. Levels normalized to control. **(F)** Expression levels of inflammatory cytokines *upd3*, *vn*, *egr* during Recovery d4 (like Fig. 1G) from guts of of *R49E06^TS^>+* (control, grey) and *R49E06^TS^>ChAT^RNAi^*(orange) flies. N=4 biological samples per genotype. Levels normalized to control guts. **(G)** Schematic of experimental conditions accompanied by mitotic division counts from control flies (*ARCENs>+,* black), flies with 6hrs thermo-activation of ARCENs (R49E06-neurons) with the TrpA1 channel (TrpA1-induction, *ARCENs>TrpA1*, yellow) and flies with TrpA1-induction in ARCENs while cholinergic neurons are inhibited by the *ChAT*-Gal80 repressor *(ARCENs > TrpA1/ChATGal80, grey*). N=10-29 guts per genotype per condition. **(H)** Expression levels of inflammatory cytokines *upd3*, *vn*, *egr* during the first 6hrs of Recovery (like Ext. Fig. 5G) from guts of *ARCENs^TS^>+* (control, black) and *ARCENs^TS^>TrpA1* (yellow) flies. N=3 biological samples per genotype. Levels normalized to control guts. **(I)** Mitotic division counts from control flies (*ARCENs^TS^>+,* grey) and flies with TNF receptors *wgn* (*wengen*, *ARCENs^TS^> wgn^RNAi^*, blue) and *grnd* (*grindelwald*, *ARCENs^TS^> grnd^RNAi^* , brown) knocked down in ARCENs for 4 days during Homeostasis, Injury and Recovery (like Fig. 1G). N=10-26 guts per genotype per condition. All p-values were obtained with one-way Anova or two-way Anova or t-test; *: 0.05>p>0.01, **: 0.01<p<0.001, ***: p<0.001. n.s.: non-significant. Mean ± s.e.m.

**Ext. Fig. 6.**
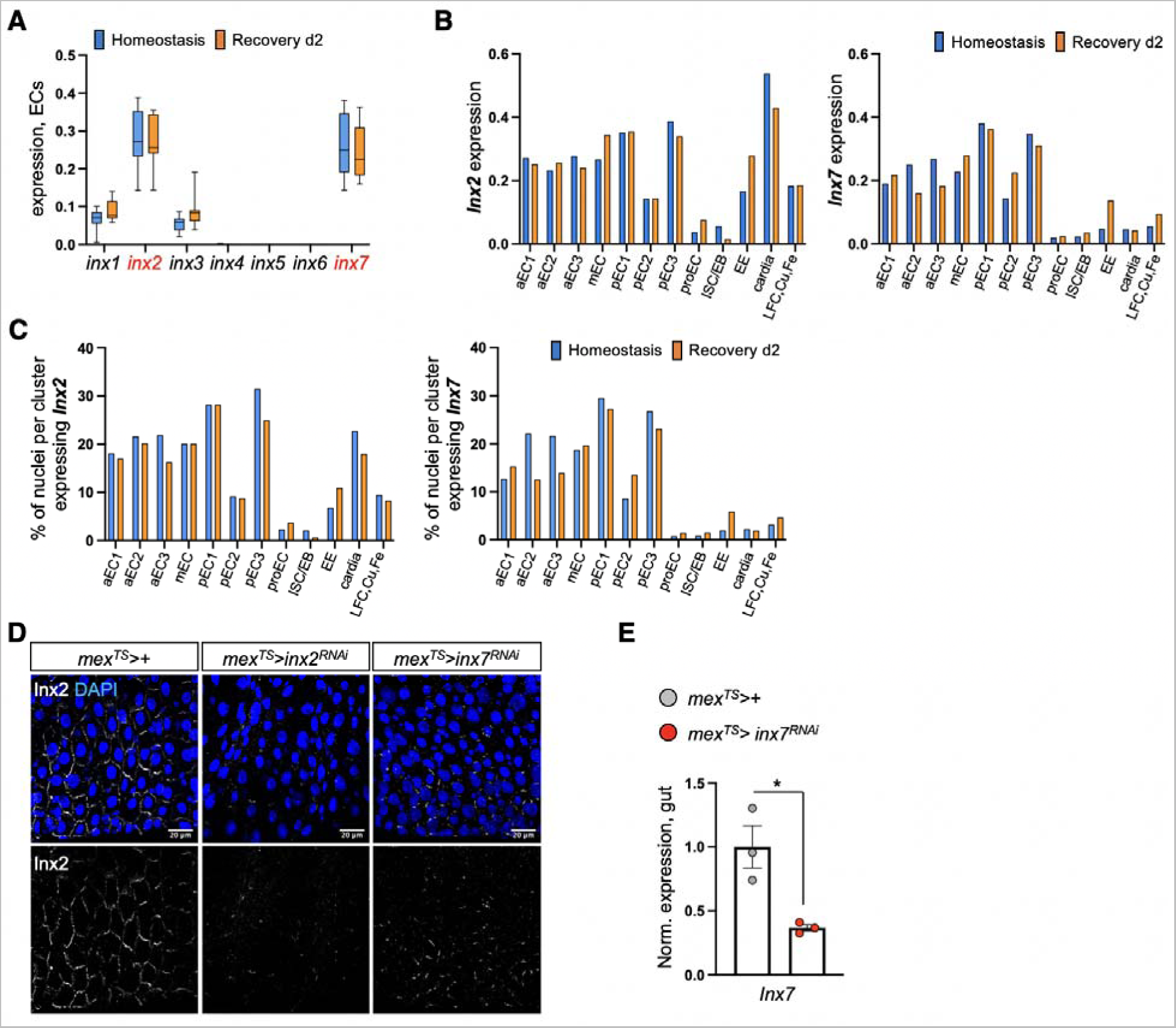
nAChR-mediated Ca^2+^ propagates via Inx2/Inx7 gap junctions. **(A)** Boxplot illustrating the mean expression levels of *Drosophila* innexins in EC clusters during Homeostasis (blue) and Recovery d2 (orange). **(B)** *inx2* and *inx7* mean expression levels in all gut clusters during Homeostasis (blue) and Recovery d2 (orange). **(C)** Percentage of nuclei expressing *inx7 and inx2* per cluster and condition in the gut. **(D)** Confocal images from posterior midgut of control flies (*mex>+*) and flies with conditional reduction of *inx2* (*mex^TS^>inx2^RNAi^*) and *inx7* (*mex^TS^>inx7^RNAi^*) for 2 days during Homeostasis. Inx2-gap junctions were visualized with anti-Inx2 (grey). DAPI: blue (nuclei). **(E)** Validation of *inx7* RNAi knock down. p-value was obtained with t-test; *: 0.05>p>0.01.

## Supplementary Information

### Supplementary Tables

**S. Table 1.** Top 20 upregulated genes in EC clusters during Homeostasis. Average expression, fold change and p-value of the 20 most significantly upregulated genes in EC clusters during Homeostasis as shown in Figure 1E.

| S. Table 1: Top 20 upregulated genes in EC clusters during Homeostasis |  |  |  |  |  |  |  |
| --- | --- | --- | --- | --- | --- | --- | --- |
| Feature ID | Feature Name | Homeostasis Average | Homeostasis Log2 Fold Change | Homeostasis P-Value | Recovery d2 Average | Recovery d2 Log2 Fold Change | Recovery d2 P-Value |
| FBgn0032889 | CG9331 | 2.226547128 | 1.289501629 | 6.07191E-05 | 0.910650724 | -1.289501629 | 6.07191E-05 |
| FBgn0259236 | comm3 | 1.228206322 | 1.228570472 | 0.000348801 | 0.523863811 | -1.228570472 | 0.000348801 |
| FBgn0038755 | Hs6st | 2.471985271 | 1.199742456 | 0.000507757 | 1.076005724 | -1.199742456 | 0.000507757 |
| FBgn0034808 | CG9896 | 2.06405015 | 1.169602911 | 0.000560433 | 0.917360782 | -1.169602911 | 0.000560433 |
| FBgn0262733 | Src64B | 1.770201449 | 1.084677475 | 0.001604504 | 0.834443637 | -1.084677475 | 0.001604504 |
| FBgn0038194 | Cyp6d5 | 1.245471626 | 1.043748644 | 0.002846527 | 0.603905217 | -1.043748644 | 0.002846527 |
| FBgn0036767 | CG16775 | 2.705574676 | 0.940550996 | 0.009212188 | 1.409591463 | -0.940550996 | 0.009212188 |
| FBgn0033188 | Drat | 1.270523243 | 0.936112546 | 0.011384258 | 0.663816449 | -0.936112546 | 0.011384258 |
| FBgn0026575 | hang | 2.10501293 | 0.924281577 | 0.012369538 | 1.109076724 | -0.924281577 | 0.012369538 |
| FBgn0031580 | CG15423 | 1.664578414 | 0.933537345 | 0.01266869 | 0.871348956 | -0.933537345 | 0.01266869 |
| FBgn0030098 | CG12057 | 1.875147414 | 0.920260944 | 0.024184345 | 0.99069213 | -0.920260944 | 0.024184345 |
| FBgn0031011 | CG8034 | 2.73231897 | 0.839338308 | 0.028253279 | 1.527017478 | -0.839338308 | 0.028253279 |
| FBgn0085358 | Diedel3 | 5.826532251 | 0.834393808 | 0.02918147 | 3.267798231 | -0.834393808 | 0.02918147 |
| **FBgn0000024** | **Ace** | **1.063678132** | **0.827671636** | **0.036466398** | **0.599112319** | **-0.827671636** | **0.036466398** |
| FBgn0043364 | cbt | 1.549814923 | 0.804559619 | 0.037446637 | 0.887165522 | -0.804559619 | 0.037446637 |
| FBgn0001145 | Gs2 | 1.901214637 | 0.82620092 | 0.040936694 | 1.072171406 | -0.82620092 | 0.040936694 |
| FBgn0033853 | CG6145 | 1.379531632 | 0.790243027 | 0.042427252 | 0.797538319 | -0.790243027 | 0.042427252 |
| FBgn0051431 | CG31431 | 1.003080301 | 0.794730924 | 0.043818809 | 0.578023565 | -0.794730924 | 0.043818809 |
| FBgn0025574 | Pli | 1.99532747 | 0.728692295 | 0.079581142 | 1.203976116 | -0.728692295 | 0.079581142 |
| FBgn0003748 | Treh | 1.237008242 | 0.722319902 | 0.084101708 | 0.749609333 | -0.722319902 | 0.084101708 |

**S. Table 2.** Top 20 upregulated genes in EC clusters during Recovery d2. Average expression, fold change and p-value of the 20 most significantly upregulated genes in EC clusters during Recovery d2 as shown in Figure 1E.

| Feature ID | Feature Name | Homeostasis Average | Homeostasis Log2 Fold Change | Homeostasis P-Value | Recovery d2 Average | Recovery d2 Log2 Fold Change | Recovery d2 P-Value |
| --- | --- | --- | --- | --- | --- | --- | --- |
| FBgn0267910 | lncRNA:CR34335 | 15.237477 | -2.112901065 | 2.67957E-15 | 65.92152665 | 2.112901065 | 2.67957E-15 |
| FBgn0263005 | CG43313 | 0.888655346 | -1.780268198 | 1.53628E-10 | 3.053555666 | 1.780268198 | 1.53628E-10 |
| FBgn0034709 | Swim | 0.716002307 | -1.498659054 | 5.29143E-06 | 2.024041058 | 1.498659054 | 5.29143E-06 |
| FBgn0265991 | Zasp52 | 1.879209838 | -1.361805013 | 8.56648E-06 | 4.830762448 | 1.361805013 | 8.56648E-06 |
| FBgn0041184 | Socs36E | 1.573173864 | -1.307146591 | 1.52612E-05 | 3.893750782 | 1.307146591 | 1.52612E-05 |
| FBgn0033448 | hebe | 1.138155913 | -1.205815961 | 8.82001E-05 | 2.626029115 | 1.205815961 | 8.82001E-05 |
| FBgn0052982 | CG32982 | 1.54270568 | -1.193673448 | 0.00018319 | 3.529490492 | 1.193673448 | 0.00018319 |
| FBgn0040687 | CG14645 | 1.477368354 | -1.138606279 | 0.000277664 | 3.253419535 | 1.138606279 | 0.000277664 |
| FBgn0004456 | mew | 1.16253046 | -1.149757158 | 0.000342608 | 2.580017289 | 1.149757158 | 0.000342608 |
| FBgn0034289 | CG10910 | 2.298655161 | -1.134609312 | 0.000691557 | 5.047880752 | 1.134609312 | 0.000691557 |
| FBgn0085249 | CG34220 | 0.597853463 | -1.060262119 | 0.001593676 | 1.247112203 | 1.060262119 | 0.001593676 |
| FBgn0085353 | CG34324 | 0.783709381 | -1.03348397 | 0.002067007 | 1.604662434 | 1.03348397 | 0.002067007 |
| FBgn0029762 | NAAT1 | 0.733606147 | -1.065348447 | 0.00386101 | 1.535644695 | 1.065348447 | 0.00386101 |
| FBgn0027932 | Akap200 | 0.85615595 | -0.97084245 | 0.00386101 | 1.678473072 | 0.97084245 | 0.00386101 |
| FBgn0001186 | Hex-A | 0.951284389 | -0.951035954 | 0.004391881 | 1.839514463 | 0.951035954 | 0.004391881 |
| FBgn0001104 | Galphai | 1.117505256 | -0.859047413 | 0.012938004 | 2.027396086 | 0.859047413 | 0.012938004 |
| FBgn0011661 | Moe | 5.50119976 | -0.821700318 | 0.01903966 | 9.724791157 | 0.821700318 | 0.01903966 |
| FBgn0031170 | ABCA | 0.574494523 | -0.853251542 | 0.023203517 | 1.038141826 | 0.853251542 | 0.023203517 |
| FBgn0267698 | Pak | 1.599579623 | -0.804512541 | 0.023988656 | 2.794259854 | 0.804512541 | 0.023988656 |
| FBgn0027556 | CG4928 | 2.73231897 | -0.852334597 | 0.025952859 | 4.933809767 | 0.852334597 | 0.025952859 |

### Supplementary Movie Legends

<u>Movies S1-S3:</u> Movies depicting cholinergic sensitivity assay described on Fig. 1I. **S1.** *mex^TS^ > GCaMP7c* midgut during Homeostasis. **S2.** *mex^TS^ > GCaMP7c* midgut during Recovery d2. **S3.** *mex^TS^ > GCaMP7c + dCas9VPR, gRNA-Ace* gut during Recovery d2.

<u>Movies S4-S6:</u> Movies depicting nicotinic sensitivity assay described on Fig. 2F-G. **S6.** *mex^TS^ > GCaMP7c* midgut during Homeostasis. **S7.** *mex^TS^ > GCaMP7c* midgut during Recovery d2. **S8.** *mex^TS^ > GCaMP7c + nAChRβ3^RNAi^*gut during Recovery d2.

<u>Movie S7:</u> Movie of R49E06 innervations at R4 in R49E06*^TS^> syt1HA + mexLexA>6xLexAopGFP* fly during Recovery d2 as described on Fig. 4D

<u>Movies S8-S9:</u> Movies depicting nicotine and heptanol assay during Recovery d2 as described on Fig. 5E. **S8** midgut from *mexLexA > LexAopGCaMP7c* flies administered Nicotine. **S9.** midgut from *mexLexA > LexAopGCaMP7c* flies administered nicotine and heptanol.

### Full Genotypes

<u>Fig. 1:</u>

*Ore* R corresponds to Oregon R

*Myo1A^TS^> dCas9VPR* corresponds to *+; Myo1A-Gal4 Tubulin-Gal80^TS^/+; UAS-3XFLAG-dCas9- VPR/+*

*Myo1A^TS^> dCas9VPR, gRNA-ACE* corresponds to *+; Myo1A-Gal4 Tubulin-Gal80^TS^/gRNA-ACE; UAS-3XFLAG-dCas9VPR/+*

*Mex^TS^>GCAMP7c* corresponds to *+; Mex-Gal4 Tubulin-Gal80^TS^/+; 20XUAS-IVS-jGCaMP7c/+*

*Mex^TS^>GCAMP7c + dCas9VPR, gRNA-ACE* corresponds to *+; Mex-Gal4 Tubulin-*

*Gal80^TS^/gRNA-ACE; 20XUAS-IVS-jGCaMP7c/UAS-3XFLAG-dCas9-VPR*

<u>Fig. 2:</u>

*Myo1ATS* > + corresponds to +; Myo1A-Gal4 Tubulin-Gal80^TS^/+; UAS- Luciferase^RNAi^/+

*Myo1A^TS^ > nAChRβ3^RNAi^* corresponds to *+; Myo1A-Gal4 Tubulin-Gal80^TS^/+; UAS-nAChRβ3^RNAi^/+*

*Mex^TS^ > +* corresponds to *+; Mex-Gal4 Tubulin-Gal80^TS^/UAS- emptyVK37; +*

*Mex^TS^ > nAChRβ3^RNAi^* corresponds to *+; Mex-Gal4 Tubulin-Gal80^TS^/+; UAS-nAChRβ3^RNAi^/+*

*Mex^TS^ > + / esg-GFP* corresponds to *+; Mex-Gal4 Tubulin-Gal80^TS^/esg-GFP; +*

*Mex^TS^ > nAChRβ3^RNAi^/ esg-GFP* corresponds to *+; Mex-Gal4 Tubulin-Gal80^TS^/esg-GFP; UAS- nAChRβ3^RNAi^/+*

*Mex > GFP / nAChRβ3-flag* corresponds to *+; Mex-Gal4 UAS-2x-GFP/nAChRβ3-flag; +*

*Myo1A > GFP / nAChRβ3-flag* corresponds to +; *Myo1A-Gal4 UAS-GFP/nAChRβ3-flag; +*

*Mex^TS^>GCAMP7c* corresponds to *+; Mex-Gal4 Tubulin-Gal80^TS^/+; 20XUAS-IVS-jGCaMP7c/+*

*Mex^TS^>GCAMP7c + nAcRβ3^RNAi^* corresponds to *+; Mex-Gal4 Tubulin-Gal80^TS^/+; 20XUAS-IVS- jGCaMP7c/UAS-nAcRβ3^RNAi^*

<u>Fig. 3:</u>

*Myo1ATS > +* corresponds to *+; Myo1A-Gal4 Tubulin-Gal80^TS^; UAS-Luciferase^RNAi^* and *+; Myo1- Gal4 Tubulin-Gal80^TS^/UAS-2x-GFP; +*

*Myo1A^TS^ > nAChRβ3^RNAi^* corresponds to *+; Myo1A-Gal4 Tubulin-Gal80^TS^/+; UAS-nAChRβ3^RNAi^/+*

*Mex^TS^ > +* corresponds to *+; Mex-Gal4 Tubulin-Gal80^TS^/UAS- emptyVK37; +*

*Mex^TS^ > nAChRβ3^RNAi^* corresponds to *+; Mex-Gal4 Tubulin-Gal80^TS^/+; UAS-nAChRβ3^RNAi^/+*

*Myo1A^TS^ > NFAT-CaLexA* corresponds to *+; Myo1A-Gal4 Tubulin-Gal80^TS^/LexAop-CD8-GFP- 2A-CD8GFP; UAS-mLexA-VP16-NFAT LexAop-rCD2-GFP/+*

*Mex^TS^ > NFAT-CaLexA* corresponds to *+; Mex-Gal4 Tubulin-Gal80^TS^/LexAop-CD8-GFP-2A- CD8GFP; UAS-mLexA-VP16-NFAT LexAop-rCD2-GFP/ +*

*Mex^TS^ > NFAT-CaLexA + nAChRβ3^RNAi^* corresponds to *+; Mex-Gal4 Tubulin-Gal80^TS^/LexAop- CD8-GFP-2A-CD8GFP; UAS-mLexA-VP16-NFAT LexAop-rCD2-GFP/UAS-nAcRβ3^RNAi^*

*Myo1A^TS^ > +* corresponds to *+; Myo1A-Gal4 Tubulin-Gal80^TS^/UAS-emptyVK37; +/+*

*Myo1A^TS^ > nAChRβ3^RNAi-2^* corresponds to *+; Myo1A-Gal4 Tubulin-Gal80^TS^/UAS-nAChRβ3^RNAi-2^; UAS-Luciferase/+*

*Myo1A^TS^ > Orai* corresponds to *+; Myo1A-Gal4 Tubulin-Gal80^TS^/UAS-emptyVK37; UAS-Orai / + Myo1A^TS^ > nAChRβ3^RNAi-2^* + Orai corresponds to *+;*

*Myo1A-Gal4 Tubulin-Gal80^TS^/UAS- nAChRβ3^RNAi-2^; UAS-Orai /+*

*Mex^TS^ > nAChRβ3* corresponds to *+; Mex-Gal4 Tubulin-Gal80^TS^/UAS-nAcRβ3; +*

*Myo1A^TS^ > nAChRβ3* corresponds to *+; Myo1A-Gal4 Tubulin-Gal80^TS^/UAS-nAcRβ3; +*

<u>Fig. 4:</u>

*R49E06 > 6xmCherry* corresponds to +; +; *R49E06-Gal4/20xUAS-6xmCherry-HA*

*R49E06^TS^> myrGFP+syt1HA* corresponds to +; *Tubulin-Gal80^TS^/10xUAS-IVS-myr::GFP;*

*R49E06-Gal4/5xUAS-IVS-Syt1::smGdP-HA*

*R49E06^TS^> syt1HA + mexLexA > LexAop6xGFP* corresponds to +; *Tubulin-Gal80^TS^/*

*13xLexAop2-6xGFP; R49E06-Gal4 mex-LexA::GAD/ 5xUAS-IVS-Syt1::smGdP-HA*

*R49E06^TS^> +* corresponds to +; *Tubulin-Gal80^TS^/+; R49E06-Gal4; UAS-Luciferase^RNAi^*

*R49E06^TS^> ChAT^RNAi^* corresponds to *+; Tubulin-Gal80^TS^/+; R49E06-Gal4; UAS-ChAT^RNAi^*

*ARCENs* + *mexLexA^TS^ > +* corresponds to *+; Tubulin-Gal80^TS^/ UAS-emptyVK37; R49E06-Gal4 mex-LexA::GAD / +*

*ARCENs> shibire^TS^* corresponds to *+; +; R49E06-Gal4/UAS-shibire^TS^*

*mexLexA^TS^ > LexAopnAChRβ3* corresponds to *+; Tubulin-Gal80^TS^/LexAop-nAChRβ3*; *mex- LexA::GAD/+*

*ARCENs* > *shibire^TS^* + *mexLexA^TS^> LexAopnAChRβ3* corresponds to *+; Tubulin-Gal80^TS^/ LexAop-nAChRβ3*; *R49E06-Gal4 mex-LexA::GAD / UAS-shibire^TS^*

*QUASCsChrimson + Mex ^TS^ > NFAT-CaLexA* corresponds to *QUASCsChrimson/+; Mex-Gal4*

*Tubulin-Gal80^TS^/LexAop-CD8-GFP-2A-CD8GFP; UAS-mLexA-VP16-NFAT LexAop-rCD2- GFP/+*

*ARCENsQF > QUASCsChrimson + Mex^TS^ > NFAT-CaLexA* corresponds to *QUASCsChrimson/+; Mex-Gal4 Tubulin-Gal80^TS^/LexAop-CD8-GFP-2A-CD8GFP; UAS-mLexA- VP16-NFAT LexAop-rCD2-GFP/R49E06-QF*

<u>Fig. 5:</u>

*MexTS > NFAT-CaLexA* corresponds to *+; Mex-Gal4 Tubulin-Gal80^TS^/ LexAop-CD8-GFP-2A- CD8GFP; UAS-mLexA-VP16-NFAT LexAop-rCD2-GFP/ +*

*Mex^TS^ > NFAT-CaLexA + nAChRβ3^RNAi^* corresponds to *+; Mex-Gal4 Tubulin-Gal80^TS^/ LexAop- CD8-GFP-2A-CD8GFP; UAS-mLexA-VP16-NFAT LexAop-rCD2-GFP/ UAS-nAChRβ3^RNAi^*

*Mex^TS^ > NFAT-CaLexA + inx2^RNAi^* corresponds to *+; Mex-Gal4 Tubulin-Gal80^TS^/ LexAop-CD8- GFP-2A-CD8GFP; UAS-mLexA-VP16-NFAT LexAop-rCD2-GFP/UAS-inx2^RNAi^*

*Mex^TS^ > NFAT-CaLexA + inx7^RNAi^* corresponds to *+; Mex-Gal4 Tubulin-Gal80^TS^/ LexAop-CD8- GFP-2A-CD8GFP; UAS-mLexA-VP16-NFAT LexAop-rCD2-GFP/UAS-inx7^RNAi^*

*Mex^TS^ > +* corresponds to *+; Mex-Gal4 Tubulin-Gal80^TS^/UAS- emptyVK37; +* and *+; Mex-Gal4 Tubulin-Gal80^TS^/* +*; UAS-Luciferase^RNAi^/+*

*Mex^TS^ > inx2^RNAi^* corresponds to *+; Mex-Gal4 Tubulin-Gal80^TS^; UAS-inx2^RNAi^/+* and *+; Mex-Gal4 Tubulin-Gal80^TS^/UAS-emptyVK37; UAS-inx2^RNAi^/+*

*Mex^TS^ > inx2^RNAi^* + *nAChRβ3* corresponds to *+; Mex-Gal4 Tubulin-Gal80^TS^/ UAS-nAChRβ3; UAS-inx2^RNAi^/+*

*Mex^TS^ > nAChRβ3* corresponds to *+; Mex-Gal4 Tubulin-Gal80^TS^/* UAS-nAChR*β*3*;UAS- Luciferase^RNAi^/+*

*MexLexA > LexAopGCAMP7c* corresponds to *+; Mex-LexA::GAD/+; 13xLexAop-IVS- jGCaMP7c/+*

<u>Ext. Fig. 1:</u>

*Ore R* corresponds to *Oregon R*

*Myo1A^TS^> dCas9-VPR* corresponds to *+; Myo1A-Gal4 Tubulin-Gal80^TS^ /+; UAS-3XFLAG-dCas9-VPR/+*

*Myo1A^TS^> dCas9-VPR, gRNA-ACE* corresponds to *+; Myo1A-Gal4 Tubulin-Gal80^TS^ / gRNA- ACE; UAS-3XFLAG-dCas9-VPR/+*

*Mex^TS^> dCas9-VPR* corresponds to *+; Mex-Gal4 Tubulin-Gal80^TS^/+ ; UAS-3XFLAG-dCas9-VPR/+*

*Mex^TS^> dCas9-VPR, gRNA-ACE* corresponds to *+; Mex-Gal4 Tubulin-Gal80^TS^ / gRNA-ACE; UAS-3XFLAG-dCas9-VPR/+*

*How^TS^> dCas9-VPR* corresponds to *+; Tubulin-Gal80^TS^ /+; how^24B^-Gal4/ UAS-3XFLAG-dCas9-VPR*

*How^TS^> dCas9-VPR, gRNA-ACE* corresponds to *+; Tubulin-Gal80^TS^ / gRNA-ACE; how^24B^-Gal4/ UAS-3XFLAG-dCas9-VPR*

*Hml^TS^ > dCas9-VPR* corresponds to *+; hml-Gal4Δ UAS-GFP/+; Tubulin-Gal80^TS^/ UAS-3XFLAG-dCas9-VPR*

*Hml^TS^ > dCas9-VPR, gRNA-ACE* corresponds to *+; hml-Gal4Δ UAS-GFP// gRNA-ACE; Tubulin- Gal80^TS^/ UAS-3XFLAG-dCas9-VPR*

<u>Ext. Fig. 2:</u>

*Myo1A^TS^ > +* corresponds to *+; Myo1A-Gal4 Tubulin-Gal80^TS^/+; +*

*Myo1A^TS^ > nAChRβ3^RNAi^* corresponds to *+; Myo1A-Gal4 Tubulin-Gal80^TS^/+; UAS-nAChRβ3^RNAi^/+*

*Myo1A^TS^ > nAChRβ3^RNAi-2^* corresponds to *+; Myo1A-Gal4 Tubulin-Gal80^TS^/+; UAS-nAChRβ3^RNAi-2^/+*

*Mex^TS^> GCAMP7c* corresponds to *+; Mex-Gal4 Tubulin-Gal80^TS^/+; 20XUAS-IVS-jGCaMP7c/+*

*Mex^TS^> GCAMP7c / nAChRβ3^RNAi^* corresponds to *+; Mex-Gal4 Tubulin-Gal80^TS^/+; 20XUAS-IVS-jGCaMP7c/ UAS-nAChRβ3^RNAi^*

*Esg^TS^ > +* corresponds to *+; esg-Gal4 Tubulin-Gal80^TS^ / +; UAS-Luciferase^RNAi^/+*

*Esg^TS^ > nAChRβ3^RNAi^*corresponds to *+; esg-Gal4 Tubulin-Gal80^TS^; UAS-nAChRβ3^RNAi^ /+*

*Su(H)GBE^TS^ > +* corresponds to *+; Su(H)Gbe-Gal4 UAS-CD8-GFP/+; Tubulin-Gal80^TS^/UAS-Luciferase^RNAi^*

*Su(H)GBE ^TS^ > nAChRβ3^RNAi^* corresponds to *+; Su(H)Gbe-Gal4 UAS-CD8-GFP/+; Tubulin-Gal80^TS^/ UAS-nAChRβ3^RNAi^*

*Pros^TS^ > +* corresponds to *+; Tubulin-Gal80^TS^ / +; prospero-Gal4/ UAS-Luciferase^RNAi^*

*Pros^TS^ > nAChRβ3^RNAi^* corresponds to *+; Tubulin-Gal80^TS^ / +; prospero-Gal4/ UAS-nAChRβ3^RNAi^*

*How^TS^ > +* corresponds to +; *Tubulin-Gal80^TS^ / +; how^24B^-Gal4/ UAS-Luciferase ^RNAi^*

*How^TS^ > nAChRβ3^RNAi^* corresponds to +; *Tubulin-Gal80^TS^ / +; how^24B^-Gal4/ UAS-nAChRβ3^RNAi^*

*Hml^TS^ > +* corresponds to *+; hml-Gal4Δ UAS-GFP/+; Tubulin-Gal80^TS^/ UAS-Lucifersa^RNAi^*

*Hml^TS^ > nAChRβ3^RNAi^* corresponds to *+; hml-Gal4Δ UAS-GFP/+; Tubulin-Gal80^TS^/ UAS-nAChRβ3^RNAi^*

*Mex^TS^ > + / nAChRβ3-flag* corresponds to *+; Mex-Gal4 Tubulin-Gal80^TS^ / nAChRβ3-flag; +*

*Mex^TS^ > UAS-nAChRβ3^RNAi^ / nAChRβ3-flag* corresponds to *+; Mex-Gal4 Tubulin-Gal80^TS^ / nAChRβ3-flag; UAS-nAChRβ3^RNAi^ /+*

<u>Ext. Fig. 3:</u>

*Myo1A^TS^ > +* corresponds to *+; Myo1A-Gal4 Tubulin-Gal80^TS^/UAS-emptyVK37; +/+* and *+;*

*Myo1-Gal4 Tubulin-Gal80^TS^/UAS-2x-GFP; +*

*Myo1A^TS^ > nAChRβ3^RNAi^* corresponds to *+; Myo1A-Gal4 Tubulin-Gal80^TS^/+; UAS-nAChRβ3^RNAi^/+*

*Myo1A^TS^ > + / Diap1-LacZ* corresponds to *+; Myo1A-Gal4 Tubulin-Gal80^TS^/+; Diap1-LacZ / +*

*Myo1A^TS^ > nAChRβ3^RNAi^* corresponds to *+; Myo1A-Gal4 Tubulin-Gal80^TS^/+; Diap1-LacZ /UAS-nAChRβ3^RNAi^*

*Mex^TS^ > +* corresponds to *+; Mex-Gal4 Tubulin-Gal80^TS^/UAS-emptyVK37; +*

*Mex^TS^ > nAChRβ3^RNAi^* corresponds to *+; Mex-Gal4 Tubulin-Gal80^TS^/+; UAS-nAChRβ3^RNAi^/+*

*Esg^TS^ > NFAT-CaLexA* corresponds to *Tubulin-Gal80^TS^/+*; *esg-Gal4 / LexAop-CD8-GFP-2A-CD8GFP; UAS-mLexA-VP16-NFAT LexAop-rCD2-GFP/ +*

*Myo1A^TS^ > nAChRβ3^RNAi-2^* corresponds to *+; Myo1A-Gal4 Tubulin-Gal80^TS^/UAS-nAChRβ3^RNAi-2^; UAS-Luciferase/+*

*Myo1A^TS^> Orai* corresponds to *+; Myo1A-Gal4 Tubulin-Gal80^TS^/UAS-emptyVK37; UAS-Orai/+*

*Myo1A^TS^> nAChRβ3^RNAi-2^* + Orai corresponds to *+; Myo1A-Gal4 Tubulin-Gal80^TS^/UAS-nAChRβ3^RNAi-2^; UAS-Orai /+*

*Myo1A^TS^> nAChRβ3* corresponds to *+; Myo1A-Gal4 Tubulin-Gal80^TS^/ UAS-nAChRβ3; +*

*Mex^TS^> GCAMP7c* corresponds to *+; Mex-Gal4 Tubulin-Gal80^TS^/+; 20XUAS-IVS-jGCaMP7c/+*

*Mex^TS^> GCAMP7c + nAChRβ3* corresponds to *+;Mex-Gal4 Tubulin-Gal80^TS^/ nAChRβ3; 20XUAS-IVS-jGCaMP7c/ +*

<u>Ext. Fig. 4:</u>

*Elav^TS^ > +* corresponds to *Elav-Gal4/+; Tubulin-Gal80^TS^/+; UAS-Luciferase^RNAi^/+*

*Elav^TS^ > ChAT^RNAi^* corresponds to *Elav-Gal4/+; Tubulin-Gal80^TS^/+; UAS-ChAT^RNAi^ /+*

*Esg^TS^ > +* corresponds to *Tubulin-Gal80^TS^/+; esg-Gal4 / +; UAS-Luciferase ^RNAi^/+*

*Esg^TS^ > ChAT^RNAi^* corresponds to *Tubulin-Gal80^TS^/+; esg-Gal4 / +; UAS-ChAT^RNAi^ /+*

*Pros^TS^ > +* corresponds to +; *Tubulin-Gal80^TS^ / +; prospero-Gal4/ UAS-Luciferase^RNAi^*

*Pros^TS^ > ChAT^RNAi^* corresponds to +; *Tubulin-Gal80^TS^ / +; prospero-Gal4/ UAS-ChAT^RNAi^*

*Myo1A^TS^ > +* corresponds to +; *Tubulin-Gal80^TS^, myo1A-Gal4 /+ ; UAS-Luciferase ^RNAi^/+*

*Myo1A^TS^ > ChAT^RNAi^* corresponds to +; *Tubulin-Gal80^TS^, myo1A-Gal4 /+; UAS-ChAT^RNAi^/+*

*Hml^TS^ > +* corresponds to *+; hml-Gal4Δ UAS-GFP/+; Tubulin-Gal80^TS^/ UAS-Lucifersa^RNAi^*

*Hml^TS^ > ChAT^RNAi^* corresponds to *+; hml-Gal4Δ UAS-GFP/+; Tubulin-Gal80^TS^/ UAS-ChAT^RNAi^*

*How^TS^ > +* corresponds to +; *Tubulin-Gal80^TS^ / +; how^24B^-Gal4/ UAS-Luciferase ^RNAi^*

*How^TS^ > ChAT^RNAi^* corresponds to +; *Tubulin-Gal80^TS^ / +; how^24B^-Gal4/ UAS-ChAT^RNAi^*

*Esg > +* corresponds to +; *esg-Gal4 / UAS-emptyVK37; +*

*Esg > sc^RNAi^* corresponds to +; *esg-Gal4/+; sc^RNAi^ /+*

*ChAT > mCD8GFP* corresponds to *+; 10xUAS-mCD8::GFP/+ ; ChAT^MI04508^-Gal4/ +* and *+; +;*

*ChAT^MI04508^-Gal4/ 10xUAS-mCD8::GFP*

*ChAT > 6XGFP* corresponds to *+; +; ChAT^MI04508^-Gal4/ 20XUAS-6XGFP*

*ChAT-QF> QUAS-TomatoHA+myo1A > GFP* corresponds to *UAS-mCD8::GFP, QUAS-mtdTomato-3xHA/+; Myo1A-Gal4, UAS-GFP/ + ; ChAT-QF/+*

*ChAT> mCD8GFP/ Tsh-Gal80* corresponds to *+; 10xUAS-mCD8::GFP/ Tsh-Gal80 ;*

*ChAT^MI04508^-Gal4/ +*

*R49E06> mCD8GFP* corresponds to *+; 10xUAS-mCD8::GFP/+; R49E06-Gal4/+*

*R49E06> 2xGFP* corresponds to *+; UAS-2x-GFP/+; R49E06-Gal4/+*

<u>Ext. Fig. 5:</u>

*R49E06^TS^ > syt1HA* corresponds to *+; Tubulin-Gal80^TS^/* +; *R49E06-Gal4/5xUAS-IVS- Syt1::smGdP-HA*

*R49E06-QF> QUAS-TomatoHA* corresponds to *+; +; R49E06-QF/ QUAS-mtdTomato-3xHA*

*mexLexA> LexAopGFP* corresponds to *+; 13xLexAop2-sfGFP/+; mex-LexA::GAD/ +*

*mexLexA^TS^> LexAopnAChRβ3* corresponds to *+; Tubulin-Gal80^TS^/ LexAop-nAChRβ3*; *mex-LexA::GAD / +*

*R49E06^TS^> syt1HA + mexLexA > LexAop6xGFP* corresponds to +; *Tubulin-Gal80^TS^/*

*13xLexAop2-6xGFP; R49E06-Gal4 mex-LexA::GAD/ 5xUAS-IVS-Syt1::smGdP-HA*

*R49E06^TS^> syt1HA + mexLexA > LexAopxGFP* corresponds to +; *Tubulin-Gal80^TS^/*

*13xLexAop2-sfGFP; R49E06-Gal4 mex-LexA::GAD/ 5xUAS-IVS-Syt1::smGdP-HA*

*R49E06^TS^> +* corresponds to *+; Tubulin-Gal80^TS^/+; R49E06-Gal4; UAS-Luciferase^RNAi^*

*R49E06^TS^> ChAT^RNAi^* corresponds to *+; Tubulin-Gal80^TS^/+; R49E06-Gal4; UAS-ChAT^RNAi^*

*ARCENs>+* corresponds to *+; UAS-2xGFP/+; R49E06-Gal4/+*

*ARCENs> TrpA1* corresponds to *+; UAS-TrpA1/+; R49E06-Gal4/+*

*ARCENs> TrpA1/ChATGal80* corresponds to *+; UAS-TrpA1/+; R49E06-Gal4/ChAT-Gal80*

*ARCENs^TS^> +* corresponds to *+; Tubulin-Gal80^TS^/+; R49E06-Gal4; UAS-Luciferase^RNAi^*

*ARCENs^TS^> wgn^RNAi^* corresponds to *+; Tubulin-Gal80^TS^/UAS-wgn^RNAi^; R49E06-Gal4/+*

*ARCENs^TS^> grnd^RNAi^* corresponds to *+; Tubulin-Gal80^TS^/UAS-grnd^RNAi^; R49E06-Gal4/+*

<u>Ext. Fig. 6:</u>

*Mex^TS^ > +* corresponds to *+; Mex-Gal4 Tubulin-Gal80^TS^/UAS-emptyVK37; +*

*Mex^TS^ > inx2^RNAi^* corresponds to *+; Mex-Gal4 Tubulin-Gal80^TS^; UAS-inx2^RNAi^/+*

*Mex^TS^ > inx7^RNAi^* corresponds to *+; Mex-Gal4 Tubulin-Gal80^TS^; UAS-inx7^RNAi^/+*

### Supplementary Txt

#### gBlock1

AACGAGGATTATCATCAAAAGAGCGCCGGAGTATAAGTAGAGGCGCTTCGTCTACGGAGCGACAATTCAATTCAAACAAGCAAAGTGAACACGTCGCTAAGCGAAAGCTAAGCAAATAAACAAGCGCAGCTGAACAAGCTAAACAATCTGCAGTAAAGTGCAAGTTAAAGTGAATCAATTAAAAGTAACCAGCAACCAAGTAAATCAACTGCAACTACTGAAATCTGCCAAGAAGTAATTATTGAATACAAGAAGAGAACTCTGAATACTTTCAACAAGTTACCGAGAAAGAAGAACTCACACACAGCGGCCAATTCGGTACCGCGGCCGCTAAGCAA

#### gBlock2

ATATTTTTTATATACATACTTTTCAAATCGCGCGCCCTCTTCATAATTCACCTCCACCACACCACGTTTCGTAGTTGCTCTTTCGCTGTCTCCCACCCGCTCTCCGCAACACATTCACCTTTTGTTCGACGACCTTGGAGCGACTGTCGTTAGTTCCGCGCGATTCGGTTCGCTCAAATGGTTCCGAGTGGTTCATTTCGTCTCAATAGAAATTAGTAATAAATATTTGTATGTACAATTTATTTGCTCCAATATATTTGTATATATTTCCCTCACAGCTATATTTATTCTAATTTAATATTATGACTTTTTAAGGTAATTTTTTGTGACCTGTTCGGAGTGATTAGCGTTACAATTTGAACTGAAAGTGACATCCAGTGTTTGTTCCTTGTGTAGATGCATCTCAAAAAAATGGTGGGCATAATAGTGTTGTTTATATATATCAAAAATAACAACTATAATAATAAGAATACATTTAATTTAGAAAATGCTTGGATTTCACTGGAACTGCTACACCGATCACTTAGGACCCATAATGCATCATTGGAGGTGAAGAAAATCTGCTGAATGTTCTTCGATGTGATATATATGATTTCGTTGCCCTCGGCTGTGATGTTGGCTAGTTTGGCACATCGTACGACAGCTCCGAATTTGGGGATAACATAGTACAATGTTCCGAAATTGTCGATGATCATGCTGCGACTGCTGCCCAGAAGACTGCCCAAATGTGTGAGTTTAATGCTATTGTTAATTGGTTTGCTTGCTAATTCGCTTTTGCCTTCAATTTCCAGTCTGTCCACCGTACTGTATAGATCACCATCCTGGTCGGAGAGTATGAGTTCCCCTTGAATTCCGAAGATGAAATCCACTGGCTTGATGGGAAACGATTGATTCATATTCTCGTACTTATTGCTCTCCAAGGATAGGCGGTGCCACGTCTGCTCCAGGATGTCGTAGGCTAGAATTTCTGGTACCTTTCCAAGGATGAAGTAAATATGGCGCTCGTGCTCGCATCCCGGACTCTTGGGTCCCATTTGGACAGTCAGAAAGTGGGTGTCATTAGCACCTATATGGTGTCCACAATCGATTCTCAATAGCTCCGCGTTGTTACGCATCAGATCGAAGACAAACAAACGGGGTGAGCACGTGCTGCCGGGAAATCCGATGTCCATGACCCACAGTCGACTCAGTGCATCCACCTGGGACCAGTGTGCCTGTTGTACCAGAGAGCAATCATCATGATCCCCCATGGAATGAACATCGATGTGGGGAAAGACGACAGTGGGCATGGGAAAATAGGAAACCGGCCATTGCGCCTCGATCAAAGTGGGCGCATCGTTGCTTTCTACATTTACGCTGAGAAACACTCGCGAAAAGTGCATCGATAGATGACGGACTTTGTAGCTGGATTCGTTTAAACCAGATTCGGATAACAGCCAGGCGTCCGTGGCCAGAATTAGACCAACAATGAGAAATCCCAAGCCGAGGACACCATGTGTGACCATCTTTAAACGCGAATTCAGACTGAGCTTAGGGCTAGAGAACTCTGCTTACGTTGTAAGTGCCCGTTCAAGAAGTCCACCAAAGTGTCTACATCAGCAGTTGGGATCGGGTGTCGCCGAATATATCGGCGACCTGCATTAATGGGGACCAGCATTGGAGTGTTTCTCTATTCACTTGTGGTCCCAAAAAAAAAAAGCTTACCAATTTCAATCGGAGAATATCGAAATCATGACACAGTTTCCTCTTAAAATTACCCAAAATTGGATTGTTATATTTTTCAAAATAAATCAAAGTTAAAACCAACAAAATGGTTGTTTTGTACCACCATAATAACCTATAATAAATAAATCATAATAAGGCTATTTTTGTCGAACTTTAGCGATATTGATAAGGTTCCTGTGAACTGAACCTACCTTAAGTGCTTAGTTTTAACATAACCTTAATGTTAACGAGTATTTCTTCGCGTAAAAGTACAAAAAAACCCTATCATATCTTTACAGCCATTATAAAGATTGCTTATAGATGACTGAT

TTCACTTGGGAATTGCCAAAAAGCTAAAGTGAACCCCCGAGGAATCAATTAATGTTGTTTAGCTTTCAGTTTTTAGTACATTCGTATTTTTTTAAGTATTAAAATTATTTAATCACACGAATAAACATTAAGAAATATTTTATTCTGGAAACTTTCTTCGTTATAGAGCATTTATCGTTGTAAAAAAAATAATTTCATTAGTTTACATTACATTTTCGTTTATATGTTTAGATTTTTTCTAATTAAATGATATTGTATAATTTATTGCAAAAATTATAATACTCACTTCAAATAATCTGAAACAAATCCACCCACTCCGAAAAATGTGGTAAGCTTTATTTTCTAACACCTCTCTTCTGATTTTATTGATTATAACTGGAATCTTTATCGGTTTCTCTCTTTTTCCATCAGTAGGTTGAGAGCTTTTCATACTTTTCACGCCGAAATGGGAAAAATGGCACTGCGCGTAGAGTCAGTTATCCATTTGAGTTGGCCAAGATCAGATCGAAATACGCTAAAATCATTTTCGGGAGCACAATCGAATACTCATACAACACACACTTCAAACGAGGATTATCATCAAAAGAGCGCCGG

## References

1 McLaughlin, K. A. & Levin, M. Bioelectric signaling in regeneration: Mechanisms of ionic controls of growth and form. Dev Biol 433, 177–189, doi:10.1016/j.ydbio.2017.08.032 (2018).

2 Tracey, K. J. The inflammatory reflex. Nature 420, 853–859, doi:10.1038/nature01321 (2002).

3 Karin, M. & Clevers, H. Reparative inflammation takes charge of tissue regeneration. Nature 529, 307–315, doi:10.1038/nature17039 (2016).

4 Tracy Cai, X., et al. AWD regulates timed activation of BMP signaling in intestinal stem cells to maintain tissue homeostasis. Nat Commun 10, 2988, doi:10.1038/s41467-019-10926-2 (2019).

5 Tracey, K. J. Reflex control of immunity. Nat Rev Immunol 9, 418–428, doi:10.1038/nri2566 (2009).

6 Levin, M. Bioelectric signaling: Reprogrammable circuits underlying embryogenesis, regeneration, and cancer. Cell 184, 1971–1989, doi:10.1016/j.cell.2021.02.034 (2021).

7 Guo, Z., Driver, I. & Ohlstein, B. Injury-induced BMP signaling negatively regulates Drosophila midgut homeostasis. J Cell Biol 201, 945–961, doi:10.1083/jcb.201302049 (2013).

8 Rieder, F., Brenmoehl, J., Leeb, S., Scholmerich, J. & Rogler, G. Wound healing and fibrosis in intestinal disease. Gut 56, 130–139, doi:10.1136/gut.2006.090456 (2007).

9 Coskun, M. Intestinal epithelium in inflammatory bowel disease. Front Med (Lausanne) 1, 24, doi:10.3389/fmed.2014.00024 (2014).

10 Kaur, A. & Goggolidou, P. Ulcerative colitis: understanding its cellular pathology could provide insights into novel therapies. J Inflamm (Lond) 17, 15, doi:10.1186/s12950-020-00246-4 (2020).

11 Beaugerie, L. & Itzkowitz, S. H. Cancers Complicating Inflammatory Bowel Disease. N Engl J Med 373, 195, doi:10.1056/NEJMc1505689 (2015).

12 Kappelman, M. D., Moore, K. R., Allen, J. K. & Cook, S. F. Recent trends in the prevalence of Crohn’s disease and ulcerative colitis in a commercially insured US population. Dig Dis Sci 58, 519–525, doi:10.1007/s10620-012-2371-5 (2013).

13 Axelrad, J. E., Lichtiger, S. & Yajnik, V. Inflammatory bowel disease and cancer: The role of inflammation, immunosuppression, and cancer treatment. World J Gastroenterol 22, 4794–4801, doi:10.3748/wjg.v22.i20.4794 (2016).

14 Horiuchi, Y. et al. Evolutional study on acetylcholine expression. Life Sci 72, 1745–1756, doi:10.1016/s0024-3205(02)02478-5 (2003).

15 Wessler, I. & Kirkpatrick, C. J. Acetylcholine beyond neurons: the non-neuronal cholinergic system in humans. Br J Pharmacol 154, 1558–1571, doi:10.1038/bjp.2008.185 (2008).

16 Hirota, C. L. & McKay, D. M. Cholinergic regulation of epithelial ion transport in the mammalian intestine. Br J Pharmacol 149, 463–479, doi:10.1038/sj.bjp.0706889 (2006).

17 Brinkman, D. J., Ten Hove, A. S., Vervoordeldonk, M. J., Luyer, M. D. & de Jonge, W. J. Neuroimmune Interactions in the Gut and Their Significance for Intestinal Immunity. Cells 8, doi:10.3390/cells8070670 (2019).

18 Goverse, G., Stakenborg, M. & Matteoli, G. The intestinal cholinergic anti-inflammatory pathway. J Physiol 594, 5771–5780, doi:10.1113/JP271537 (2016).

19 Takahashi, T., Shiraishi, A. & Murata, J. The Coordinated Activities of nAChR and Wnt Signaling Regulate Intestinal Stem Cell Function in Mice. Int J Mol Sci 19, doi:10.3390/ijms19030738 (2018).

20 Takahashi, T. Multiple Roles for Cholinergic Signaling from the Perspective of Stem Cell Function. Int J Mol Sci 22, doi:10.3390/ijms22020666 (2021).

21 Miguel-Aliaga, I., Jasper, H. & Lemaitre, B. Anatomy and Physiology of the Digestive Tract of Drosophila melanogaster. Genetics 210, 357–396, doi:10.1534/genetics.118.300224 (2018).

22 Jiang, H., Tian, A. & Jiang, J. Intestinal stem cell response to injury: lessons from Drosophila. Cell Mol Life Sci 73, 3337–3349, doi:10.1007/s00018-016-2235-9 (2016).

23 Panayidou, S. & Apidianakis, Y. Regenerative inflammation: lessons from Drosophila intestinal epithelium in health and disease. Pathogens 2, 209–231, doi:10.3390/pathogens2020209 (2013).

24 Micchelli, C. A. & Perrimon, N. Evidence that stem cells reside in the adult Drosophila midgut epithelium. Nature 439, 475–479, doi:10.1038/nature04371 (2006).

25 Ohlstein, B. & Spradling, A. The adult Drosophila posterior midgut is maintained by pluripotent stem cells. Nature 439, 470–474, doi:10.1038/nature04333 (2006).

26 Biteau, B. & Jasper, H. EGF signaling regulates the proliferation of intestinal stem cells in Drosophila. Development 138, 1045–1055, doi:10.1242/dev.056671 (2011).

27 Buchon, N., Broderick, N. A., Kuraishi, T. & Lemaitre, B. Drosophila EGFR pathway coordinates stem cell proliferation and gut remodeling following infection. BMC Biol 8, 152, doi:10.1186/1741-7007-8-152 (2010).

28 Jiang, H. et al. Cytokine/Jak/Stat signaling mediates regeneration and homeostasis in the Drosophila midgut. Cell 137, 1343–1355, doi:10.1016/j.cell.2009.05.014 (2009).

29 Jiang, H., Grenley, M. O., Bravo, M. J., Blumhagen, R. Z. & Edgar, B. A. EGFR/Ras/MAPK signaling mediates adult midgut epithelial homeostasis and regeneration in Drosophila. Cell Stem Cell 8, 84–95, doi:10.1016/j.stem.2010.11.026 (2011).

30 Karpowicz, P., Perez, J. & Perrimon, N. The Hippo tumor suppressor pathway regulates intestinal stem cell regeneration. Development 137, 4135–4145, doi:10.1242/dev.060483 (2010).

31 Lin, G., Xu, N. & Xi, R. Paracrine Wingless signalling controls self-renewal of Drosophila intestinal stem cells. Nature 455, 1119–1123, doi:10.1038/nature07329 (2008).

32 Ren, F. et al. Hippo signaling regulates Drosophila intestine stem cell proliferation through multiple pathways. Proc Natl Acad Sci U S A 107, 21064–21069, doi:10.1073/pnas.1012759107 (2010).

33 Shaw, R. L. et al. The Hippo pathway regulates intestinal stem cell proliferation during Drosophila adult midgut regeneration. Development 137, 4147–4158, doi:10.1242/dev.052506 (2010).

34 Staley, B. K. & Irvine, K. D. Warts and Yorkie mediate intestinal regeneration by influencing stem cell proliferation. Curr Biol 20, 1580–1587, doi:10.1016/j.cub.2010.07.041 (2010).

35 Tian, A. & Jiang, J. Intestinal epithelium-derived BMP controls stem cell self-renewal in Drosophila adult midgut. Elife 3, e01857, doi:10.7554/eLife.01857 (2014).

36 Tamamouna, V. P., M.; Theophanous, A.; Demosthenous, M.; Michail, M.; Papadopoulou, M.; Teloni, S.; Pitsouli, C; Apidianakis, >Y. Evidence of two types of balance between stem cell mitosis and enterocyte nucleus growth in the Drosophila midgut. Development 147 (2020).

37 Chakrabarti, S. et al. Remote Control of Intestinal Stem Cell Activity by Haemocytes in Drosophila. PLoS Genet 12, e1006089, doi:10.1371/journal.pgen.1006089 (2016).

38 Ayyaz, A., Li, H. & Jasper, H. Haemocytes control stem cell activity in the Drosophila intestine. Nat Cell Biol 17, 736–748, doi:10.1038/ncb3174 (2015).

39 Tian, A., Wang, B. & Jiang, J. Injury-stimulated and self-restrained BMP signaling dynamically regulates stem cell pool size during Drosophila midgut regeneration. Proc Natl Acad Sci U S A 114, E2699–E2708, doi:10.1073/pnas.1617790114 (2017).

40 Subramanian, S., Geng, H. & Tan, X. D. Cell death of intestinal epithelial cells in intestinal diseases. Sheng Li Xue Bao 72, 308–324 (2020).

41 Sommer, J. et al. Interleukin-6, but not the interleukin-6 receptor plays a role in recovery from dextran sodium sulfate-induced colitis. Int J Mol Med 34, 651–660, doi:10.3892/ijmm.2014.1825 (2014).

42 Popivanova, B. K. et al. Blocking TNF-alpha in mice reduces colorectal carcinogenesis associated with chronic colitis. J Clin Invest 118, 560–570, doi:10.1172/JCI32453 (2008).

43 Agaisse, H., Petersen, U. M., Boutros, M., Mathey-Prevot, B. & Perrimon, N. Signaling role of hemocytes in Drosophila JAK/STAT-dependent response to septic injury. Dev Cell 5, 441–450, doi:10.1016/s1534-5807(03)00244-2 (2003).

44 Igaki, T. et al. Eiger, a TNF superfamily ligand that triggers the Drosophila JNK pathway. EMBO J 21, 3009–3018, doi:10.1093/emboj/cdf306 (2002).

45 Hung, R. J. et al. A cell atlas of the adult Drosophila midgut. Proc Natl Acad Sci U S A 117, 1514–1523, doi:10.1073/pnas.1916820117 (2020).

46 Buchon, N. et al. Morphological and molecular characterization of adult midgut compartmentalization in Drosophila. Cell Rep 3, 1725–1738, doi:10.1016/j.celrep.2013.04.001 (2013).

47 Wiesner, J., Kriz, Z., Kuca, K., Jun, D. & Koca, J. Acetylcholinesterases--the structural similarities and differences. J Enzyme Inhib Med Chem 22, 417–424, doi:10.1080/14756360701421294 (2007).

48 Miceli, P. C. & Jacobson, K. Cholinergic pathways modulate experimental dinitrobenzene sulfonic acid colitis in rats. Auton Neurosci 105, 16–24, doi:10.1016/S1566-0702(03)00023-7 (2003).

49 Russell, W. S., Henson, S. M., Hussein, A. S., Tippins, J. R. & Selkirk, M. E. Nippostrongylus brasiliensis: infection induces upregulation of acetylcholinesterase activity on rat intestinal epithelial cells. Exp Parasitol 96, 222–230, doi:10.1006/expr.2000.4565 (2000).

50 Hardy, S. P., Smith, P. M., Bayston, R. & Spitz, L. Electrogenic colonic ion transport in Hirschsprung’s disease: reduced secretion to the neural secretagogues acetylcholine and iloprost. Gut 34, 1405–1411, doi:10.1136/gut.34.10.1405 (1993).

51 Czarnewski, P. et al. Conserved transcriptomic profile between mouse and human colitis allows unsupervised patient stratification. Nat Commun 10, 2892, doi:10.1038/s41467-019-10769-x (2019).

52 Lin, S., Ewen-Campen, B., Ni, X., Housden, B. E. & Perrimon, N. In Vivo Transcriptional Activation Using CRISPR/Cas9 in Drosophila. Genetics 201, 433–442, doi:10.1534/genetics.115.181065 (2015).

53 Brand, A. H. & Perrimon, N. Targeted gene expression as a means of altering cell fates and generating dominant phenotypes. Development 118, 401–415 (1993).

54 McGuire, S. E., Mao, Z. & Davis, R. L. Spatiotemporal gene expression targeting with the TARGET and gene-switch systems in Drosophila. Sci STKE 2004, pl6, doi:10.1126/stke.2202004pl6 (2004).

55 Yang, X. et al. Molecular mechanisms of calcium signaling in the modulation of small intestinal ion transports and bicarbonate secretion. Oncotarget 9, 3727–3740, doi:10.18632/oncotarget.23197 (2018).

56 Gratz, S. J., Rubinstein, C. D., Harrison, M. M., Wildonger, J. & O’Connor-Giles, K. M. CRISPR-Cas9 Genome Editing in Drosophila. Curr Protoc Mol Biol 111, 31 32 31-31 32 20, doi:10.1002/0471142727.mb3102s111 (2015).

57 Lansdell, S. J. & Millar, N. S. Dbeta3, an atypical nicotinic acetylcholine receptor subunit from Drosophila : molecular cloning, heterologous expression and coassembly. J Neurochem 80, 1009–1018, doi:10.1046/j.0022-3042.2002.00789.x (2002).

58 Asano, S. M. et al. Expansion Microscopy: Protocols for Imaging Proteins and RNA in Cells and Tissues. Curr Protoc Cell Biol 80, e56, doi:10.1002/cpcb.56 (2018).

59 Li, M., Sun, S., Priest, J., Bi, X. & Fan, Y. Characterization of TNF-induced cell death in Drosophila reveals caspase- and JNK-dependent necrosis and its role in tumor suppression. Cell Death Dis 10, 613, doi:10.1038/s41419-019-1862-0 (2019).

60 Masuyama, K., Zhang, Y., Rao, Y. & Wang, J. W. Mapping neural circuits with activity-dependent nuclear import of a transcription factor. J Neurogenet 26, 89–102, doi:10.3109/01677063.2011.642910 (2012).

61 Deng, H., Gerencser, A. A. & Jasper, H. Signal integration by Ca(2+) regulates intestinal stem-cell activity. Nature 528, 212–217, doi:10.1038/nature16170 (2015).

62 Xu, C., Luo, J., He, L., Montell, C. & Perrimon, N. Oxidative stress induces stem cell proliferation via TRPA1/RyR-mediated Ca(2+) signaling in the Drosophila midgut. Elife 6, doi:10.7554/eLife.22441 (2017).

63 Amcheslavsky, A. et al. Enteroendocrine cells support intestinal stem-cell-mediated homeostasis in Drosophila. Cell Rep 9, 32–39, doi:10.1016/j.celrep.2014.08.052 (2014).

64 Kumar, A. & Brockes, J. P. Nerve dependence in tissue, organ, and appendage regeneration. Trends Neurosci 35, 691–699, doi:10.1016/j.tins.2012.08.003 (2012).

65 Farkas, J. E. & Monaghan, J. R. A brief history of the study of nerve dependent regeneration. Neurogenesis (Austin) 4, e1302216, doi:10.1080/23262133.2017.1302216 (2017).

66 Mahmoud, A. I. et al. Nerves Regulate Cardiomyocyte Proliferation and Heart Regeneration. Dev Cell 34, 387–399, doi:10.1016/j.devcel.2015.06.017 (2015).

67 Gabanyi, I. et al. Neuro-immune Interactions Drive Tissue Programming in Intestinal Macrophages. Cell 164, 378–391, doi:10.1016/j.cell.2015.12.023 (2016).

68 Lai, N. Y. et al. Gut-Innervating Nociceptor Neurons Regulate Peyer’s Patch Microfold Cells and SFB Levels to Mediate Salmonella Host Defense. Cell 180, 33–49 e22, doi:10.1016/j.cell.2019.11.014 (2020).

69 Matheis, F. et al. Adrenergic Signaling in Muscularis Macrophages Limits Infection-Induced Neuronal Loss. Cell 180, 64–78 e16, doi:10.1016/j.cell.2019.12.002 (2020).

70 Kenmoku, H., Ishikawa, H., Ote, M., Kuraishi, T. & Kurata, S. A subset of neurons controls the permeability of the peritrophic matrix and midgut structure in Drosophila adults. J Exp Biol 219, 2331–2339, doi:10.1242/jeb.122960 (2016).

71 Han, H. et al. Gut-neuron interaction via Hh signaling regulates intestinal progenitor cell differentiation in Drosophila. Cell Discov 1, 15006, doi:10.1038/celldisc.2015.6 (2015).

72 Cognigni, P., Bailey, A. P. & Miguel-Aliaga, I. Enteric neurons and systemic signals couple nutritional and reproductive status with intestinal homeostasis. Cell Metab 13, 92–104, doi:10.1016/j.cmet.2010.12.010 (2011).

73 Littleton, J. T., Bellen, H. J. & Perin, M. S. Expression of synaptotagmin in Drosophila reveals transport and localization of synaptic vesicles to the synapse. Development 118, 1077–1088, doi:10.1242/dev.118.4.1077 (1993).

74 McCorry, L. K. Physiology of the autonomic nervous system. Am J Pharm Educ 71, 78, doi:10.5688/aj710478 (2007).

75 Kim, A. J., Fenk, L. M., Lyu, C. & Maimon, G. Quantitative Predictions Orchestrate Visual Signaling in Drosophila. Cell 168, 280–294 e212, doi:10.1016/j.cell.2016.12.005 (2017).

76 Jenett, A. et al. A GAL4-driver line resource for Drosophila neurobiology. Cell Rep 2, 991–1001, doi:10.1016/j.celrep.2012.09.011 (2012).

77 Yagi, R., Mayer, F. & Basler, K. Refined LexA transactivators and their use in combination with the Drosophila Gal4 system. Proc Natl Acad Sci U S A 107, 16166–16171, doi:10.1073/pnas.1005957107 (2010).

78 Potter, C. J., Tasic, B., Russler, E. V., Liang, L. & Luo, L. The Q system: a repressible binary system for transgene expression, lineage tracing, and mosaic analysis. Cell 141, 536–548, doi:10.1016/j.cell.2010.02.025 (2010).

79 Kitamoto, T. Conditional modification of behavior in Drosophila by targeted expression of a temperature-sensitive shibire allele in defined neurons. J Neurobiol 47, 81–92, doi:10.1002/neu.1018 (2001).

80 Rosenzweig, M. et al. The Drosophila ortholog of vertebrate TRPA1 regulates thermotaxis. Genes Dev 19, 419–424, doi:10.1101/gad.1278205 (2005).

81 Inagaki, H. K. et al. Optogenetic control of Drosophila using a red-shifted channelrhodopsin reveals experience-dependent influences on courtship. Nat Methods 11, 325–332, doi:10.1038/nmeth.2765 (2014).

82 Matteoli, G. et al. A distinct vagal anti-inflammatory pathway modulates intestinal muscularis resident macrophages independent of the spleen. Gut 63, 938–948, doi:10.1136/gutjnl-2013-304676 (2014).

83 Jarret, A. et al. Enteric Nervous System-Derived IL-18 Orchestrates Mucosal Barrier Immunity. Cell 180, 813–814, doi:10.1016/j.cell.2020.02.004 (2020).

84 Huston, J. M. et al. Splenectomy inactivates the cholinergic antiinflammatory pathway during lethal endotoxemia and polymicrobial sepsis. J Exp Med 203, 1623–1628, doi:10.1084/jem.20052362 (2006).

85 Babcock, D. T., Landry, C. & Galko, M. J. Cytokine signaling mediates UV-induced nociceptive sensitization in Drosophila larvae. Curr Biol 19, 799–806, doi:10.1016/j.cub.2009.03.062 (2009).

86 Andersen, D. S. et al. The Drosophila TNF receptor Grindelwald couples loss of cell polarity and neoplastic growth. Nature 522, 482–486, doi:10.1038/nature14298 (2015).

87 Kanda, H., Igaki, T., Kanuka, H., Yagi, T. & Miura, M. Wengen, a member of the Drosophila tumor necrosis factor receptor superfamily, is required for Eiger signaling. J Biol Chem 277, 28372–28375, doi:10.1074/jbc.C200324200 (2002).

88 Guiza, J., Barria, I., Saez, J. C. & Vega, J. L. Innexins: Expression, Regulation, and Functions. Front Physiol 9, 1414, doi:10.3389/fphys.2018.01414 (2018).

89 Peters, M. A., Teramoto, T., White, J. Q., Iwasaki, K. & Jorgensen, E. M. A calcium wave mediated by gap junctions coordinates a rhythmic behavior in C. elegans. Curr Biol 17, 1601–1608, doi:10.1016/j.cub.2007.08.031 (2007).

90 Ho, K. Y. L., Khadilkar, R. J., Carr, R. L. & Tanentzapf, G. A gap-junction-mediated, calcium-signaling network controls blood progenitor fate decisions in hematopoiesis. Curr Biol 31, 4697–4712 e4696, doi:10.1016/j.cub.2021.08.027 (2021).

91 Dahl, G. & Muller, K. J. Innexin and pannexin channels and their signaling. FEBS Lett 588, 1396–1402, doi:10.1016/j.febslet.2014.03.007 (2014).

92 Bao, L. et al. Innexins form two types of channels. FEBS Lett 581, 5703–5708, doi:10.1016/j.febslet.2007.11.030 (2007).

93 Xie, Z. et al. The role of the Hippo pathway in the pathogenesis of inflammatory bowel disease. Cell Death Dis 12, 79, doi:10.1038/s41419-021-03395-3 (2021).

94 Mudter, J. & Neurath, M. F. Il-6 signaling in inflammatory bowel disease: pathophysiological role and clinical relevance. Inflamm Bowel Dis 13, 1016–1023, doi:10.1002/ibd.20148 (2007).

95 Tournier, J. M. et al. alpha3alpha5beta2-Nicotinic acetylcholine receptor contributes to the wound repair of the respiratory epithelium by modulating intracellular calcium in migrating cells. Am J Pathol 168, 55–68, doi:10.2353/ajpath.2006.050333 (2006).

96 Maouche, K. et al. {alpha}7 nicotinic acetylcholine receptor regulates airway epithelium differentiation by controlling basal cell proliferation. Am J Pathol 175, 1868–1882, doi:10.2353/ajpath.2009.090212 (2009).

97 Maouche, K. et al. Contribution of alpha7 nicotinic receptor to airway epithelium dysfunction under nicotine exposure. Proc Natl Acad Sci U S A 110, 4099–4104, doi:10.1073/pnas.1216939110 (2013).

98 Harris, M. P. Bioelectric signaling as a unique regulator of development and regeneration. Development 148, doi:10.1242/dev.180794 (2021).

99 Tyler, S. E. B. Nature’s Electric Potential: A Systematic Review of the Role of Bioelectricity in Wound Healing and Regenerative Processes in Animals, Humans, and Plants. Front Physiol 8, 627, doi:10.3389/fphys.2017.00627 (2017).

100 Margolis, K. G. & Gershon, M. D. Enteric Neuronal Regulation of Intestinal Inflammation. Trends Neurosci 39, 614–624, doi:10.1016/j.tins.2016.06.007 (2016).

101 Olofsson, P. S., Rosas-Ballina, M., Levine, Y. A. & Tracey, K. J. Rethinking inflammation: neural circuits in the regulation of immunity. Immunol Rev 248, 188–204, doi:10.1111/j.1600-065X.2012.01138.x (2012).

102 Shannon, E. K. et al. Multiple Mechanisms Drive Calcium Signal Dynamics around Laser-Induced Epithelial Wounds. Biophys J 113, 1623–1635, doi:10.1016/j.bpj.2017.07.022 (2017).

103 Levin, M. Bioelectric mechanisms in regeneration: Unique aspects and future perspectives. Semin Cell Dev Biol 20, 543–556, doi:10.1016/j.semcdb.2009.04.013 (2009).

104 Tseng, A. S., Beane, W. S., Lemire, J. M., Masi, A. & Levin, M. Induction of vertebrate regeneration by a transient sodium current. J Neurosci 30, 13192–13200, doi:10.1523/JNEUROSCI.3315-10.2010 (2010).

105 Gould, L. et al. Chronic wound repair and healing in older adults: current status and future research. J Am Geriatr Soc 63, 427–438, doi:10.1111/jgs.13332 (2015).

106 Kloth, L. C. Electrical Stimulation Technologies for Wound Healing. Adv Wound Care (New Rochelle) 3, 81–90, doi:10.1089/wound.2013.0459 (2014).

107 Howland, R. H. Vagus Nerve Stimulation. Curr Behav Neurosci Rep 1, 64–73, doi:10.1007/s40473-014-0010-5 (2014).

108 Kim, A. A. et al. Independently paced Ca2+ oscillations in progenitor and differentiated cells in an ex vivo epithelial organ. J Cell Sci 135, doi:10.1242/jcs.260249 (2022).

109 Fan, F. et al. Pharmacological targeting of kinases MST1 and MST2 augments tissue repair and regeneration. Sci Transl Med 8, 352ra108, doi:10.1126/scitranslmed.aaf2304 (2016).

110 Dey, A., Varelas, X. & Guan, K. L. Targeting the Hippo pathway in cancer, fibrosis, wound healing and regenerative medicine. Nat Rev Drug Discov 19, 480–494, doi:10.1038/s41573-020-0070-z (2020).

111 Gareb, B., Otten, A. T., Frijlink, H. W., Dijkstra, G. & Kosterink, J. G. W. Review: Local Tumor Necrosis Factor-alpha Inhibition in Inflammatory Bowel Disease. Pharmaceutics 12, doi:10.3390/pharmaceutics12060539 (2020).

112 D’Haens, G. R. & van Deventer, S. 25 years of anti-TNF treatment for inflammatory bowel disease: lessons from the past and a look to the future. Gut 70, 1396–1405, doi:10.1136/gutjnl-2019-320022 (2021).

113 Payne, S. C. et al. Anti-inflammatory Effects of Abdominal Vagus Nerve Stimulation on Experimental Intestinal Inflammation. Front Neurosci 13, 418, doi:10.3389/fnins.2019.00418 (2019).

114 Bonaz, B., Sinniger, V. & Pellissier, S. Therapeutic Potential of Vagus Nerve Stimulation for Inflammatory Bowel Diseases. Front Neurosci 15, 650971, doi:10.3389/fnins.2021.650971 (2021).

115 Phillips, M. D. & Thomas, G. H. Brush border spectrin is required for early endosome recycling in Drosophila. J Cell Sci 119, 1361–1370, doi:10.1242/jcs.02839 (2006).

116 He, L., Si, G., Huang, J., Samuel, A. D. T. & Perrimon, N. Mechanical regulation of stem-cell differentiation by the stretch-activated Piezo channel. Nature 555, 103–106, doi:10.1038/nature25744 (2018).

117 Diao, F. et al. Plug-and-play genetic access to drosophila cell types using exchangeable exon cassettes. Cell Rep 10, 1410–1421, doi:10.1016/j.celrep.2015.01.059 (2015).

118 Perkins, L. A. et al. The Transgenic RNAi Project at Harvard Medical School: Resources and Validation. Genetics 201, 843–852, doi:10.1534/genetics.115.180208 (2015).

119 Chakraborty, S. et al. Mutant IP3 receptors attenuate store-operated Ca2+ entry by destabilizing STIM-Orai interactions in Drosophila neurons. J Cell Sci 129, 3903–3910, doi:10.1242/jcs.191585 (2016).

120 Nern, A., Pfeiffer, B. D. & Rubin, G. M. Optimized tools for multicolor stochastic labeling reveal diverse stereotyped cell arrangements in the fly visual system. Proc Natl Acad Sci U S A 112, E2967–2976, doi:10.1073/pnas.1506763112 (2015).

121 Hamada, F. N. et al. An internal thermal sensor controlling temperature preference in Drosophila. Nature 454, 217–220, doi:10.1038/nature07001 (2008).

122 Ni, J. Q. et al. A genome-scale shRNA resource for transgenic RNAi in Drosophila. Nat Methods 8, 405–407, doi:10.1038/nmeth.1592 (2011).

123 Riabinina, O. et al. Improved and expanded Q-system reagents for genetic manipulations. Nat Methods 12, 219–222, 215 p following 222, doi:10.1038/nmeth.3250 (2015).

124 Gratz, S. J., Harrison, M. M., Wildonger, J. & O’Connor-Giles, K. M. Precise Genome Editing of Drosophila with CRISPR RNA-Guided Cas9. Methods Mol Biol 1311, 335–348, doi:10.1007/978-1-4939-2687-9_22 (2015).

125 Port, F., Chen, H. M., Lee, T. & Bullock, S. L. Optimized CRISPR/Cas tools for efficient germline and somatic genome engineering in Drosophila. Proc Natl Acad Sci U S A 111, E2967–2976, doi:10.1073/pnas.1405500111 (2014).

126 Kim, K. et al. Drosophila as a model for studying cystic fibrosis pathophysiology of the gastrointestinal system. Proc Natl Acad Sci U S A 117, 10357–10367, doi:10.1073/pnas.1913127117 (2020).

127 Li, H. et al. Fly Cell Atlas: A single-nucleus transcriptomic atlas of the adult fruit fly. Science 375, eabk2432, doi:10.1126/science.abk2432 (2022).

128 Buchon, N., Broderick, N. A., Poidevin, M., Pradervand, S. & Lemaitre, B. Drosophila intestinal response to bacterial infection: activation of host defense and stem cell proliferation. Cell Host Microbe 5, 200–211, doi:10.1016/j.chom.2009.01.003 (2009).

129 Chen, J., Xu, N., Huang, H., Cai, T. & Xi, R. A feedback amplification loop between stem cells and their progeny promotes tissue regeneration and tumorigenesis. Elife 5, doi:10.7554/eLife.14330 (2016).

